# Complete genome sequence of *Comamonas testosteroni* TA441 and comparative analysis of steroid degradation gene clusters across Proteobacteria and Actinomycetota

**DOI:** 10.64898/2026.09.18.752586

**Authors:** Masae Horinouchi, Nozomi Kobayashi-Yoshida, Hideaki Nojiri, Hayashi Toshiaki

## Abstract

*Comamonas testosteroni* TA441 is a model aerobic steroid-degrading bacterium whose sterane degradation pathway has been elucidated in greater detail than that of any other bacterium to date. Similar pathways have been identified in many bacterial genera, including members of Proteobacteria and Actinomycetota, such as *Mycobacterium tuberculosis*. Steroid degradation genes in TA441 form an approximately 120-kb mega-cluster, and 40 genes involved mainly in the degradation of the sterane structure and the C20–C22 portion of the side chain at C17 have been identified. Here, we present the complete genome sequence of TA441 together with our findings on its steroid degradation genes. We also compare the genes in the mega-cluster with corresponding genes in other *Comamonas* strains, β-Proteobacteria other than *Comamonas*, γ-Proteobacteria, α-Proteobacteria, and Actinomycetota using CAGECAT to analyze similarities and differences in steroid degradation genes among these bacteria.

**IMPORTANCE:** *Comamonas testosteroni* TA441 is a key model organism for studying bacterial aerobic steroid degradation, and its pathways for cleavage of the A-, B-, C-, and D-rings have been extensively studied. These pathways and their enzymes show striking similarities among bacteria, including *Mycobacterium tuberculosis*, in which cholesterol degradation contributes to persistence in chronically infected lungs. However, the ecological and physiological significance of bacterial steroid degradation remains incompletely understood. Although TA441 is one of the best-studied steroid-degrading bacteria, its complete genome sequence had not been determined. In addition, TA441 and *Comamonas thiooxydans* CNB-2 have sometimes been described as nearly identical strains, although CNB-2 has not been studied for steroid degradation since its genome was published in 2009. Here, we present the complete genome sequence of TA441 and a detailed, gene-by-gene comparison of its steroid degradation genes with those of other bacteria, including *M. tuberculosis*. This study provides a foundation for understanding the diversity and potential ecological and physiological significance of bacterial steroid degradation.

## INTRODUCTION

*Comamonas testosteroni* TA441 is a model strain for studying aerobic steroid degradation, whose 120-kb mega-cluster of steroid degradation genes and the genes responsible for degradation of the sterane structure (the four-ring steroid nucleus) were revealed in detail for the first time among bacteria. Aerobic degradation of steroids by bacteria has been studied for over 70 years, with pioneering studies in the 1950s and 1960s using two representative steroid-degrading bacteria, the actinomycete *Rhodococcus equi* (formerly *Nocardia restrictus*) and the proteobacterium *Comamonas testosteroni* (formerly *Pseudomonas testosteroni*), for the purpose of obtaining materials for steroid drugs (1–4). These studies established that bacterial aerobic steroid degradation starts with B-ring cleavage accompanied by aromatization of the A-ring, followed by cleavage and removal of the aromatized A-ring.

Studies on steroid degradation genes began in the 1990s. Enzymes involved in B-ring cleavage accompanied by aromatization of the A-ring, such as 3α-hydroxysteroid dehydrogenase (3α-DH) (*C. testosteroni*) (5–7), 3β-dehydrogenase (3β-DH) (*Comamonas* strains) (8), Δ1-dehydrogenase (Δ1-DH) (*N. corallina* (9) and *R. rhodochrous*(10)), Δ4-5 isomerase (ksi) (*C. testosteroni*(11), *P. putida*(12), and *Arthrobacter simplex* (13)), and 9α-hydroxylase (KshAB, KshA2) (*R. erythropolis* (14)) (see Supplementary Materials, Fig. S1), were identified before we identified the aromatized A-ring cleavage enzyme TesB.

TesB was identified in *C. testosteroni* TA441 in 2001 as the first enzyme involved in aromatized A-ring cleavage of the sterane structure to be identified among steroid-degrading bacteria (15). The entire A- and B-ring cleavage pathway and the genes involved in the production of 9,17-dioxo-1,2,3,4,10,19-hexanorandrostan-5-oic acid (9,17-DOHNA, also known as HIP), an important intermediate containing the C- and D-rings after B-ring cleavage, in *C. testosteroni* TA441 were identified and reported between 2001 and 2006 (16–22). In Actinomycetota, a similar A- and B-ring cleavage process was reported in *R. jostii* RHA1 in 2007 (23) and *M. tuberculosis* in 2009 (24). In TA441, 9,17-DOHNA is converted to its CoA ester by ScdA (21), initiating B-, C-, and D-ring degradation, which proceeds mainly through several cycles of β-oxidation (25–35) (the abbreviated degradation pathway is shown in Fig. 1 and the detailed degradation pathway elucidated in TA441 is shown in Supplementary Materials, Fig. S1). A similar but not identical B-, C-, and D-ring degradation pathway was reported in *R. jostii* RHA1 and *M. tuberculosis* (36). Although this pathway is relatively incomplete in these bacteria, these findings imply that B-, C-, and D-ring degradation processes are also similar between Proteobacteria and Actinomycetota. These steroid degradation genes constitute the 120-kb mega-cluster in *C. testosteroni* TA441, which was also identified for the first time among bacteria (34, 37).

**Fig 1.**
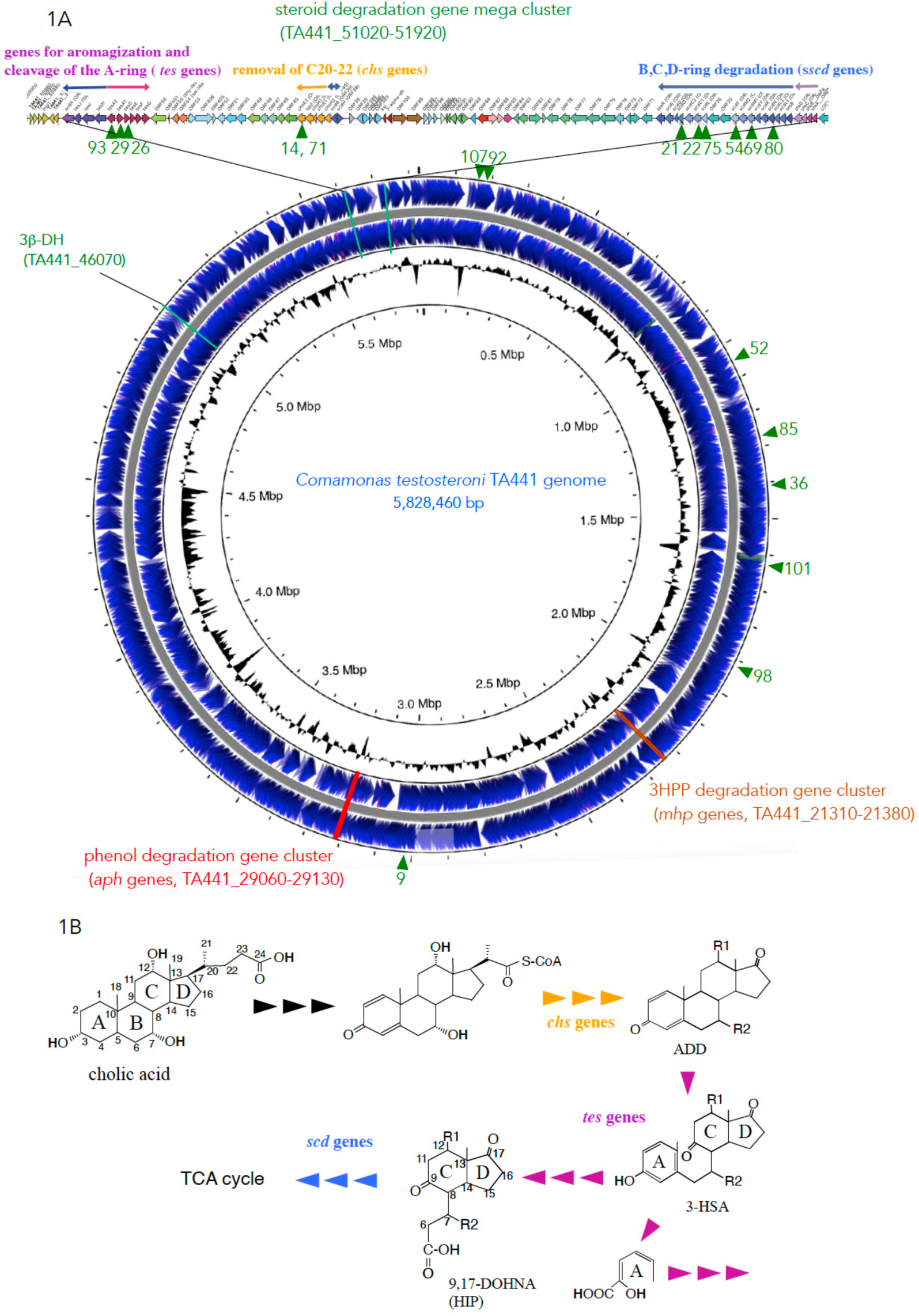
Circular chromosome map of the complete genome of *C. testosteroni* TA441 generated using the Proksee tool (https://proksee.ca). Genes above the circular map indicate the 120-kb steroid degradation mega-cluster in TA441, which contains three gene clusters involved in A-ring aromatization and subsequent A- and B-ring cleavage (*tes* genes), C20–22 side-chain removal (*chs* genes), and B-, C-, and D-ring degradation (*scd* genes). Abbreviated steroid degradation pathway is shown below the map (the whole degradation pathway is in the Supplementary Materials Fig. S1). Green arrowheads indicate insertion of transposon in the mutants with lower growth on steroids generated in the previous study (17). The precise locations of the transposon insertion are listed in Table 2. Two aromatic compound degradation gene clusters in TA441, the *aph* genes for phenol degradation and the *mhp* genes for 3-(3-hydroxyphenyl)propionic acid (3HPP) degradation, are indicated in red and in dark pink.

**Table 1.** Genome features of *Comamonas testosteroni* TA441.

| Feature | Value |
| --- | --- |
| Genome size | 5,828,460 bp |
| Replicons | 1 chromosome |
| GC content | 61.2 % |
| Total genes | 5,395 |
| Coding sequences (CDSs) | 5,273 |
| rRNA genes | 21 |
| tRNA genes | 101 |
| rRNA operons | 7 |
| Coding ratio | 87.0 % |
| Average protein length | 320.7 aa |
| CRISPR loci | Not detected |

**Table 2.**
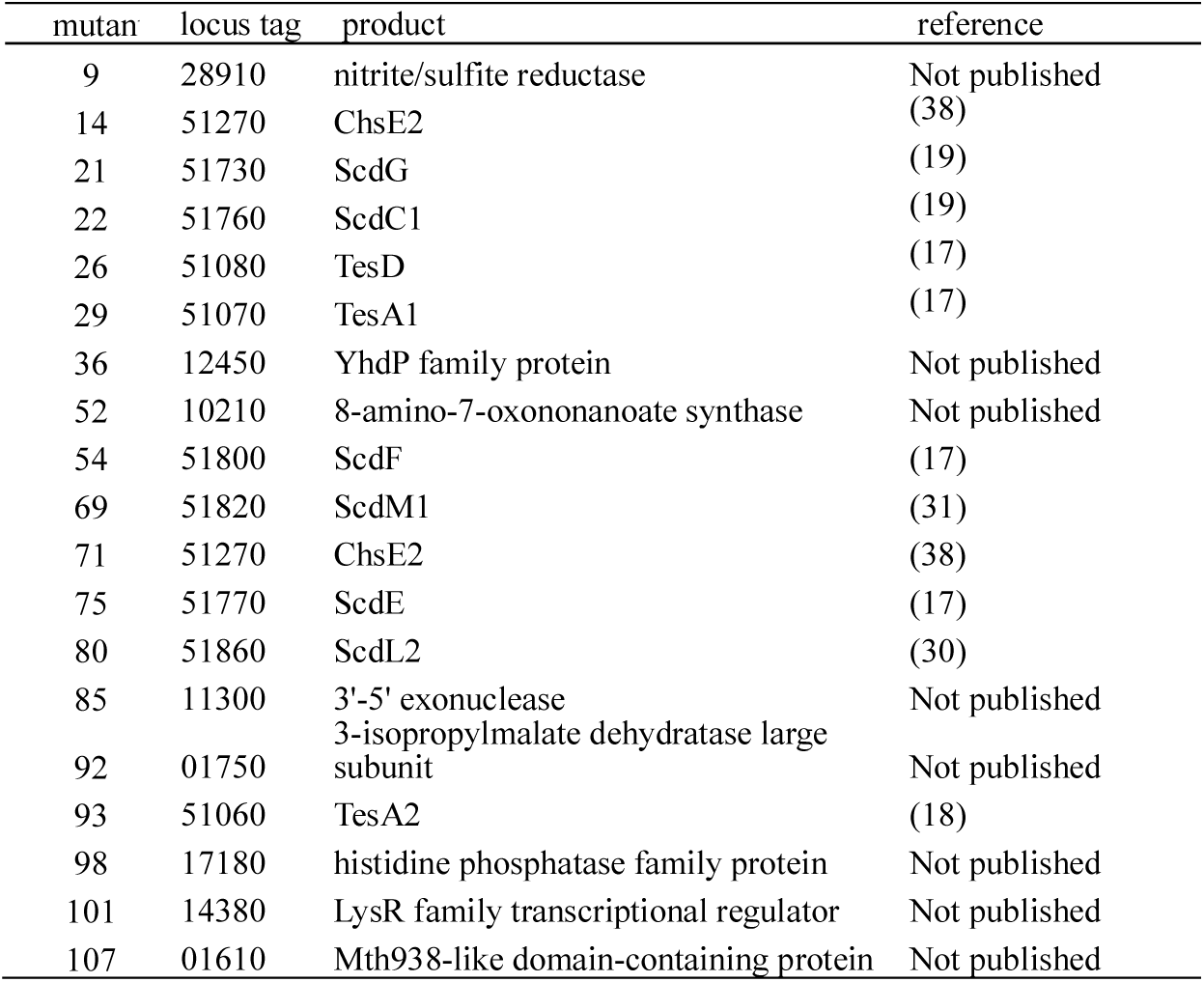
Transposon mutants of TA441.

However, despite its importance as a model organism for aerobic bacterial steroid degradation, the complete genome sequence of TA441 had not yet been determined. Here, we describe the complete genome sequence of TA441 and the distribution of aerobic bacterial steroid degradation genes similar to those of TA441 among bacteria.

## RESULTS AND DISCUSSION

### General genome features

The complete genome sequence of *C. testosteroni* TA441 consists of a single circular chromosome of 5,828,460 bp with a GC content of 61.2% (Fig. 1). Genome annotation predicted 5,273 coding sequences (CDSs), 101 tRNA genes, and 21 rRNA genes corresponding to seven rRNA operons. General features of the genome are summarized in Table 1.

### Steroid degradation gene cluster and two aromatic compound degradation gene clusters

TA441 is a representative aerobic steroid-degrading proteobacterium in which the degradation pathway of the sterane structure has been elucidated in detail, the most comprehensively among bacteria (33, 37). The sterane degradation genes and other genes, such as those involved in C17 side-chain degradation, form an approximately 120-kb mega-cluster of steroid degradation genes. Steroid degradation begins with removal of the C17 side chain (*chs* genes for removal of C20–C22 and the nearby genes yet to be identified, Fig. 1A) (38), followed by aromatization of the A-ring accompanied by cleavage of the B-ring and cleavage of the aromatized A-ring (*tes* genes) (16–22).

The remaining C- and D-rings and the cleaved B-ring are then degraded mainly by β-oxidation (*scd* genes) (25–35) (an abbreviated degradation pathway is shown in Fig. 1B, and the detailed degradation pathway is shown in Supplementary Materials, Fig. S1).

The *chs*, *tes*, and *scd* genes each form a cluster; the *tes* and *scd* clusters are located at both ends of the mega-cluster, whereas the *chs* cluster is located in the DNA region between them. The locus of the mega-cluster on the TA441 genome is TA441_51020 to TA441_51920 (Fig. 1A). Among the previously characterized steroid degradation genes in TA441, only the 3β-dehydrogenase (3β-DH) gene is located approximately 600 kb away from the mega-cluster (locus TA441_51020).

TA441 has at least two aromatic compound degradation gene clusters: phenol degradation genes (*aph* genes) (39) and 3-(3-hydroxyphenyl)propionic acid (3HPP) (*mhp* genes) (40). The steroid degradation gene cluster and these two clusters are located separately on the TA441 genome (*aph*, TA441_29060 to 29130; *mhp*, TA441_21310 to 21380).

We identified the steroid degradation genes using transposon mutants that showed little growth on cholic acid and/or testosterone but grew well with *p*-hydroxybenzoate (a positive-control substrate used instead of a commonly used substrate such as citrate) (16). The location of the transposon insertion in each mutant is summarized in Table 1 and indicated by green arrowheads in Fig. 1. Eight mutants had transposon insertions outside the mega-cluster. These mutants have not yet been analyzed, but the locations of the disrupted genes did not indicate the presence of other steroid degradation gene clusters, except for those in mutants 92 and 107, which were relatively close to the mega-cluster and the disrupted genes encode a 3-isopropylmalate dehydratase-like protein and an Mth938-like domain-containing protein (adipogenesis associated Mth938 domain containing protein is involved in lipid metabolism).

### Distribution of the steroid degradation gene mega-cluster among *Comamonas* species

We then compared the TA441 genome sequence with those of other *Comamonas* strains for which complete genome sequences are available. We basically selected one strain for each species, choosing the strain whose steroid degradation gene(s) showed the highest identity to the corresponding genes in TA441. Mauve analysis indicated that *C. thiooxydans* CNB-2 (41), *C. testosteroni* DSM14576 (KF1) (42), a well-known polychlorinated biphenyl degrader *C. testosteroni* TK102 (43), *C. fluminis* CJ34 (44), and *C. resistens* ZM22 (45) have steroid degradation gene clusters similar to that of TA441 (indicated by red boxes in Fig. 2; the results for all tested *Comamonas* and related species are shown in Supplementary Materials, Fig. S2). *C. testosteroni* strain ATCC11996, the type strain whose steroid degradation has been studied since the 1950s, has steroid degradation genes highly similar to those of TA441 (46–49), but its genome sequence is available in 63 contigs, and we were therefore unable to compare it with the TA441 genome (50).

**Fig. 2.**
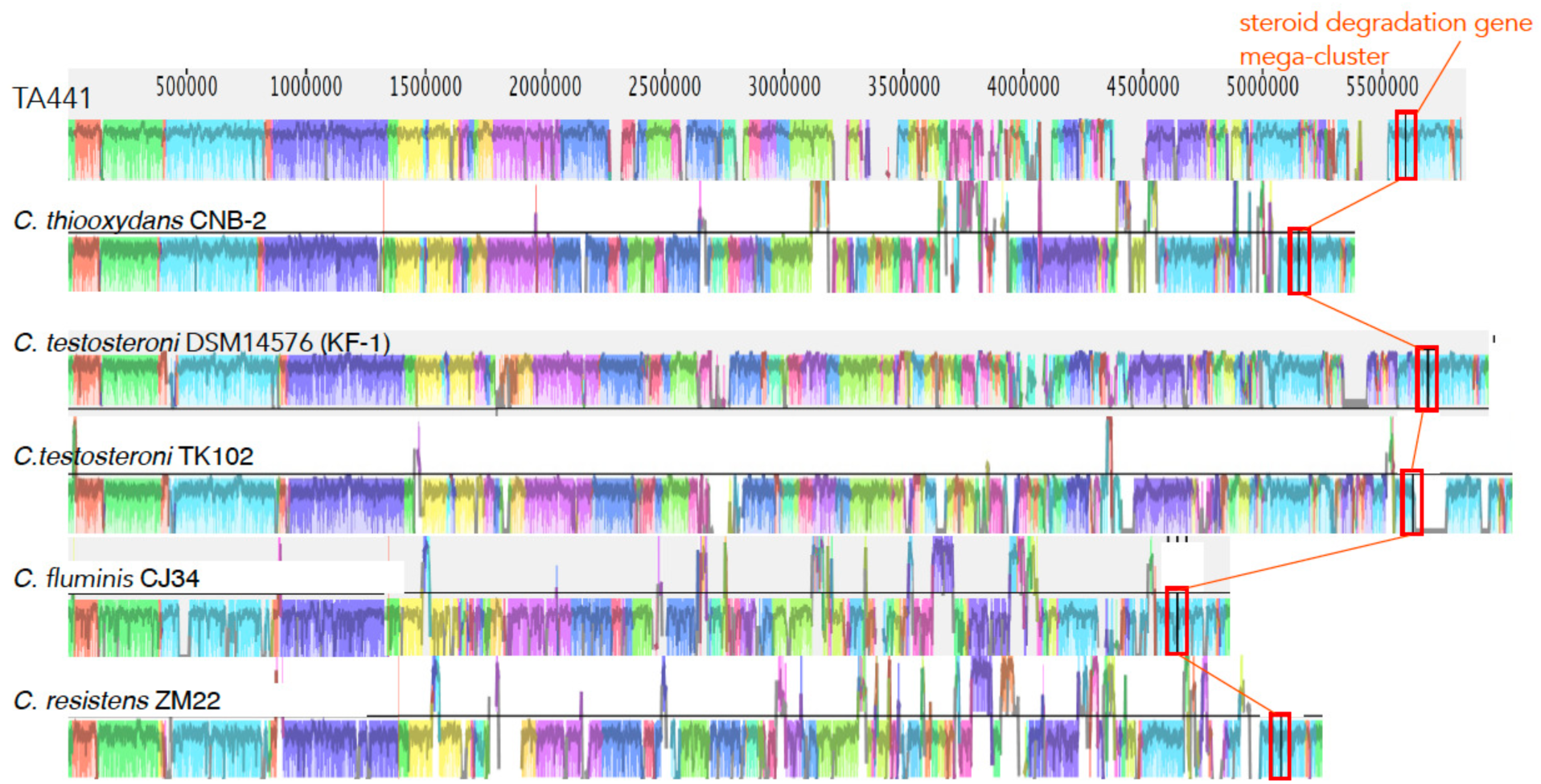
Mauve alignment of *Comamonas* strain genomes with the steroid degradation mega-cluster (indicated with red boxes). Only the genomes indicated to have putative steroid degradation gene mega-clusters similar to that of TA441 are shown: *C. thiooxydans* CNB-2 (41), *C. testosteroni* DSM14576 (KF1) (42), *C. testosteroni* TK102 (43), *C. fluminis* CJ34 (44), and *C. resistens* ZM22 (45) (the results for all tested strains are shown in Supplementary Fig. S2).

A phylogenetic tree based on 16S rRNA also indicated that TA441 and these species (and probably *C. brasiliensis*, for which only a shotgun sequence is available) are genetically closely related (four spices described above are indicated by pink boxes in Fig. 3A. The tree was generated directly using the BLAST Tree View based on BLASTn searches, and the strains included in the tree were not necessarily the same as those used in the Mauve analysis).

**Fig. 3.**
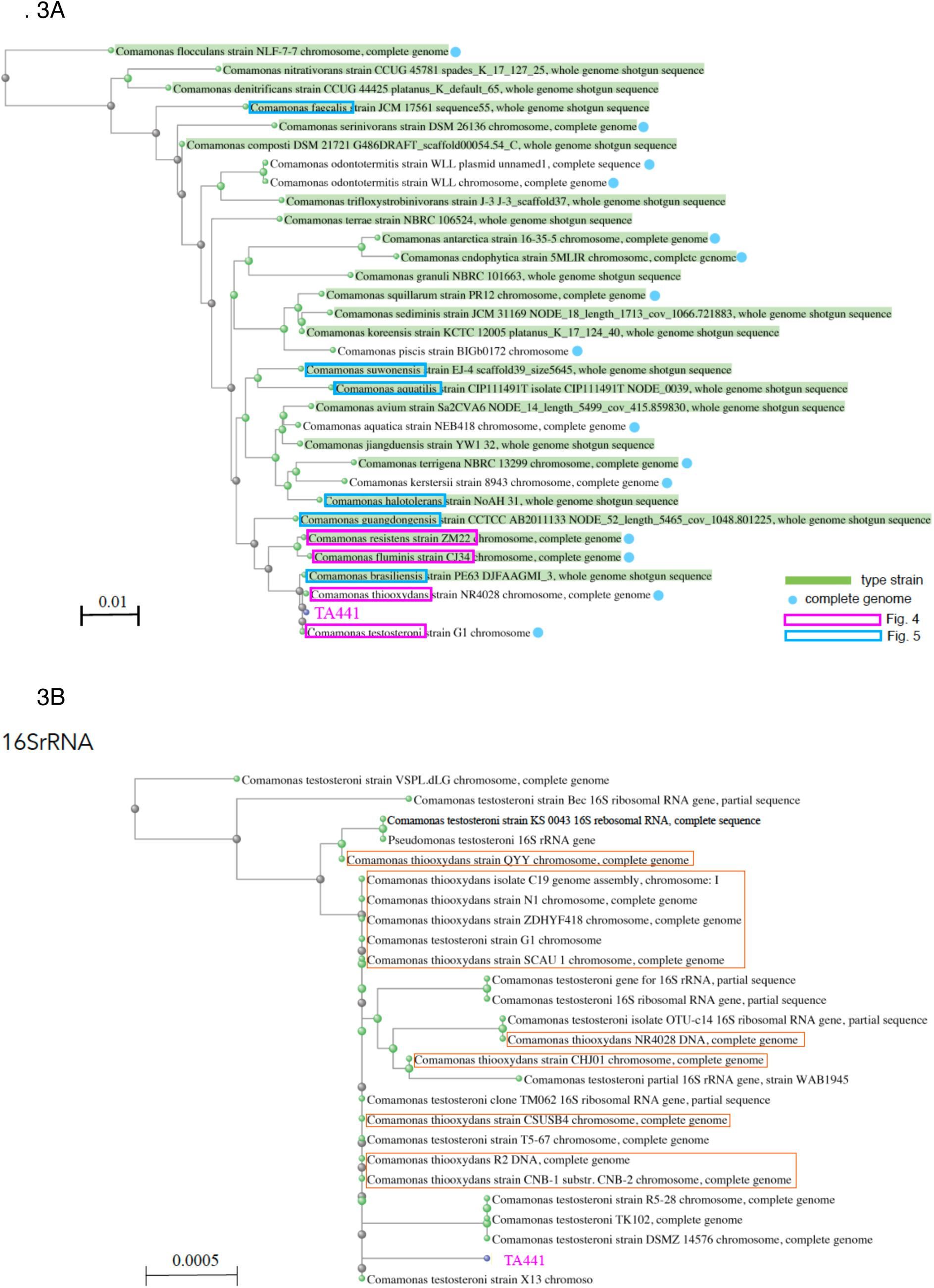
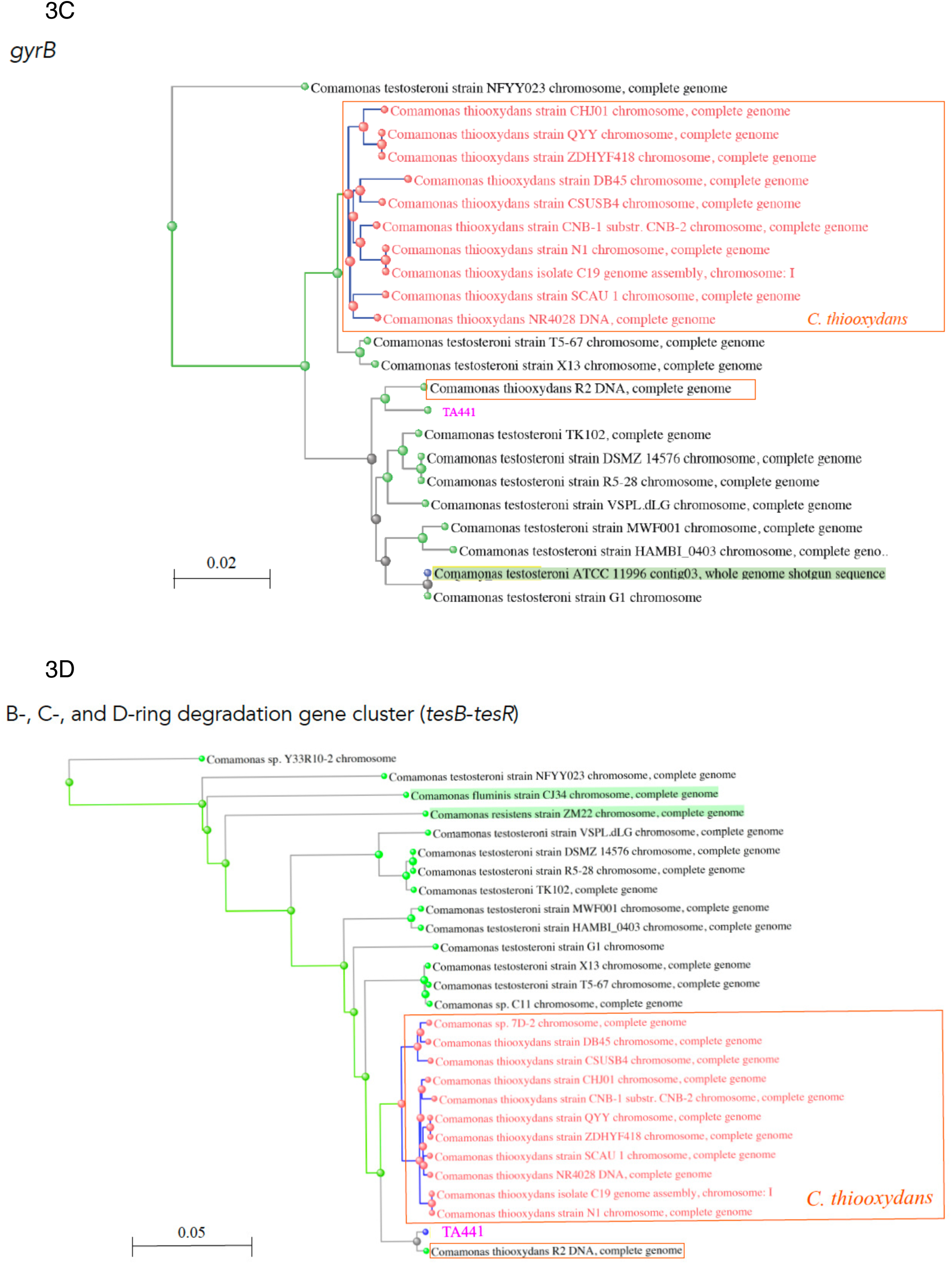
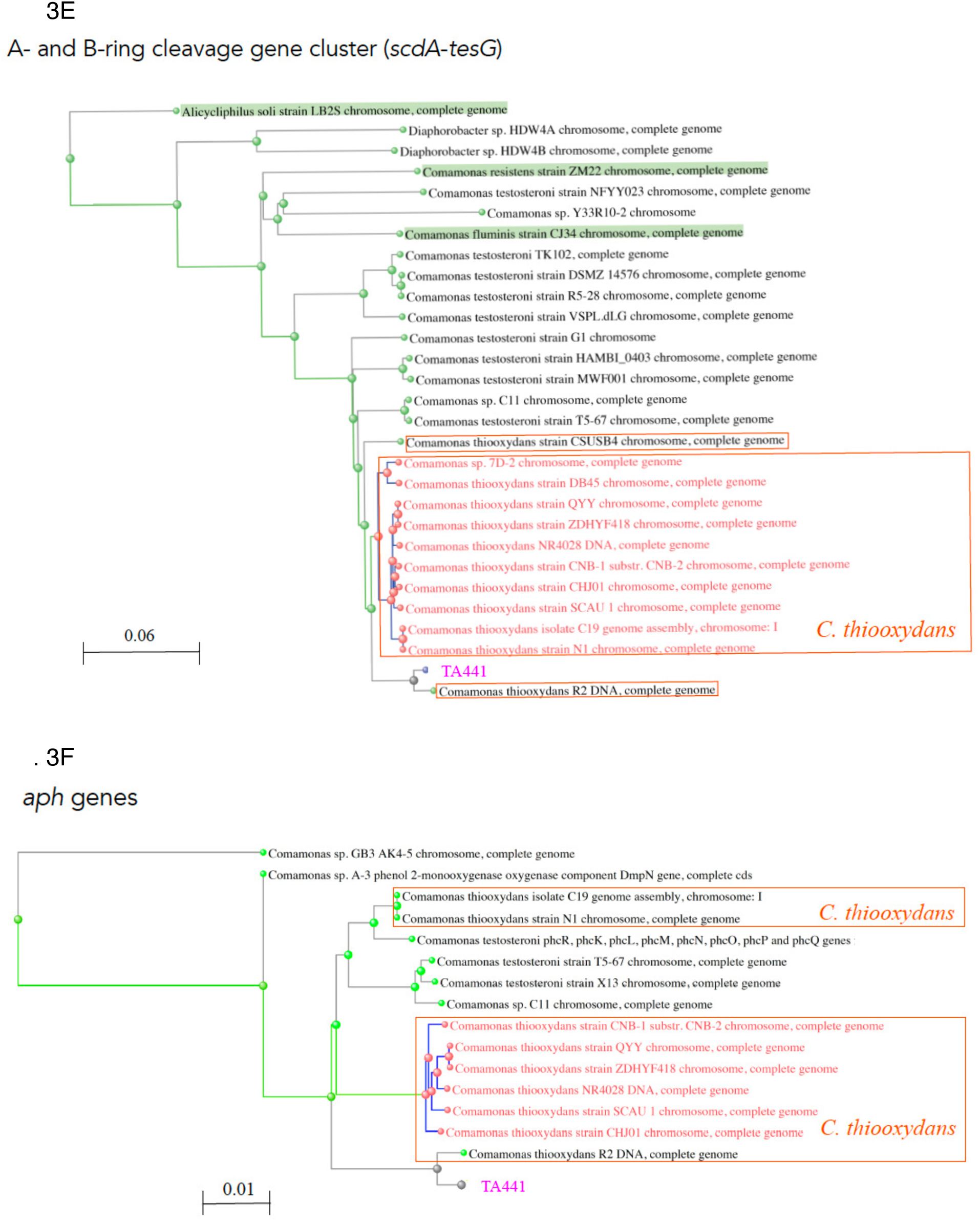
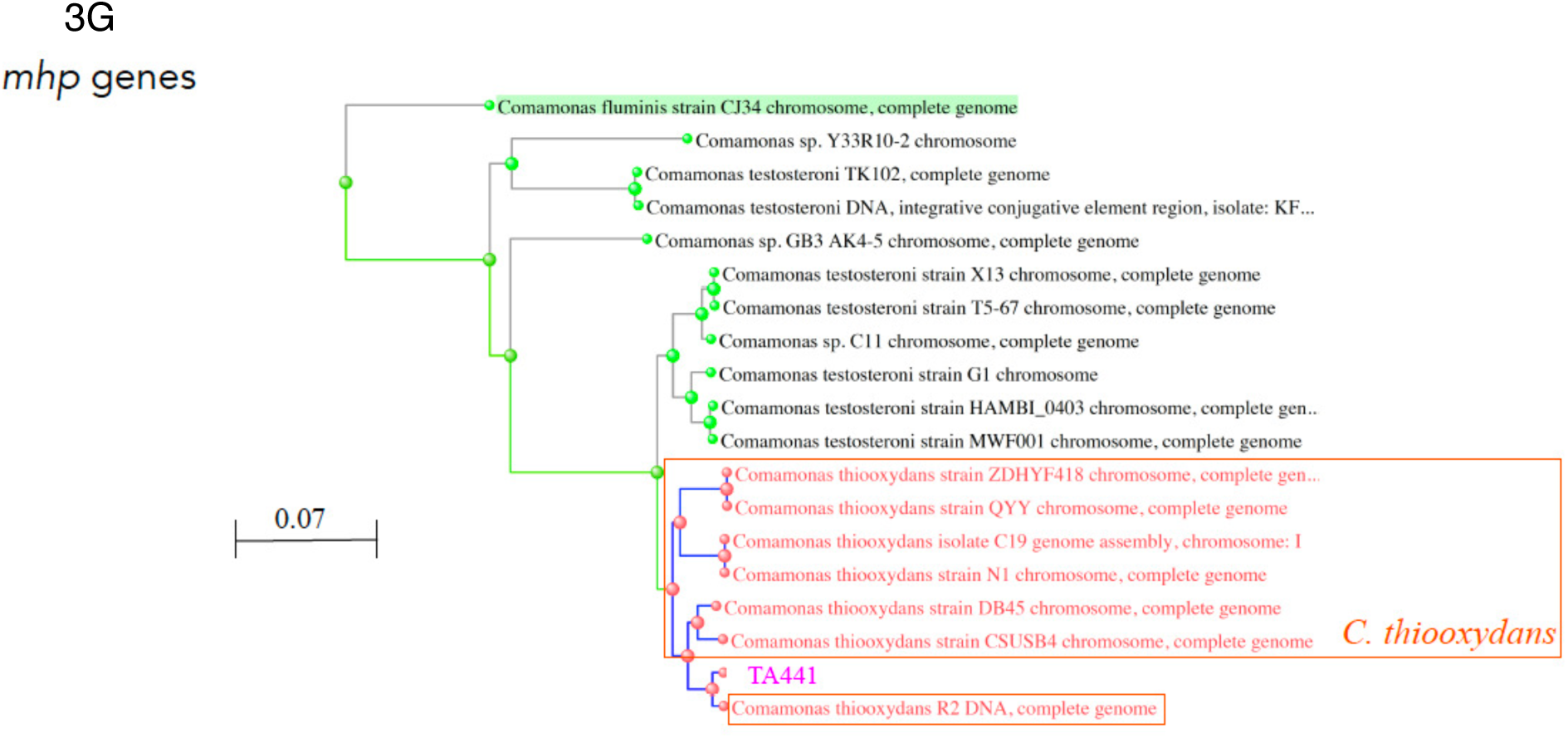
**3A** Phylogenetic tree based on 16S rRNA gene sequences of TA441 and *Comamonas* strains. The tree was generated from the resulting BLAST sequence sets using the Tree View function of BLAST. Pink box: strains with a putative steroid degradation mega-cluster suggested by Mauve analysis (presented in Fig. 4). Light blue box: other *Comamonas* strains with a putative steroid degradation mega-cluster (presented in Fig. 5). Light blue circle indicates the strain with a complete genome, and light green background indicates type strains. **3B–G**: Phylogenetic trees based on 16S rRNA gene (**B**), gyrase B gene (**C**), *tesB* to *tesR* (**D**), *scdA* to *tesG* (**E**), *aph* genes for phenol degradation (**F**), and *mhp* genes for 3-(3-hydroxyphenyl)propionic acid (3HPP) degradation (**G**) in TA441 and corresponding sequences in *C. testosteroni* and *C. thiooxydans* strains. The trees were generated from the resulting BLAST sequence sets using the Tree View function of BLAST. Red boxes indicate *C. thiooxydans*.

### C. testosteroni and C. thiooxydans

*C. thiooxydans* was separated from *C. testosteroni* as a distinct species approximately fifteen years ago (51), but the high sequence identities of their 16S rRNA genes make it difficult to distinguish the two species (Fig. 3B; *C. thiooxydans* strains are indicated by red boxes). In contrast, the two species can be distinguished based on sequence analysis of the housekeeping gene *gyrB* (52). Phylogenetic analysis of *gyrB* sequences from various *C. testosteroni* and *C. thiooxydans* strains showed that *C. thiooxydans* strains except for strain R2 (53) form a cluster separate from the *C. testosteroni* strains (Fig. 3C; indicated by a large red box).

We then conducted BLAST similarity searches using the DNA sequences of the *scd* genes (*tesB* to *tesR*) and the *tes* genes (*scdA* to *tesG*), respectively (see Fig. 1), and directly generated phylogenetic trees from the BLAST results (Fig. 3D and 3E; *C. thiooxydans* strains are indicated by red boxes). The resulting two trees were quite similar to each other and also similar to the phylogenetic tree based on *gyrB* (Fig. 3C), with *C. thiooxydans* strains forming a separate cluster.

For comparison, we also generated phylogenetic trees based on the *aph* and *mhp* genes (Fig. 3F and 3G). In these trees, *C. testosteroni* and *C. thiooxydans* strains basically formed separate clusters, although the *C. thiooxydans* strains formed two clusters in the tree based on the *aph* genes. In the phylogenetic trees based on *gyrB* and the steroid degradation genes, TA441 and *C. thiooxydans* R2 (53) were located on the same branch and somewhat separated from the other *C. testosteroni*. The same association was also observed in the trees based on the *aph* and *mhp* genes.

### Comparison of the steroid degradation gene clusters I: *Comamonas* strains Mauve analysis indicated the presence of similar mega cluster of steroid degradation gene to that of TA441 (Fig. 4, identities are in table 3)

Mauve analysis indicated the presence of a steroid degradation gene mega-cluster similar to that of TA441 in *C. thiooxydans* CNB-2 (41), *C. testosteroni* DSM14576 (42), *C.testosteroni* TK102 (43), *C. fluminis* CJ34 (44), and *C. resistens* ZM22 (45). A BLAST analysis using the DNA sequences from *tesB* to *tesR* showed high similarity to DNA regions in these strains. Then approximately 200-kb DNA regions containing the putative *tesB* to *tesR*, the region extended approximately 50 kb upstream from the beginning of *tesB* and 150 kb downstream from the end of *tesR*, in these strains were subjected to comparative gene cluster analysis using CAGECAT, a tool for rapid search and visualization of homologous gene clusters (54). The identities between corresponding genes in these strains were high, and the cluster structures, including the locations of corresponding genes, were also highly similar, particularly for the *tes*, *scd*, and *chs* genes (Fig. 4; identities between the corresponding enzymes and those of TA441 are listed in Table 3). The structures of the TA441 mega-cluster and those of *C. thiooxydans* CNB-2’s (41), and that of *C. testosteroni* DSM14576’s (42) and *C. testosteroni* TK102’s(43) were almost identical. *C. fluminis* CF34 (44) also had a highly similar mega-cluster, except for a few genes. These results suggest that these strains may have the capacity to degrade cholic acid. In contrast, *C. resistens* ZM22 (45) lacks most of the genes located between the *tes* and *chs* genes and between the *chs* and *scd* genes, including genes necessary for cholic acid degradation.

**Fig. 4.**
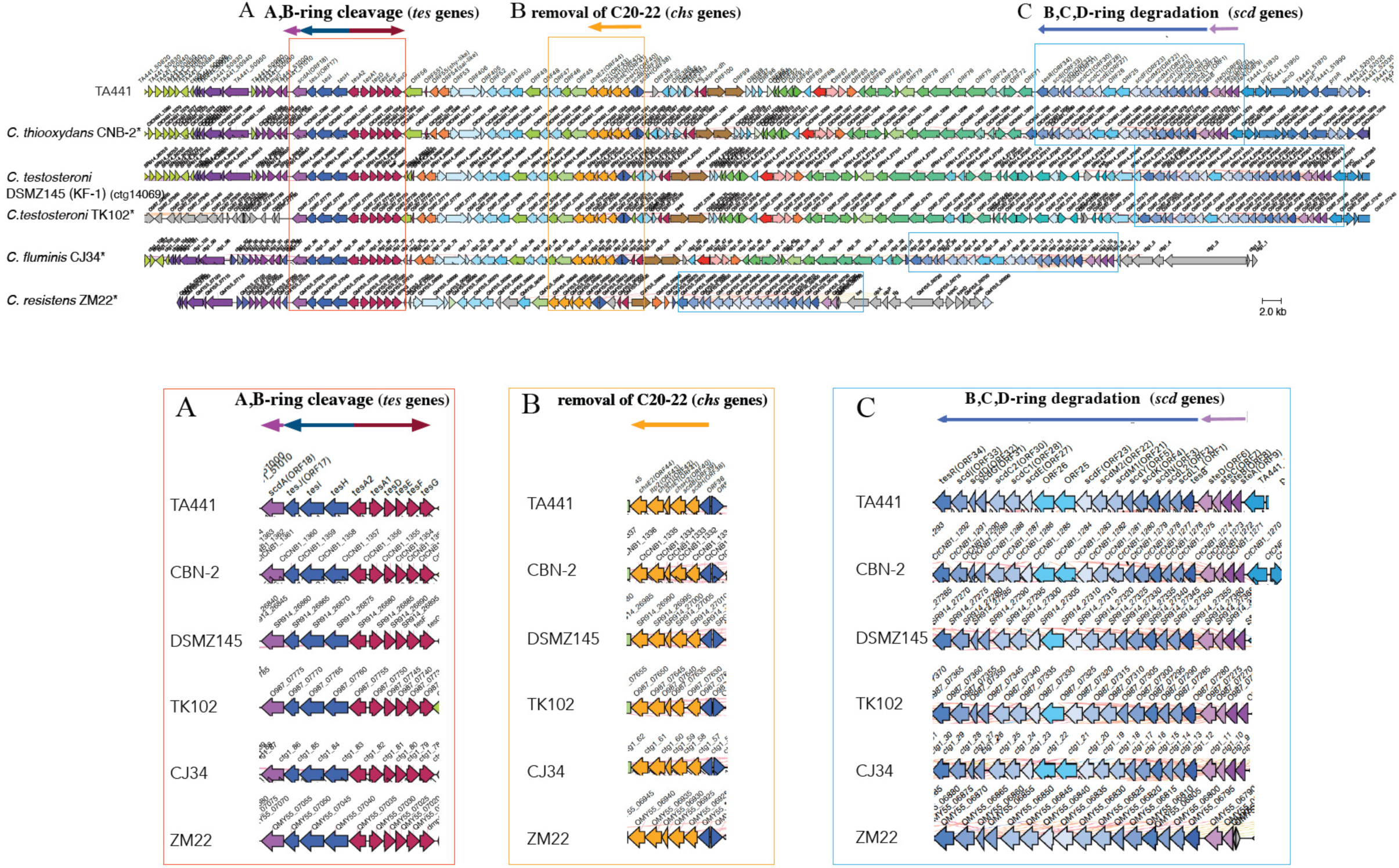
Comparison of the steroid degradation gene clusters of *Comamonas* strains suggested to have a putative steroid degradation mega-cluster similar to that of TA441 by Mauve analysis using CAGECAT. *: strains with complete genome sequences. The genes involved in A- and B-ring cleavage (*tes* genes) are shown in dark blue (*tesHIJ*) and dark red (*tesA1A2DEFG*); the CoA transferase acting after A- and B-ring cleavage (*scdA*) in pink; genes involved in B-, C-, and D-ring degradation (*scd* genes) in light blue; and *chs* genes in orange. The colors of genes outside the boxes do not necessarily indicate their functions and are used to indicate similar genes among the strains. Identities between TA441 steroid degradation enzymes and the corresponding enzymes in other strains are listed in Table 3.

**Table 3.** Amino acid identity (%) between the enzymes in Fig. 4 (locus tag or gene name) and corresponding enzymes in TA441.

| TA441 | TesB | ScdL1 | ScdL2 | ScdN | ScdK | ScdY | ScdM1 | ScdM2 | ScdF | ScdE | ScdC1 | ScdC2 | ScdG | ScdD | ScdJ | TesR |
| --- | --- | --- | --- | --- | --- | --- | --- | --- | --- | --- | --- | --- | --- | --- | --- | --- |
| CNB-2 | CtCNB1_1275 | CtCNB1_1276 | CtCNB1_1277 | CtCNB1_1278 | CtCNB1_1279 | CtCNB1_1280 | CtCNB1_1281 | CtCNB1_1282 | CtCNB1_1283 | CtCNB1_1286 | CtCNB1_12787 | CtCNB1_1288 | CtCNB1_1289 | CtCNB1_1290 | CtCNB1_1291 | CtCNB1_1292 |
|  | 93 | 99 | 97 | 98 | 99 | 97 | 99 | 99 | 98 | 92 | 97 | 100 | 99 | 97 | 94 | 100 |
| DSMZ145 | SR914_27350 | SR914_27345 | SR914_27340 | SR914_27335 | SR914_27330 | SR914_27325 | SR914_27320 | SR914_27315 | SR914_27310 | SR914_27300 | SR914_27295 | SR914_27290 | SR914_27285 | SR914_27280 | SR914_27275 | SR914_27270 |
|  | 96 | 98 | 96 | 96 | 97 | 96 | 97 | 94 | 95 | 96 | 90 | 93 | 96 | 93 | 97 | 99 |
| TK102 | O987_07285 | O987_07290 | O987_07295 | O987_07300 | O987_07305 | O987_07310 | O987_07315 | O987_07320 | O987_07325 | O987_07335 | O987_07340 | O987_07345 | O987_07350 | O987_07355 | O987_07360 | O987_07365 |
|  | 96 | 98 | 97 | 96 | 97 | 96 | 97 | 94 | 96 | 97 | 89 | 93 | 96 | 93 | 91 | 99 |
| CJ34 | ctg1_12 | ctg1_13 | ctg1_14 | ctg1_15 | ctg1_16 | ctg1_17 | ctg1_18 | ctg1_19 | ctg1_20 | ctg1_23 | ctg1_24 | ctg1_25 | ctg1_26 | ctg1_27 | ctg1_28 | ctg1_29 |
|  | 92 | 95 | 93 | 95 | 93 | 95 | 93 | 81 | 95 | 96 | 84 | 89 | 97 | 86 | 97 | 95 |
| ZM22 | QMY55_06795 | QMY55_06800 | QMY55_06805 | QMY55_06810 | QMY55_06815 | QMY55_06820 | QMY55_06825 | QMY55_06830 | QMY55_06835 | QMY55_06840 | QMY55_06845 | QMY55_06850 | QMY55_06855 | QMY55_06860 | QMY55_06865 | QMY55_06870 |
|  | 92 | 94 | 90 | 93 | 96 | 95 | 95 | 86 | 92 | 94 | 88 | 88 | 96 | 83 | 95 | 70 |

| TA441 | ScdA | TesJ | TesI | TesH | TesA2 | TesA1 | TesD | TesE | TesF | TesG |
| --- | --- | --- | --- | --- | --- | --- | --- | --- | --- | --- |
| CNB-2 | CtCNB1_1360 | CtCNB1_1359 | CtCNB1_1358 | CtCNB1_1357 | CtCNB1_1356 | CtCNB1_1355 | CtCNB1_1354 | CtCNB1_1353 | CtCNB1_1352 | CtCNB1_1351 |
|  | 98 | 99 | 96 | 99 | 98 | 99 | 99 | 81 | 100 | 97 |
| DSMZ145 | SR914_26860 | SR914_26865 | SR914_26870 | SR914_26875 | SR914_26880 | SR914_26885 | SR914_26890 | SR914_26895 | tesF | tesG |
|  | 94 | 94 | 92 | 93 | 94 | 95 | 97 | 92 | 97 | 97 |
| TK102 | O987_07775 | O987_07770 | O987_07765 | O987_07760 | O987_07755 | O987_07750 | O987_07745 | O987_07740 | O987_07735 | O987_07730 |
|  | 94 | 94 | 92 | 94 | 94 | 95 | 97 | 92 | 97 | 97 |
| CJ34 | ctg1_86 | ctg1_85 | ctg1_84 | ctg1_83 | ctg1_82 | ctg1_81 | ctg1_80 | ctg1_79 | ctg1_78 | ctg1_77 |
|  | 88 | 94 | 88 | 92 | 94 | 96 | 96 | 95 | 97 | 96 |
| ZM22 | QMY55_07055 | QMY55_07050 | QMY55_07045 | QMY55_07040 | QMY55_07035 | QMY55_07030 | QMY55_07025 | QMY55_07020 | QMY55_07015 | dmpG |
|  | 89 | 90 | 85 | 89 | 93 | 94 | 95 | 94 | 97 | 95 |

| ChsH2 | ChsE1 | ChsH1 | Ltp2 | ChsE2 |
| --- | --- | --- | --- | --- |
| CtCNB1_1335 | CtCNB1_1334 | CtCNB1_1333 | CtCNB1_1332 | CtCNB1_1331 |
| 98 | 98 | 99 | 100 | 99 |
| SR914_26990 | SR914_26995 | SR914_27000 | SR914_27005 | SR914_27010 |
| 94 | 96 | 98 | 98 | 97 |
| O987_07650 | O987_07645 | O987_07640 | O987_07635 | O987_07630 |
| 94 | 78 | 98 | 99 | 97 |
| ctg1_57 | ctg1_58 | ctg1_59 | ctg1_60 | ctg1_61 |
| 86 | 82 | 91 | 96 | 93 |
| QMY55_06940 | QMY55_06935 | QMY55_06930 | QMY55_06925 | QMY55_06920 |
| 88 | 85 | 80 | 76 | 66 |

### Comparison of the steroid degradation gene clusters II: other *Comamonas* strains (Fig. 5; identities are shown in Table 4)

To examine whether other *Comamonas* species to possess a steroid degradation gene mega-cluster similar to that of TA441, we conducted BLAST searches using the DNA sequences from *tesB* to *tesR* and the amino acid sequences of TesH, ScdL1, and ScdY. These three enzymes were selected because they are characteristic enzymes associated with steroid degradation: ScdL1 and ScdY are involved in B-, C-, and D-ring degradation, whereas TesH is a Δ1-dehydrogenase involved in A- and B-ring cleavage. All the four search results were similar and largely overlapping. We then obtained approximately 200-kb DNA regions from the *Comamonas* strains as described above and subjected these regions to CAGECAT analysis.

**Fig. 5.**
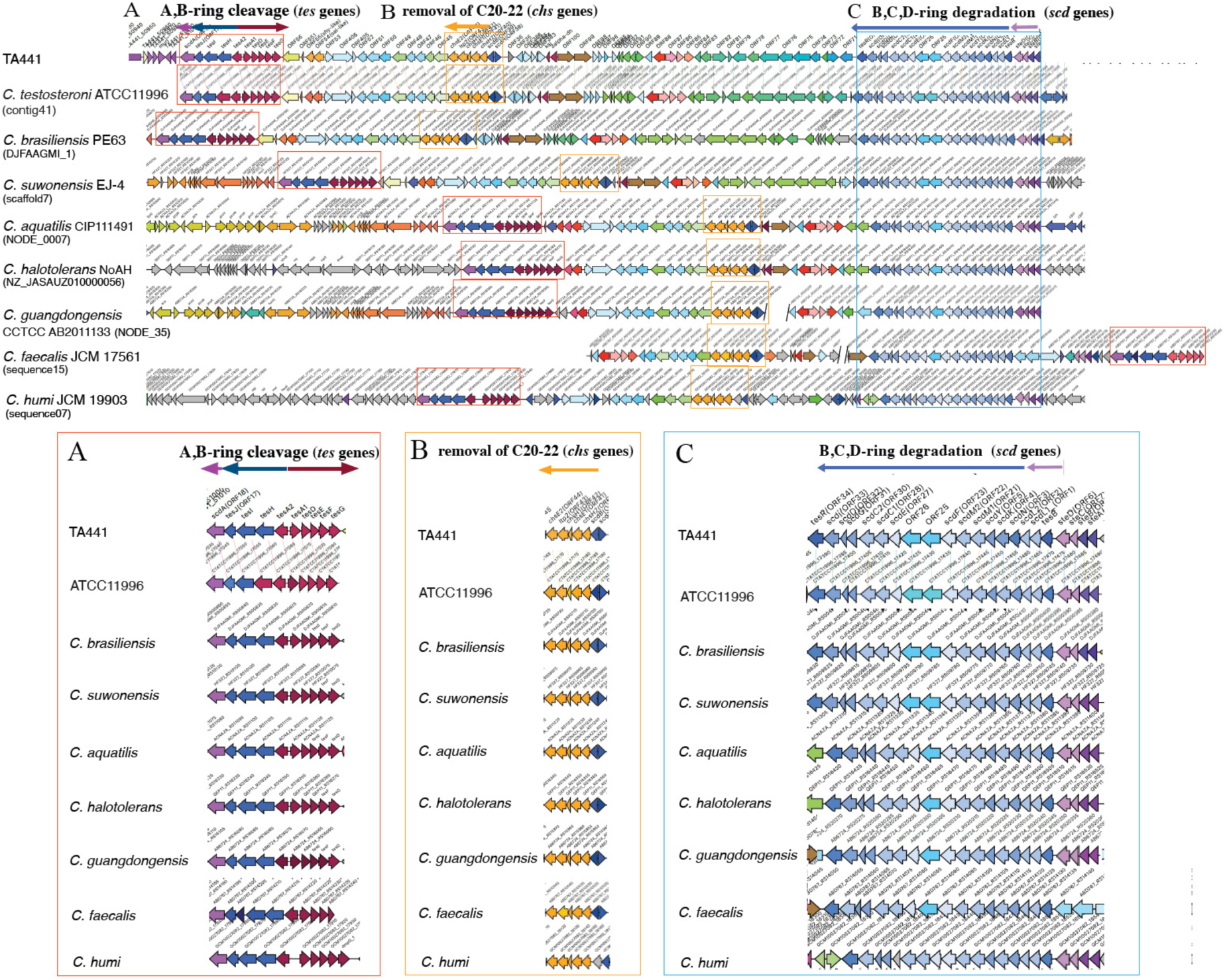
Comparison of the steroid degradation gene clusters of *Comamonas* strains other than those presented in Fig. 4 using CAGECAT. The genes involved in A- and B-ring cleavage (*tes* genes) are shown in dark blue (*tesHIJ*) and dark red (*tesA1A2DEFG*); the CoA transferase acting after A- and B-ring cleavage (*scdA*) in pink; genes involved in B-, C-, and D-ring degradation (*scd* genes) in light blue; and *chs* genes in orange. The colors of genes outside the boxes do not necessarily indicate their functions and are used to indicate similar genes among the strains. Identities between TA441 steroid degradation enzymes and the corresponding enzymes in other strains are listed in Table 4.

**Table 4.**
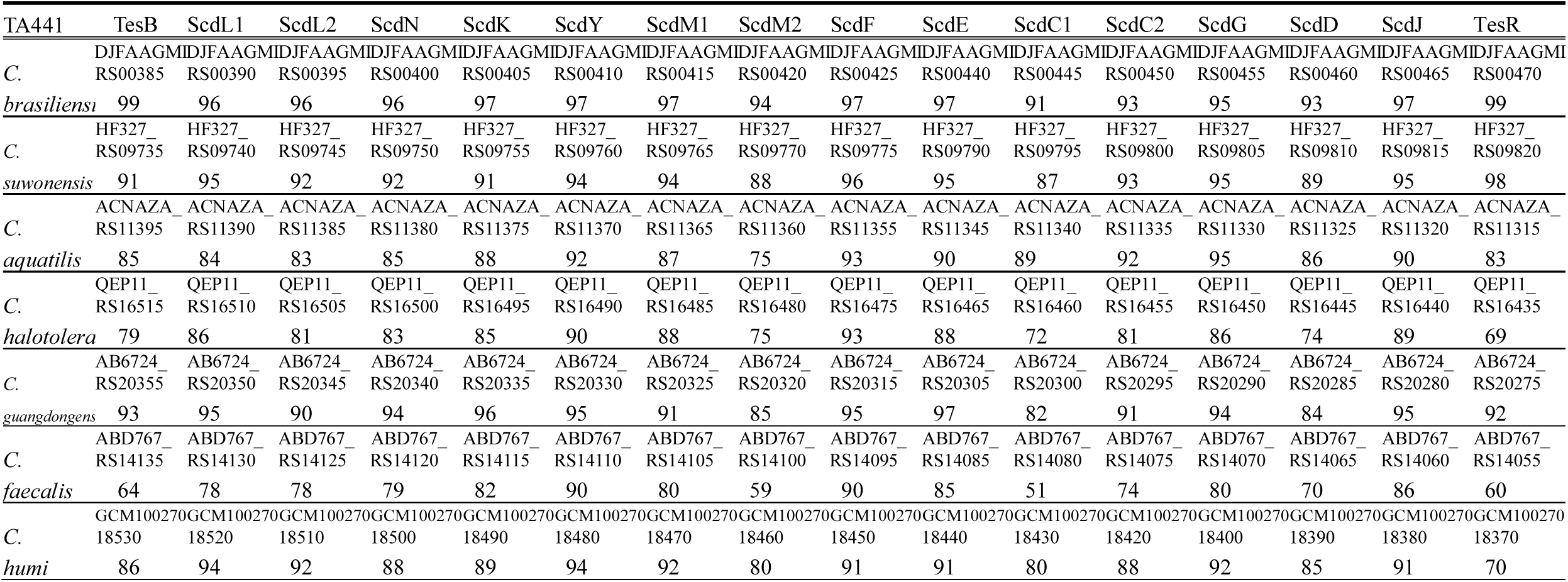
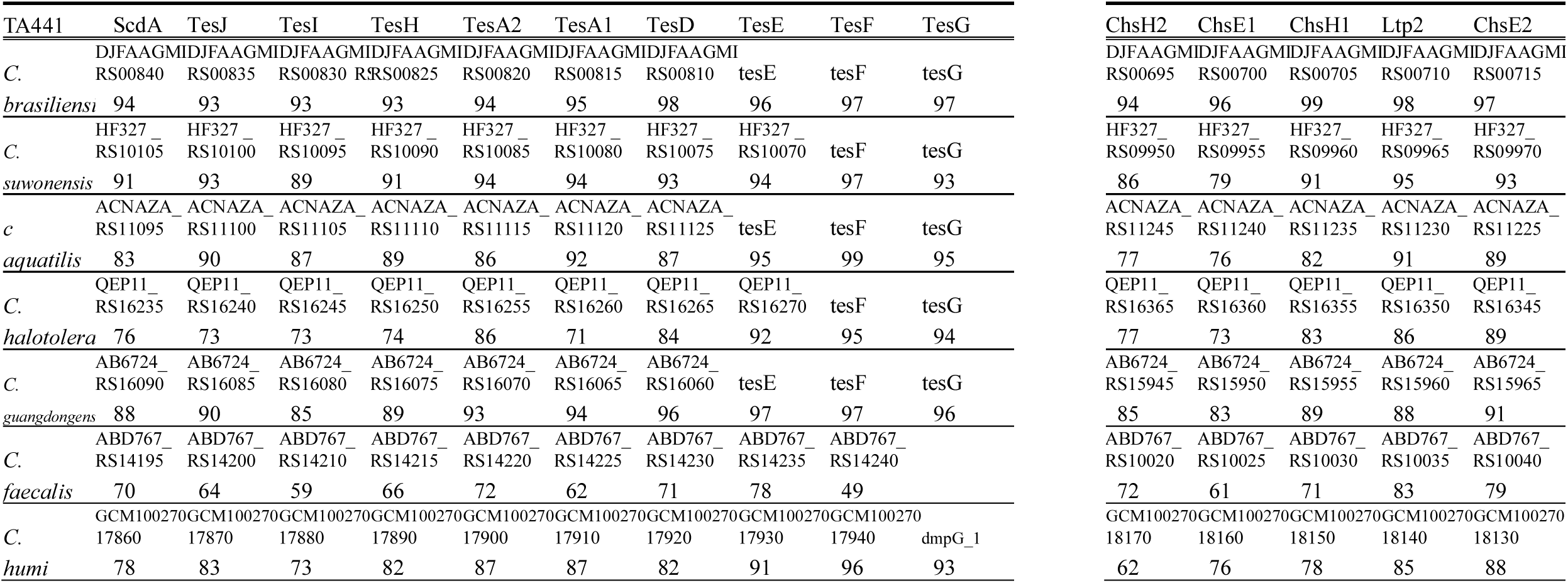
Amino acid identity (%) between the enzymes in Fig. 5 (locus tag or gene name) and corresponding enzymes in TA441.

As a result, *C. testosteroni* ATCC11996 (50), *C. brasiliensis* PE63 (55), *C. suwonensis* EJ-4(56), *C. aquatilis* CIP111491(57), *C. halotolerans* NoAH6 (58), *C. guangdongensis* CCTCC AB2011133 (59), *C. faecalis* JCM 17561 (60), and *C. humi* JCM 19903 (61) were found to possess a steroid degradation gene mega-cluster similar to that of TA441 (Fig. 5; identities between the corresponding enzymes and those of TA441 are listed in Table 4). *C. testosteroni* ATCC11996 is a type strain of *C. testosteroni* and a representative steroid-degrading bacterium that has been studied for decades. ATCC11996 was included in this analysis because its complete genome sequence was not available to subject to the Mauve analysis. As expected from our previous identification of the *tesB* to *tesR* region (19), the steroid degradation gene mega-clusters of TA441 and ATCC11996 were quite similar.

All the strains possessed *tes*, *scd*, and *chs* genes, and the locations of these genes within the cluster were highly similar. Most of these *Comamonas* strains were closely related to TA441 in the 16S rRNA gene phylogenetic tree (indicated with light blue boxes in Fig. 3A), except for *C. faecalis* JCM 17561, in which the location of the *tes* genes differed from those in the other strains (Fig. 5).

*C. testosteroni* is not generally considered to inhabit the intestine, despite its high cholic acid degradation ability and reports of more than 50 cases of its isolation from humans (62). Among the *Comamonas* strains analyzed here, *C. faecalis* JCM 17561 was isolated from domestic pig feces (60), and *C. halotolerans* NoAH (KACC 23135, JCM 35999) was isolated from a fecal sample of a zoo animal, *Naemorhedus caudatus* (58). Amino acid identities between TesH of TA441 and the corresponding enzymes in the other *Comamonas* strains were approximately 89–99%, except for *C. faecalis* (66%) and *C. humi* (88%) (Tables 3 and 4).

### Comparison of the steroid degradation gene clusters III: β-Proteobacteria except for *Comamonas* sp. (Fig. 6A and B; identities are shown in Tables 5 and 6)

We conducted BLAST searches using the DNA sequences from *tesB* to *tesR* (*scd* genes) and from *scdA* to *tesG* (*tes* genes), excluding *Comamonas* strains, to identify similar genes in other β-Proteobacteria. The sequence of *Alicycliphilus denitrificans* (63), showed the highest similarity to the overall *tesB* to *tesR* region, whereas the sequence of *Cupriavidus necator* NH9 (64) showed high similarity but covered less than 20% of the compared region. The sequence of *Diaphorobacter* sp. HDW4A (65) showed the highest similarity to the *scdA* to *tesG* region. Approximately 200-kb DNA regions containing these sequences were analyzed using CAGECAT in the same manner as for the *Comamonas* strains (Fig. 6A; identities between the corresponding enzymes and those of TA441 are listed in Tables 5 and 6).

**Fig. 6.**
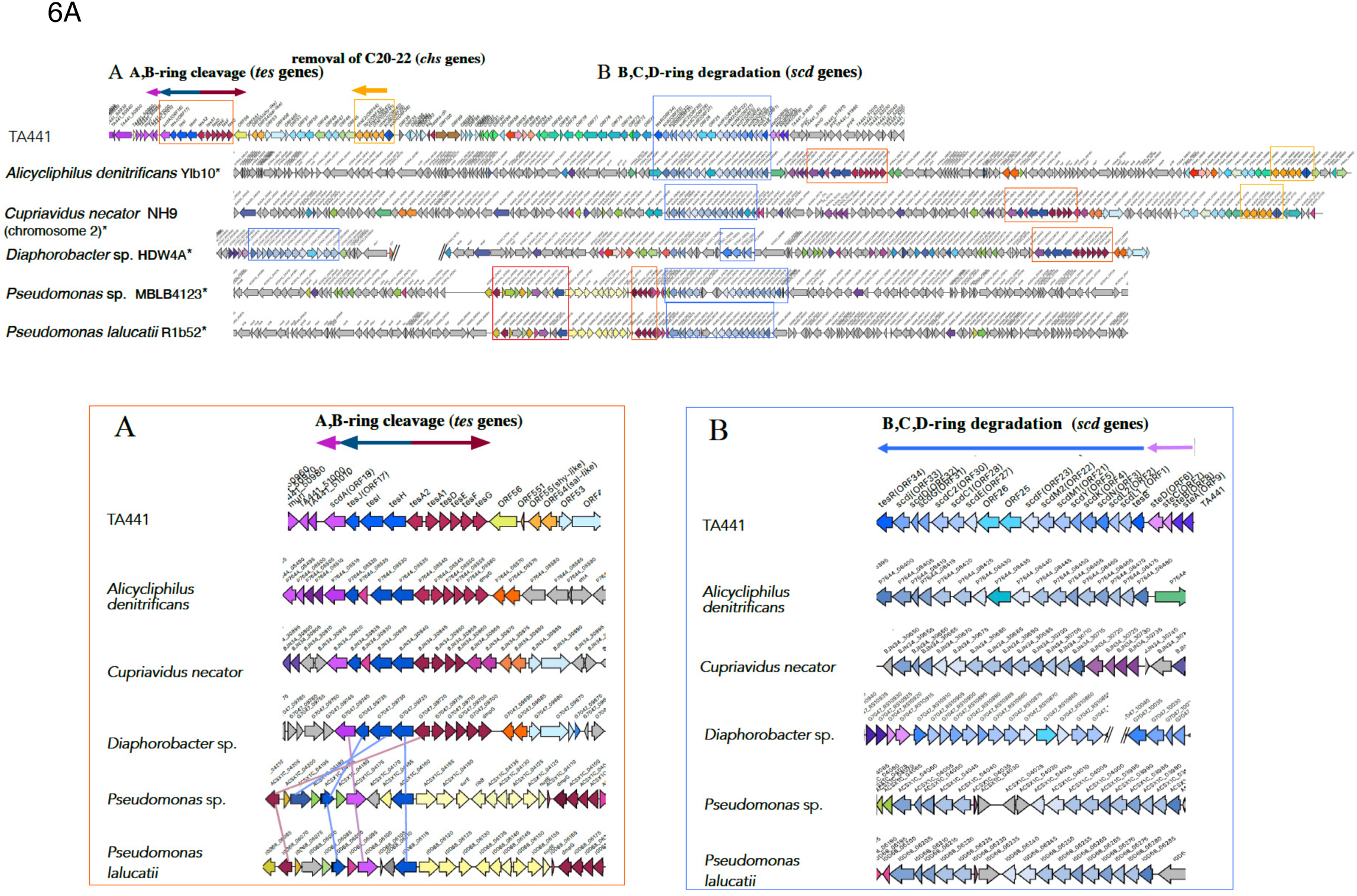
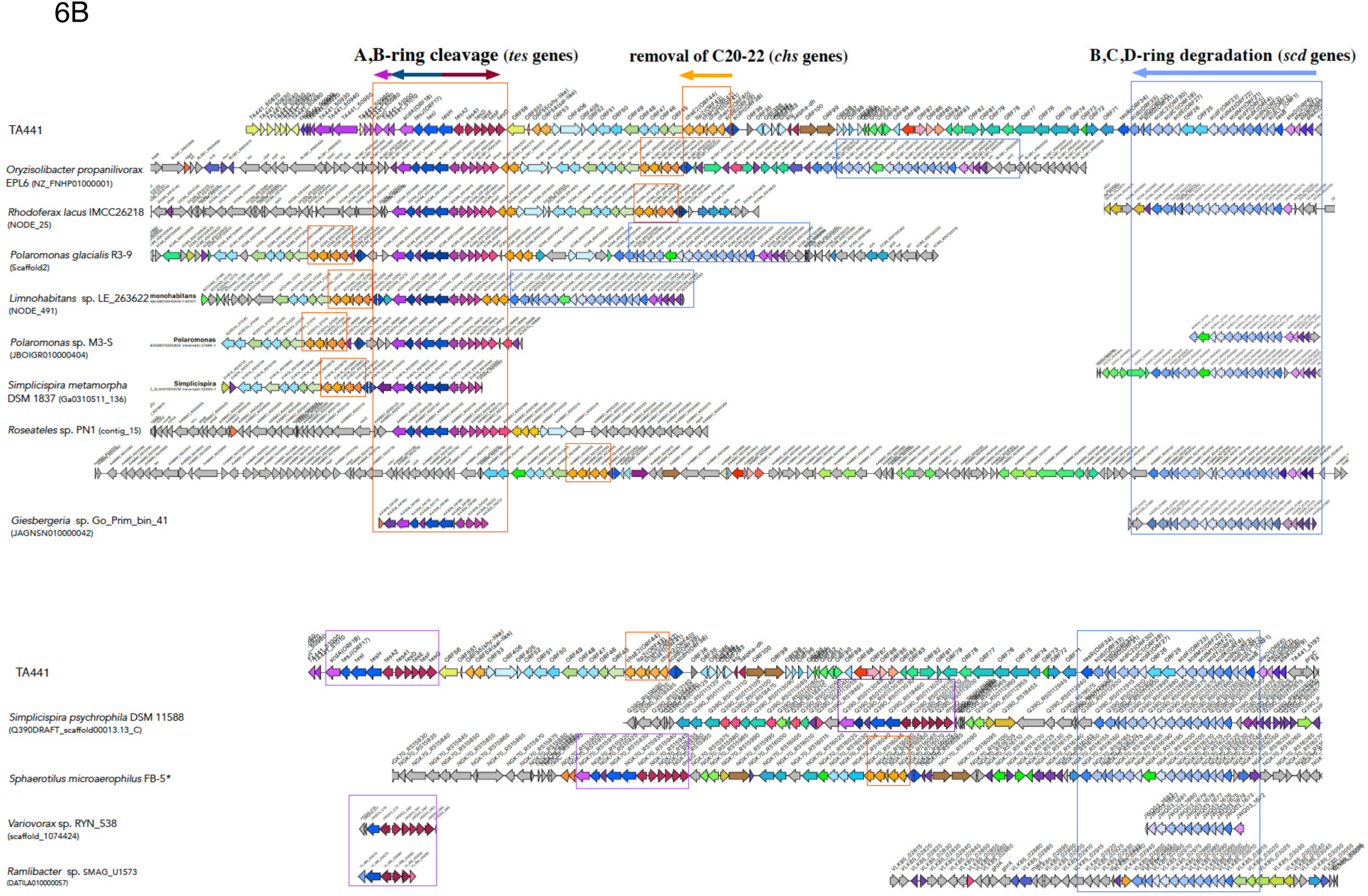
Comparison of the steroid degradation gene clusters of β-Proteobacteria except for *Comamonas* sp. using CAGECAT. **6A**: The putative clusters selected based on higher homology to the DNA sequences of *scdA–tesG* (*Diaphorobacter* sp. HDW4A) and *tesB–tesR* (others). Two *Pseudomonas* (γ-Proteobacteria) strains are presented here because they showed higher similarity than other β-Proteobacteria. **6B**: The putative clusters selected based on higher homology to the amino acid sequences of TesH, ScdL1, or ScdY. *: strains with complete genome sequences. The genes involved in A-and B-ring cleavage (*tes* genes) are shown in dark blue (*tesHIJ*) and dark red (*tesA1A2DEFG*); the CoA transferase acting after A- and B-ring cleavage (*scdA*) in pink; genes involved in B-, C-, and D-ring degradation (*scd* genes) in light blue; and *chs* genes in orange. The colors of genes outside the boxes do not necessarily indicate their functions and are used to indicate similar genes among the strains. Identities between the TA441 enzymes and the corresponding enzymes in other strains are listed in Tables 5 and 6.

**Table 5.**
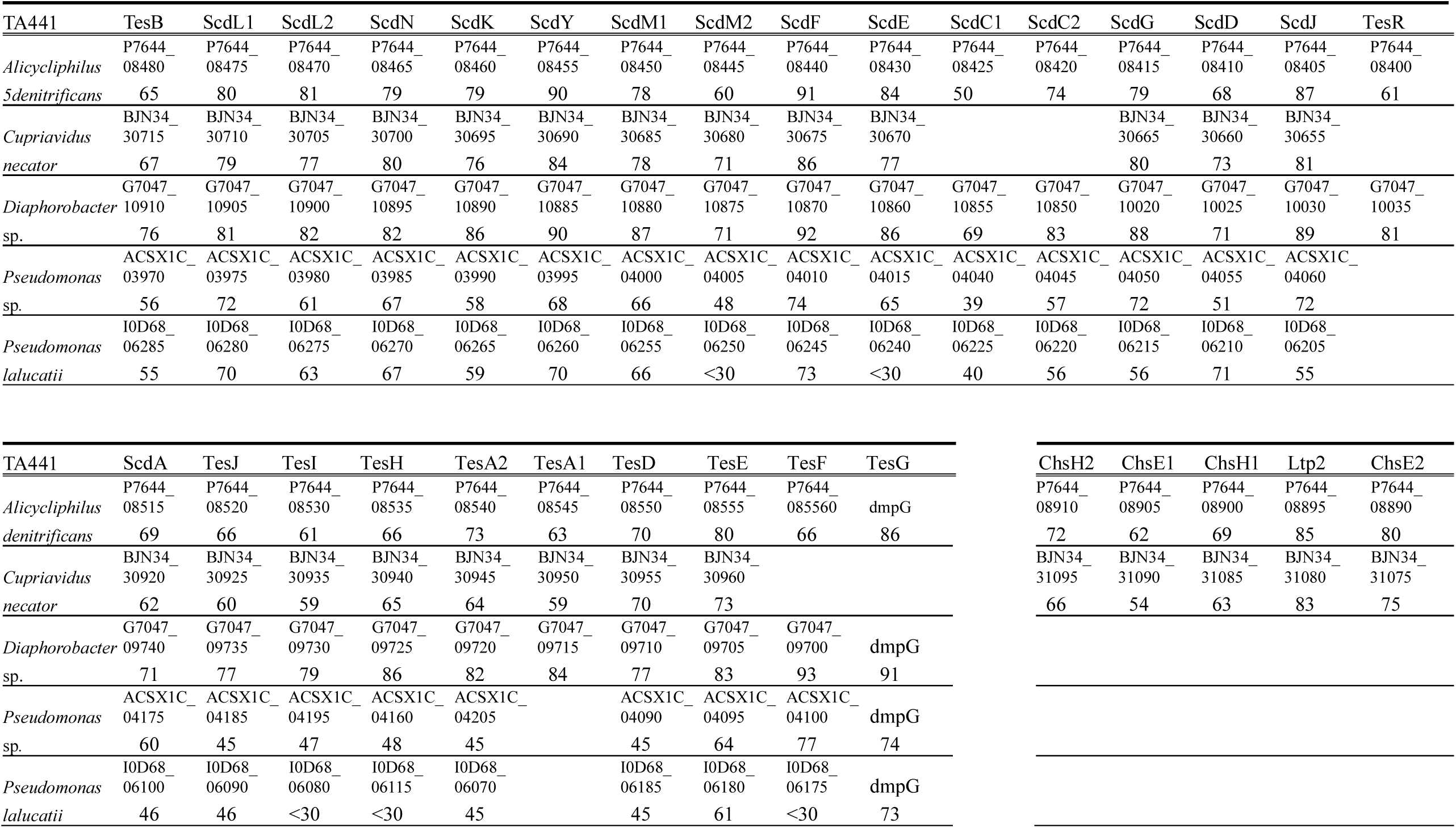
Amino acid identity (%) between the enzymes in Fig. 6A (locus tag or gene name) and corresponding enzymes in TA441.

**Table 6.** Amino acid identity (%) between the enzymes in Fig. 6B (locus tag or gene name) and corresponding enzymes in TA441.

| TA441 | TesB | ScdL1 | ScdL2 | ScdN | ScdK | ScdY | ScdM1 | ScdM2 | ScdF | ScdE | ScdC1 | ScdC2 | ScdG | ScdD | ScdJ | TesR |
| --- | --- | --- | --- | --- | --- | --- | --- | --- | --- | --- | --- | --- | --- | --- | --- | --- |
| *Oryzolibacter* | BLQ61_<br>RS00085 | BLQ61_<br>RS00090 | BLQ61_<br>RS00095 | BLQ61_<br>RS00100 | BLQ61_<br>RS00105 | BLQ61_<br>RS00110 | BLQ61_<br>RS00115 | BLQ61_<br>RS00120 | BLQ61_<br>RS00125 | BLQ61_<br>RS00135 | BLQ61_<br>RS00140 | BLQ61_<br>RS00145 | BLQ61_<br>RS00150 | BLQ61_<br>RS00155 | BLQ61_<br>RS00160 | BLQ61_<br>RS00165 |
| *propanilivorax* | 75 | 82 | 84 | 85 | 83 | 92 | 87 | 73 | 94 | 92 | 64 | 81 | 87 | 73 | 86 | 76 |
| *Rhodoferrax* | DIC66_<br>RS20770 | DIC66_<br>RS20765 | DIC66_<br>RS20760 | DIC66_<br>RS20755 | DIC66_<br>RS20750 | DIC66_<br>RS20745 | DIC66_<br>RS20740 | DIC66_<br>RS20735 | DIC66_<br>RS20730 | DIC66_<br>RS20725 | DIC66_<br>RS20720 | DIC66_<br>RS20715 | DIC66_<br>RS20710 | DIC66_<br>RS20705 | DIC66_<br>RS20700 | DIC66_<br>RS20695 |
| *lacus* | 69 | 80 | 80 | 78 | 82 | 85 | 80 | 63 | 87 | 79 | 50 | 71 | 81 | 66 | 85 | 61 |
| *Polaromonas* | EC86_<br>RS0120320 | EC86_<br>RS0120325 | EC86_<br>RS0120330 | EC86_<br>RS0120335 | EC86_<br>RS0120340 | EC86_<br>RS0120345 | EC86_<br>RS0120350 | EC86_<br>RS0120355 | EC86_<br>RS0120360 | EC86_<br>RS0120365 | EC86_<br>RS0120375 | EC86_<br>RS0120380 | EC86_<br>RS0120385 | EC86_<br>RS0120390 | EC86_<br>RS0120395 | EC86_<br>RS0120400 |
| *glacialis* | 67 | 74 | 73 | 78 | 78 | 79 | 88 | 60 | 91 | 74 | 51 | 72 | 82 | 69 | 86 | 60 |
| *Limnohabitans* | ACOVN3_<br>12415 | ACOVN3_<br>12410 | ACOVN3_<br>12405 | ACOVN3_<br>12400 | ACOVN3_<br>12395 | ACOVN3_<br>12390 | ACOVN3_<br>12385 | ACOVN3_<br>12380 | ACOVN3_<br>12375 | ACOVN3_<br>12370 | ACOVN3_<br>12360 | ACOVN3_<br>12355 | ACOVN3_<br>12350 | ACOVN3_<br>12345 | ACOVN3_<br>1238540 | ACOVN3_<br>12335 |
| *sp.* | 63 | 77 | 75 | 79 | 77 | 86 | 76 | 59 | 87 | 70 | 45 | 68 | 82 | 64 | 86 | 57 |
| *Polaromonas* | ACREXV_<br>07185 | ACREXV_<br>07180 | ACREXV_<br>07175 | ACREXV_<br>07170 | ACREXV_<br>07165 | ACREXV_<br>07160 | ACREXV_<br>07155 | ACREXV_<br>07150 | ACREXV_<br>07145 | ACREXV_<br>07140 | ACREXV_<br>07130 |  |  |  |  |  |
| *sp.* | 67 | 78 | 76 | 79 | 81 | 88 | 79 | 61 | 88 | 74 | 46 |  |  |  |  |  |
| *Simplicispira* | D3C67_<br>RS05905 | D3C67_<br>RS05900 | D3C67_<br>RS05895 | D3C67_<br>RS05890 | D3C67_<br>RS05885 | D3C67_<br>RS05880 | D3C67_<br>RS05875 | D3C67_<br>RS05870 | D3C67_<br>RS05865 | D3C67_<br>RS05860 | D3C67_<br>RS05855 | D3C67_<br>RS05845 | D3C67_<br>RS05840 | D3C67_<br>RS05835 | D3C67_<br>RS05830 | D3C67_<br>RS05825 |
| *metamorphia* | 67 | 78 | 73 | 79 | 81 | 89 | 80 | 62 | 90 | 73 | 51 | 72 | 83 | 70 | 86 | 57 |
| *Roseateles* | AABB87_<br>RS16215 | AABB87_<br>RS16210 | AABB87_<br>RS16205 | AABB87_<br>RS16200 | AABB87_<br>RS16195 | AABB87_<br>RS16190 | AABB87_<br>RS16185 | AABB87_<br>RS16180 | AABB87_<br>RS16175 | AABB87_<br>RS16165 | AABB87_<br>RS16155 | AABB87_<br>RS16150 | AABB87_<br>RS16145 | AABB87_<br>RS16140 | AABB87_<br>RS16135 | AABB87_<br>RS16130 |
| *sp.* | 63 | 79 | 79 | 79 | 80 | 87 | 77 | 63 | 89 | 79 | 41 | 58 | 81 | 71 | 96 | 60 |
| *Giesbergeria* | KA309_<br>01385 | KA309_<br>01390 | KA309_<br>01395 | KA309_<br>01400 | KA309_<br>01405 | KA309_<br>01410 | KA309_<br>01415 | KA309_<br>01420 | KA309_<br>01425 | KA309_<br>01430 | KA309_<br>01435 | KA309_<br>01440 | KA309_<br>01445 | KA309_<br>01450 | KA309_<br>01455 | KA309_<br>01460 |
| *sp.* | 67 | 74 | 74 | 79 | 81 | 88 | 78 | 59 | 91 | 72 | 52 | 70 | 84 | 71 | 85 | 59 |
| *Simplicispira* | Q390_<br>RS0112840 | Q390_<br>RS0112845 | Q390_<br>RS0112850 | Q390_<br>RS0112855 | Q390_<br>RS0112860 | Q390_<br>RS0112865 | Q390_<br>RS0112870 | Q390_<br>RS0112875 | Q390_<br>RS0112880 | Q390_<br>RS0112885 | Q390_<br>RS0112890 | Q390_<br>RS0112895 | Q390_<br>RS0112900 | Q390_<br>RS0112905 | Q390_<br>RS0112910 | Q390_<br>RS0112915 |
| *psychrophila* | 67 | 76 | 74 | 78 | 79 | 88 | 79 | 60 | 91 | 71 | 48 | 69 | 83 | 70 | 85 | 58 |
| *Sphaerotilus* | NGK70_<br>RS16240 | NGK70_<br>RS16235 | NGK70_<br>RS16230 | NGK70_<br>RS16225 | NGK70_<br>RS16220 | NGK70_<br>RS16215 | NGK70_<br>RS16210 | NGK70_<br>RS16205 | NGK70_<br>RS16200 | NGK70_<br>RS16190 | NGK70_<br>RS16185 | NGK70_<br>RS16180 | NGK70_<br>RS16175 | NGK70_<br>RS16170 | NGK70_<br>RS16165 | NGK70_<br>RS16160 |
| *microaerophilus* | 62 | 79 | 80 | 78 | 79 | 88 | 79 | 57 | 88 | 79 | 42 | 57 | 81 | 70 | 86 | 65 |
| *Variovorax* | JWQ03_<br>1673 | JWQ03_<br>1674 | JWQ03_<br>1675 | JWQ03_<br>1676 | JWQ03_<br>1677 | JWQ03_<br>1678 | JWQ03_<br>1679 | JWQ03_<br>1680 | JWQ03_<br>1681 | JWQ03_<br>1682 |  |  |  |  |  |  |
| *sp.* | 67 | 77 | 72 | 79 | 80 | 88 | 80 | 61 | 88 | 72 |  |  |  |  |  |  |
| *Ramlibacter* sp. | VLK85_<br>03010 | VLK85_<br>03005 | VLK85_<br>03000 | VLK85_<br>02995 | VLK85_<br>02990 | VLK85_<br>02985 | VLK85_<br>02980 | VLK85_<br>02975 | VLK85_<br>02970 | VLK85_<br>02965 |  |  | VLK85_<br>02960 | VLK85_<br>02955 | VLK85_<br>02950 |  |
| SMAG U15732 | 63 | 78 | 69 | 74 | 75 | 81 | 77 | 61 | 82 | 77 |  |  | 77 | 65 | 80 |  |

| TA441 | ScdA | TesJ | TesI | TesH | TesA2 | TesA1 | TesD | TesE | TesF | TesG |
| --- | --- | --- | --- | --- | --- | --- | --- | --- | --- | --- |
| *Oryzolibacter* | BLQ61_ RS00370 | BLQ61_ RS00365 | BLQ61_ RS00360 | BLQ61_ RS00355 | BLQ61_ RS00350 | BLQ61_ RS00345 | BLQ61_ RS00340 | BLQ61_ RS00335 | tesF | dmpG |
| *propanilivorax* | 75 | 73 | 76 | 81 | 84 | 77 | 80 | 84 | 94 | 87 |
| *Rhodoferrax* | DIC66_ RS14995 | DIC66_ RS14990 | DIC66_ RS14980 | DIC66_ RS14975 | DIC66_ RS14970 | DIC66_ RS14965 | DIC66_ RS14960 | DIC66_ RS14955 | DIC66_ RS14950 | dmpG |
| *lacus* | 63 | 63 | 60 | 67 | 73 | 63 | 69 | 77 | 72 | 64 |
| *Polaromonas* | EC86_ RS0120505 | EC86_ RS0120500 | EC86_ RS0120490 | EC86_ RS0120485 | EC86_ RS0120480 | EC86_ RS0120475 | EC86_ RS0120470 | EC86_ RS0120465 | EC86_ RS0120460 | dmpG |
| *glacialis* | 68 | 66 | 59 | 65 | 66 | 66 | 73 | 78 | 79 | 85 |
| *Limnohabitans* | ACOVN3_ 12275 | ACOVN3_ 12280 | ACOVN3_ 12290 | ACOVN3_ 12295 | ACOVN3_ 12300 | ACOVN3_ 12305 | ACOVN3_ 12310 | ACOVN3_ 12315 |  |  |
| *sp.* | 63 | 63 | 59 | 66 | 66 | 65 | 74 | 75 |  |  |
| *Polaromonas* | ACREXV_ 06955 | ACREXV_ 06950 | ACREXV_ 06940 | ACREXV_ 06935 | ACREXV_ 06930 | ACREXV_ 06925 | ACREXV_ 06920 | ACREXV_ 06915 | ACREXV_ 06910 | dmpG |
| *sp.* | 66 | 65 | 61 | 67 | 69 | 57 | 73 | 79 | 82 | 86 |
| *Simplicispira* | EV674_ RS15630 | EV674_ RS15625 | EV674_ RS15615 | EV674_ RS15610 | EV674_ RS15605 | EV674_ RS15600 | EV674_ RS15595 | EV674_ RS15590 | - |  |
| *metamorphia* | 68 | 66 | 60 | 66 | 66 | 66 | 72 | 78 |  |  |
| *Roseateles* | AABB87_ RS20130 | AABB87_ RS20135 | AABB87_ RS20145 | AABB87_ RS20150 | AABB87_ RS20155 | AABB87_ RS20160 | AABB87_ RS20165 | AABB87_ RS20170 | AABB87_ RS20175 | dmpG |
| *sp.* | 78 | 83 | 73 | 82 | 87 | 87 | 82 | 91 | 96 | 93 |
| *Giesbergeria* | KA309_ 04165 | KA309_ 04170 | KA309_ 04180 | KA309_ 04185 | KA309_ 04195 | KA309_ 04200 | KA309_ 04205 | KA309_ 04210 |  |  |
| *sp.* | 66 | 67 | 61 | 66 | 62 | 64 | 73 | 77 |  |  |
| *Simplicispira* | Q390_ RS011304 | Q390_ RS011304 | Q390_ RS011303 | Q390_ RS011302 | Q390_ RS011302 | Q390_ RS011301 | Q390_ RS011301 | Q390_ RS011300 | Q390_ RS011300 | dmpG |
| *psychrophila* | 66 | 66 | 60 | 64 | 66 | 64 | 73 | 78 | 80 | 85 |
| *Sphaerotilus* | NGK70_ RS15915 | NGK70_ RS15920 | NGK70_ RS15930 | NGK70_ RS15935 | NGK70_ RS15940 | NGK70_ RS15945 | NGK70_ RS15955 | NGK70_ RS15960 | NGK70_ RS15965 | dmpG |
| *microaerophilus* | 66 | 66 | 61 | 66 | 71 | 63 | 41 | 78 | 82 | 85 |
| *Variovorax* |  |  | JWQ03_ 578 | JWQ03_ 579 | JWQ03_ 580 | JWQ03_ 581 | JWQ03_ 582 | JWQ03_ 583 |  |  |
| *sp.* |  |  | 58 | 68 | 65 | 73 | 78 | 85 |  |  |
| *Ramlibacter* sp. |  |  |  | VLJ86_ 00675 | HSW 00680 | HSW 00685 | HSW 00690 | HSW 00695 |  |  |
| SMAG U15732 |  |  |  | 65 | 68 | 62 | 72 | 66 |  |  |

| ChsH2 | ChsE1 | ChsH1 | Ltp2 | ChsE2 |
| --- | --- | --- | --- | --- |
| BLQ61_ RS00240 | BLQ61_ RS00245 | BLQ61_ RS00250 | BLQ61_ RS00255 | BLQ61_ RS00260 |
| 77 | 74 | 77 | 91 | 88 |
| DIC66_ RS14855 | DIC66_ RS14860 | DIC66_ RS14865 | DIC66_ RS14870 | DIC66_ RS14875 |
| 67 | 60 | 66 | 88 | 79 |
| EC86_ RS0120540 | EC86_ RS0120545 | EC86_ RS0120550 | EC86_ RS0120555 | EC86_ RS0120560 |
| 69 | 58 | 63 | 85 | 80 |
| ACOVN3_ 12255 | COVN3_ 12250 | ACOVN3_ 12245 | ACOVN3_ 12240 | ACOVN3_ 12235 |
| 71 | 56 | 64 | 85 | 79 |
| ACREXV_ 06990 | ACREXV_ 06995 | ACREXV_ 07000 | ACREXV_ 07005 | ACREXV_ 07010 |
| 69 | 59 | 64 | 87 | 80 |
| EV674_ RS15650 | EV674_ RS15655 | EV674_ RS15660 | EV674_ RS15665 | EV674_ RS15670 |
| 70 | 60 | 66 | 84 | 81 |
| AABB87_ RS15845 | AABB87_ RS15840 | AABB87_ RS15835 | AABB87_ RS15830 | AABB87_ RS15825 |
| 62 | 76 | 78 | 85 | 88 |

| Q390_ RS0113065 | Q390_ RS0113070 | Q390_ RS0113075 | Q390_ RS0113080 | Q390_ RS0113085 |
| --- | --- | --- | --- | --- |
| 68 | 55 | 65 | 86 | 81 |
| NGK70_ RS16065 | NGK70_ RS16060 | NGK70_ RS16055 | NGK70_ RS16050 | NGK70_ RS16045 |
| 72 | 64 | 73 | 88 | 82 |

All three strains possessed genes corresponding to the *scd* and *tes* genes. The *scd* genes of *Diaphorobacter* sp. HDW4A were divided into two smaller clusters, but all members of the *scd* cluster were found in the same order as those in TA441. In the BLAST search using *tesB* to *tesR*, many sequences from γ-Proteobacteria Pseudomonadaceae strains were identified as partially similar sequences. We compared the DNA regions of *Pseudomonas* sp. MBLB4123 (66) and *P. lalucatii* R1b52 (67) with the TA441 steroid degradation gene cluster using CAGECAT and found putative *scd* genes forming two smaller clusters, *tesB* to *scdE* and *scdCI* to *scdJ*, in both strains.

Because BLAST searches using nucleotide sequences did not perform as well as expected, we subsequently conducted BLAST searches using the amino acid sequences of TesH, ScdL1, and ScdY and analyzed the DNA regions containing the identified genes using CAGECAT.

Enzymes in many bacteria showed considerable homology to TesH and among them, the enzymes in *Diaphorobacter* sp. HDW4A, *Oryzisolibacter propanilivorax* EPL6 (68), *Rhodoferax lacus* IMCC26218 (69), *Polaromonas glacialis* R3-9 (70), *Limnohabitans* sp. LE_263622 (71), *Polaromonas* sp. M3-S (72), *Simplicispira metamorpha* DSM 1837 (73), *Roseateles* sp. PN1 (74), and *Giesbergeria* sp.

Go_Prim_bin_41 (75) showed highest homology to TesH. The BLAST search results obtained using ScdL1 or ScdY largely overlapping with those obtained using TesH. Therefore, *Simplicispira psychrophila* DSM 11588 (76), *Sphaerotilus* sp. FB-5 (77), *Variovorax* sp. RYN_538 (78), and *Ramlibacter* sp. SMAG_U15732 k141_119062 (79) from the search results with ScdL1 and ScdY were also subjected to the CAGECAT analysis. For the analysis, 200kb DNA region of bacteria with a gene encoding an enzyme corresponding to TesH, ScdL1, or ScdY (the region encoding the enzyme extended approximately 100 kb upstream and downstream from the beginning) was used (Fig. 6B; identities between the corresponding enzymes and those of TA441 are listed in Tables 5 and 6). When the genes were located on a short contig containing only part of the putative cluster, we searched other contigs for additional putative steroid degradation genes and found that all the strains possessed both *tes* and *scd* gene clusters. The structures of the clusters were quite similar in many strains, and even when some differences were observed, genes corresponding to those required for degradation of the sterane structure were present. These results suggest that all of these bacteria may possess the genetic capacity for aerobic steroid degradation.

All three enzymes, ScdL1, ScdY, and TesH were useful for identifying putative steroid degradation gene clusters. However, Δ1-dehydrogenases (TesH in TA441) are identified and well-studied in β-, γ-, and α-Proteobacteria and Actinomyces, whereas studies on enzymes corresponding to ScdL1 or ScdY are relatively limited. In addition, these two are the enzymes involved in β-oxidation, and when the identities of the corresponding enzymes are relatively low, it is difficult to determine whether they are involved in steroid degradation or in other metabolic pathways without analyzing the surrounding gene cluster. From these results, TesH, a Δ1-dehydrogenase, appeared to be more suitable marker for identifying bacterial aerobic steroid degradation gene clusters.

Therefore, in the following analyses of steroid degradation genes in other genera of bacteria, we basically used the amino acid sequence of Δ1-dehydrogenase to identify DNA regions for subsequent CAGECAT analysis.

Amino acid identities between TesH of TA441 and the corresponding enzymes in these β-Proteobacteria were approximately 65–80% (Tables 5 and 6).

### Comparison of the steroid degradation gene clusters IV: γ-Proteobacteria (**Fig. 7**; identities are shown in **Table 7**)

Aerobic bacterial steroid degradation has been reported in β-, γ-, and α-Proteobacteria and Actinomyces. Among the γ-Proteobacteria, steroid degradation has been studied in detail in *Pseudomonas stutzeri* Chol1 (formerly *Pseudomonas* sp. Chol1) (Fig. 7A) (80–85). In the genomic sequence of Chol1, *stdH* (*kstD1*), which corresponds to *tesH*, is missing (80). However, 632 bp downstream of the first “ATG” in the DNA region where *stdH* (*kstD1*) is expected to be located, a repeated sequence, “GGGCAACG,” is present (“GGGCAACGGGGCAACG”).

**Fig. 7.**
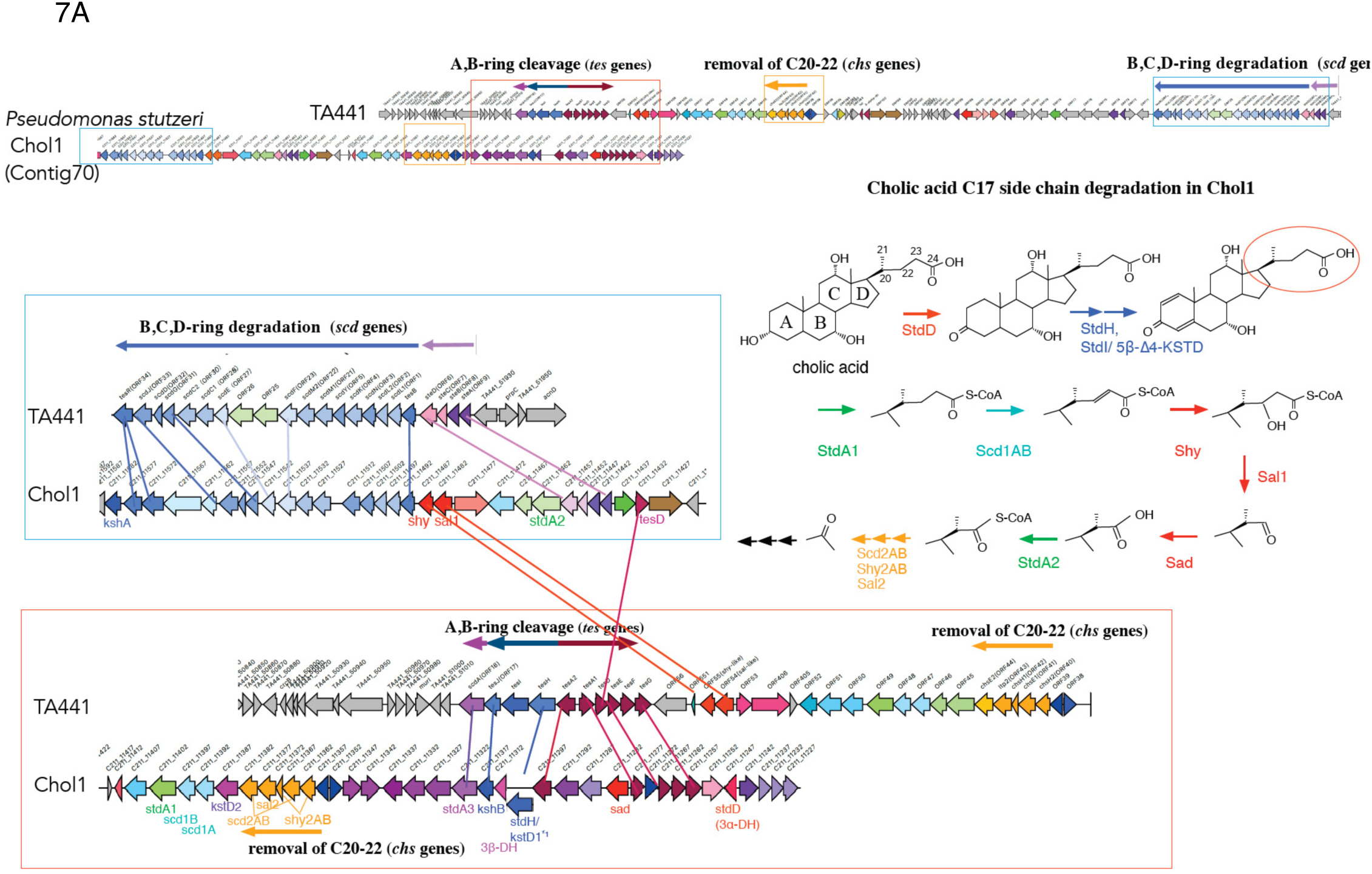
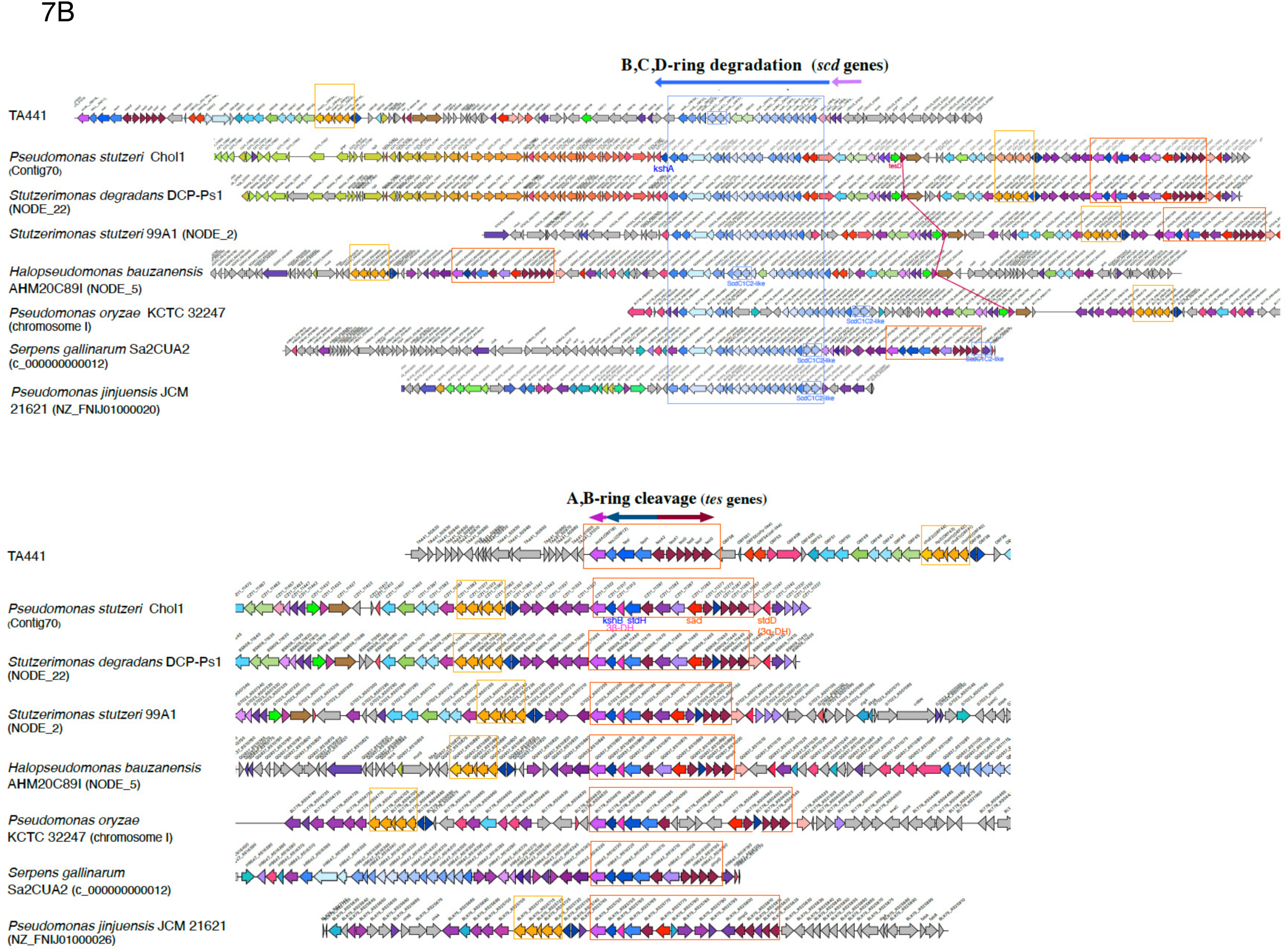
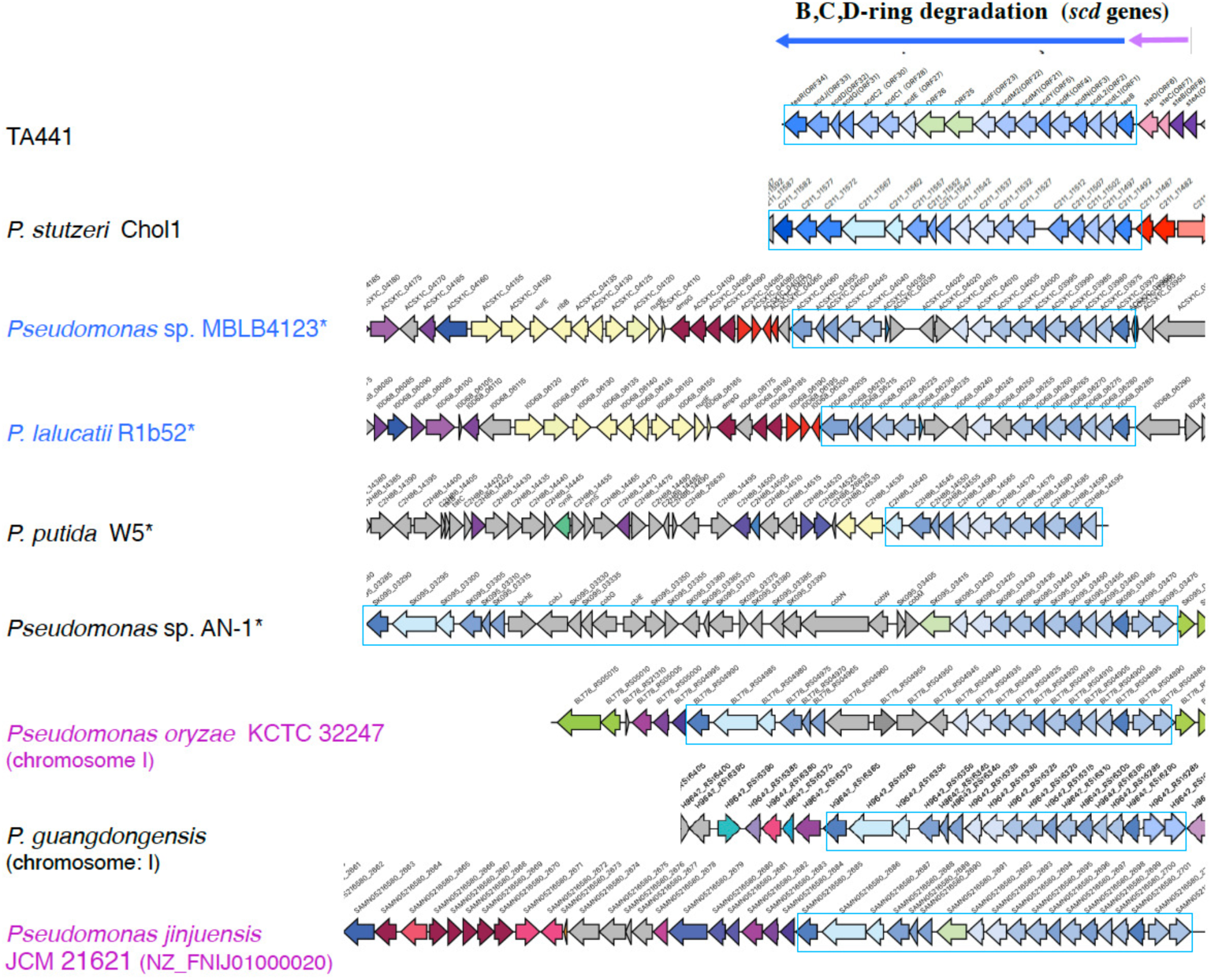
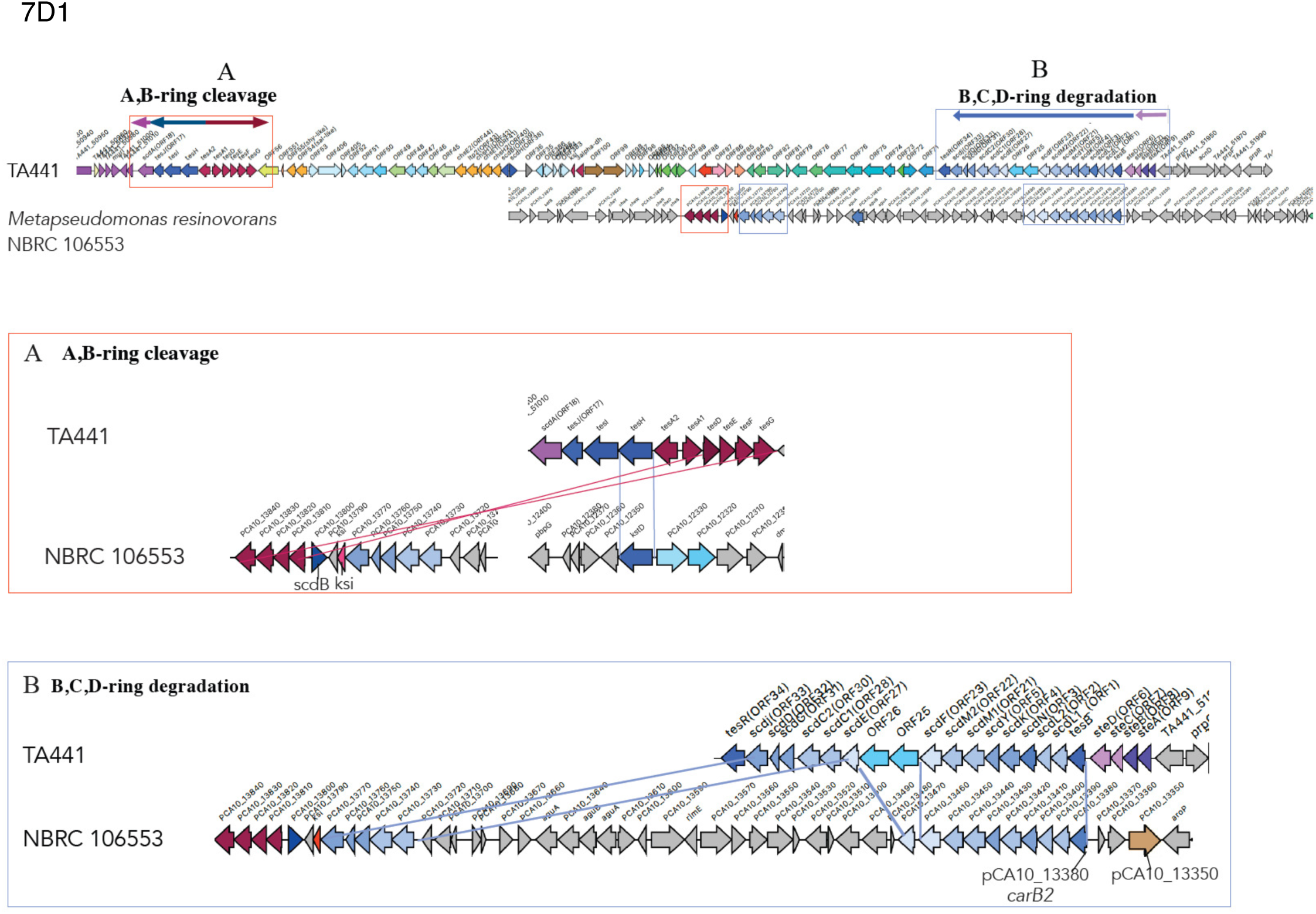
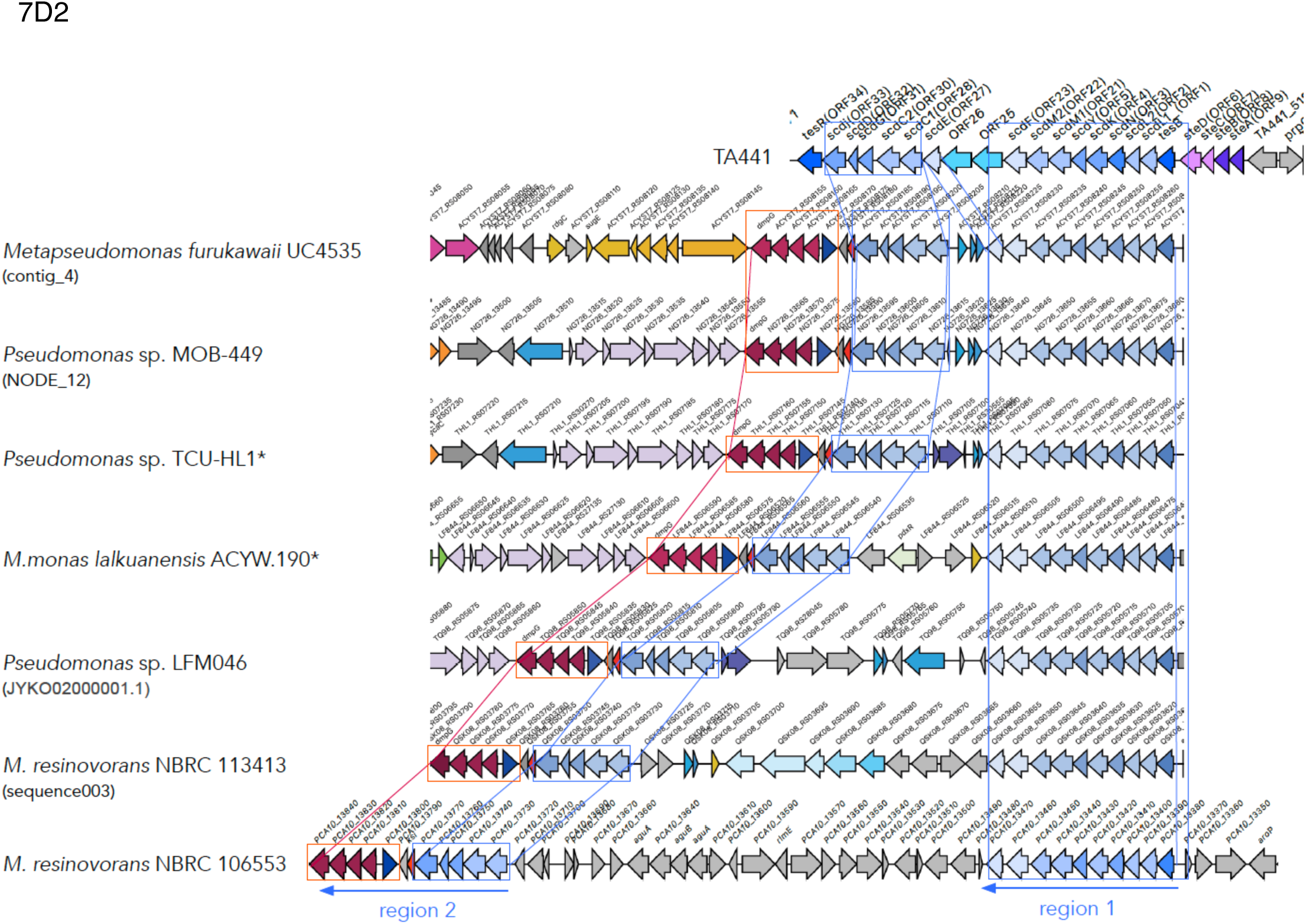
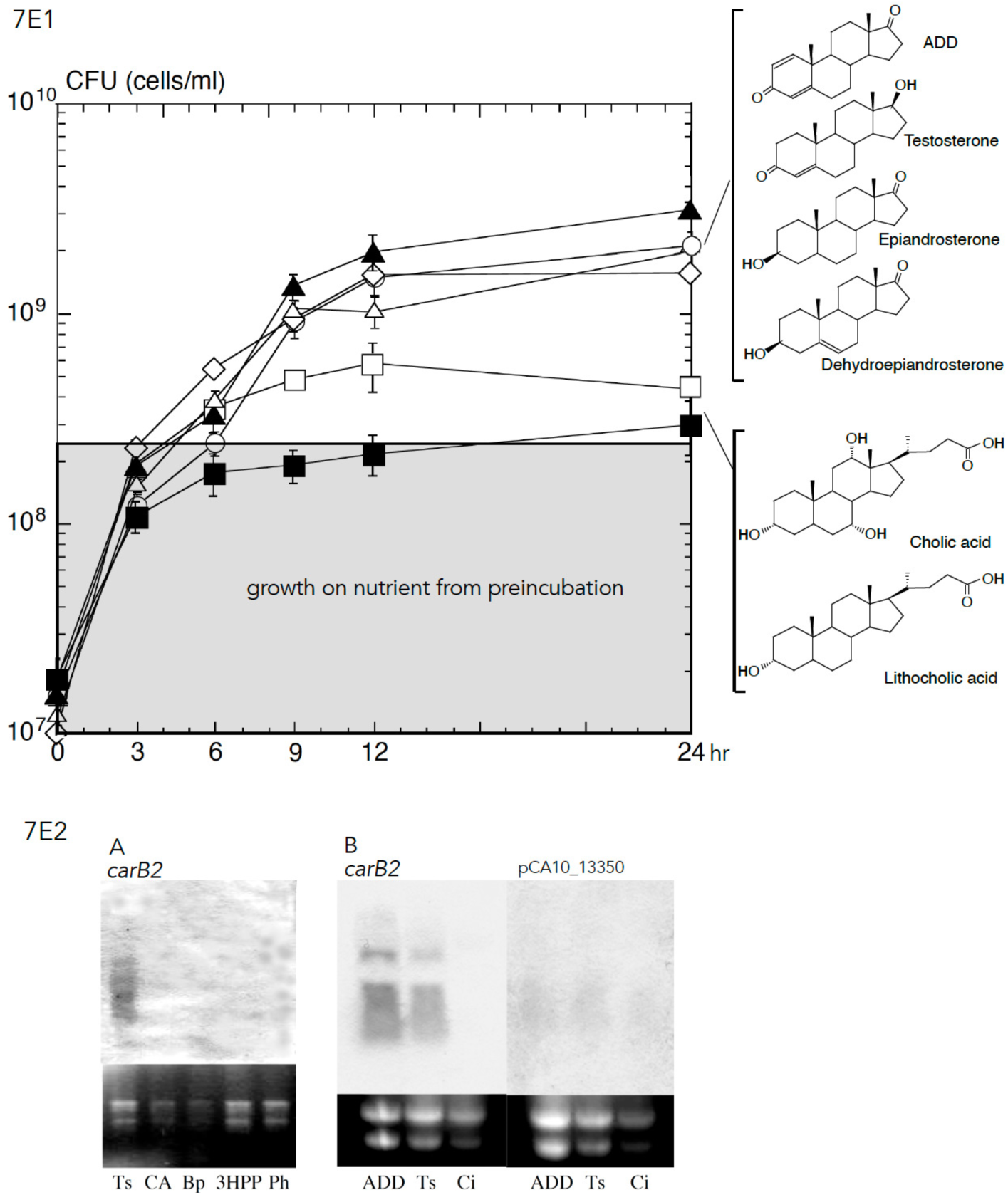
**7A-7D2:** Comparison of steroid degradation gene clusters using CAGECAT. The query sequences and search targets used for the analysis are summarized in Table 10. *: strains with complete genome sequences. The genes involved in A- and B-ring cleavage (*tes* genes) are shown in dark blue (*tesHIJ*) and dark red (*tesA1A2DEFG*); the CoA transferase acting after A- and B-ring cleavage (*scdA*) in pink; genes involved in B-, C-, and D-ring degradation (*scd* genes) in light blue; and *chs* genes in orange. The colors of genes outside the boxes do not necessarily indicate their functions and are used to indicate similar genes among the strains. Identities between the TA441 enzymes and the corresponding enzymes in other strains are listed in Table 7. **7A:** Comparison of steroid degradation gene clusters in TA441 and *Pseudomonas stutzeri* Chol1 with the degradation pathway of the C17 side chain of cholic acid in Chol1. **7B:** Comparison of steroid degradation gene clusters in γ-Proteobacteria. **7C:** Comparison of the genes corresponding to *tesB–tesR* (*scd* genes) in *Pseudomonas* strains. The strains whose names are shown in blue are presented in Fig. 6A, whereas those whose names are shown in pink are presented in Fig. 7B. **7D1:** Comparison of steroid degradation gene clusters in TA441 and *Metapseudomonas resinovorans* NBRC 106553. **7D2:** Comparison of the genes corresponding to *tesB–tesR* (*scd* genes) in *Metapseudomonas* and *Pseudomonas* strains. **7E1:** Growth of NBRC 106553 on ADD (▴), testosterone (△), epiandrosterone (○), dehydroandrosterone (◇), cholic acid (□), and lithocholic acid (▪) as the sole carbon source. The strains were preincubated in LB medium, washed, and inoculated into 10 ml of C medium (15) supplemented with 0.1% (w/v) of each steroid. Growth in initial 3 h was resulting from preincubation. **7E2A:** Induction of *carB2* (PCA10_13380) in NBRC 106553 incubated with Ts (testosterone), CA (cholic acid), Bp (biphenyl), 3HPP {3-(3-hydroxyphenyl)propionic acid}, and Ph (phenol). Total RNA was extracted from *M. resinovorans* NBRC 106553 cells cultured for 8 h in LB medium supplemented with each compound. Northern analysis was performed as described previously (16). **7E2B:** Induction of *carB2* (left panels) and a gene of locus PCA10_13350 (right panels) in NBRC 106553 incubated with ADD (1,4-androstadien-3,17-dion), Ts, and Ci (sodium citrate, negative control).

**Table 7.**
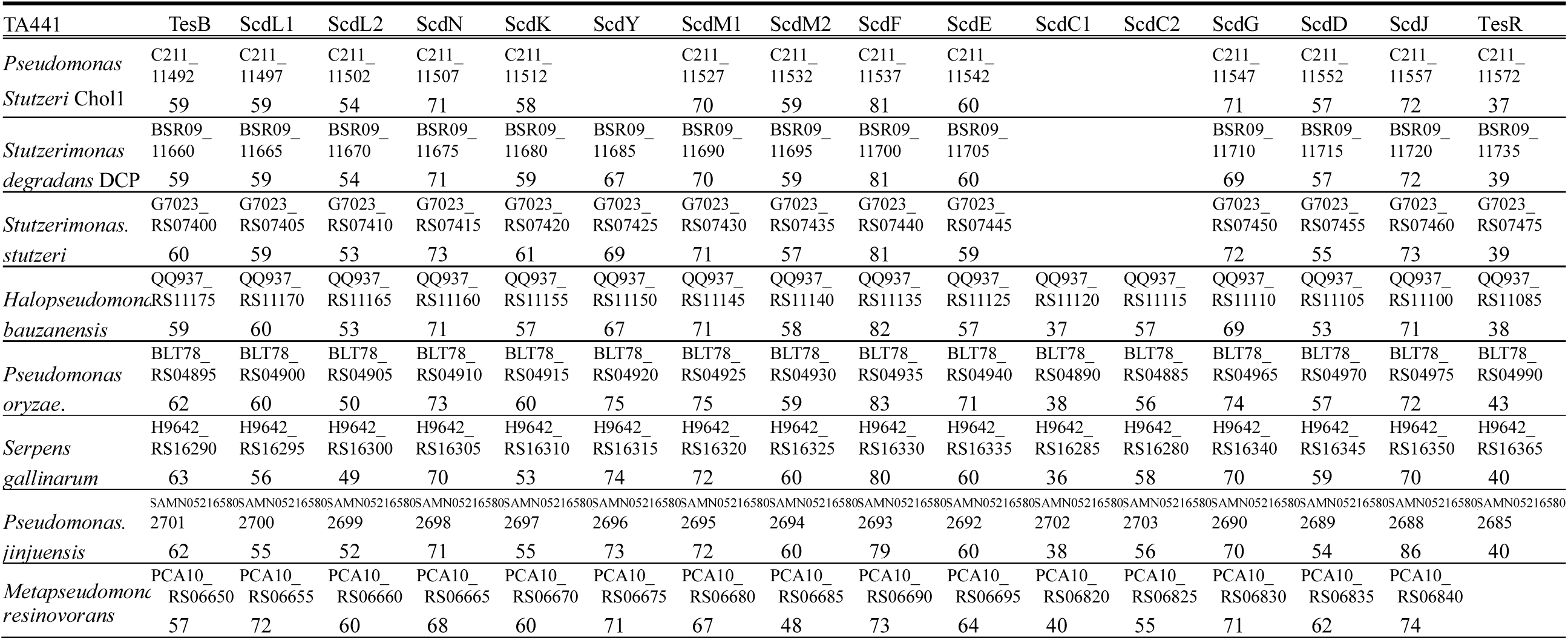
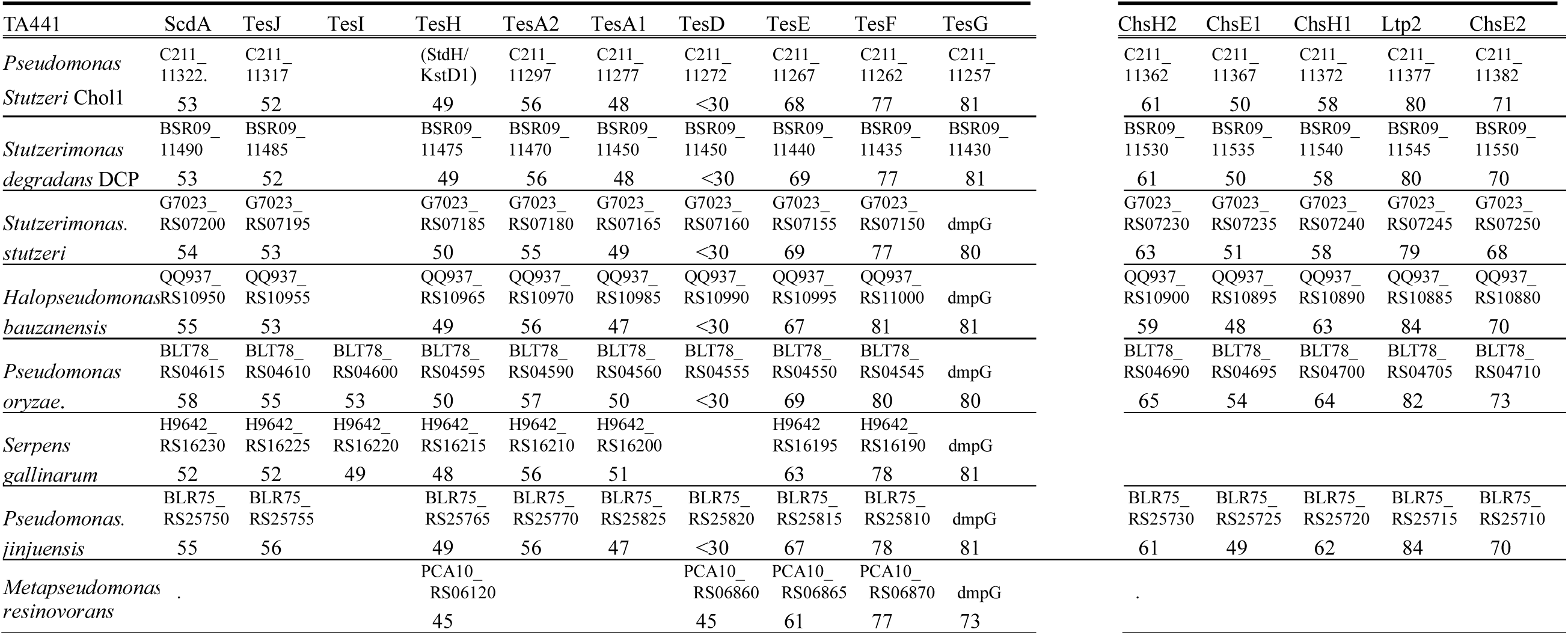
Amino acid identity (%) between the enzymes in Fig. 7BC (locus tag or gene name) and corresponding enzymes in TA441.

**Table 8.**
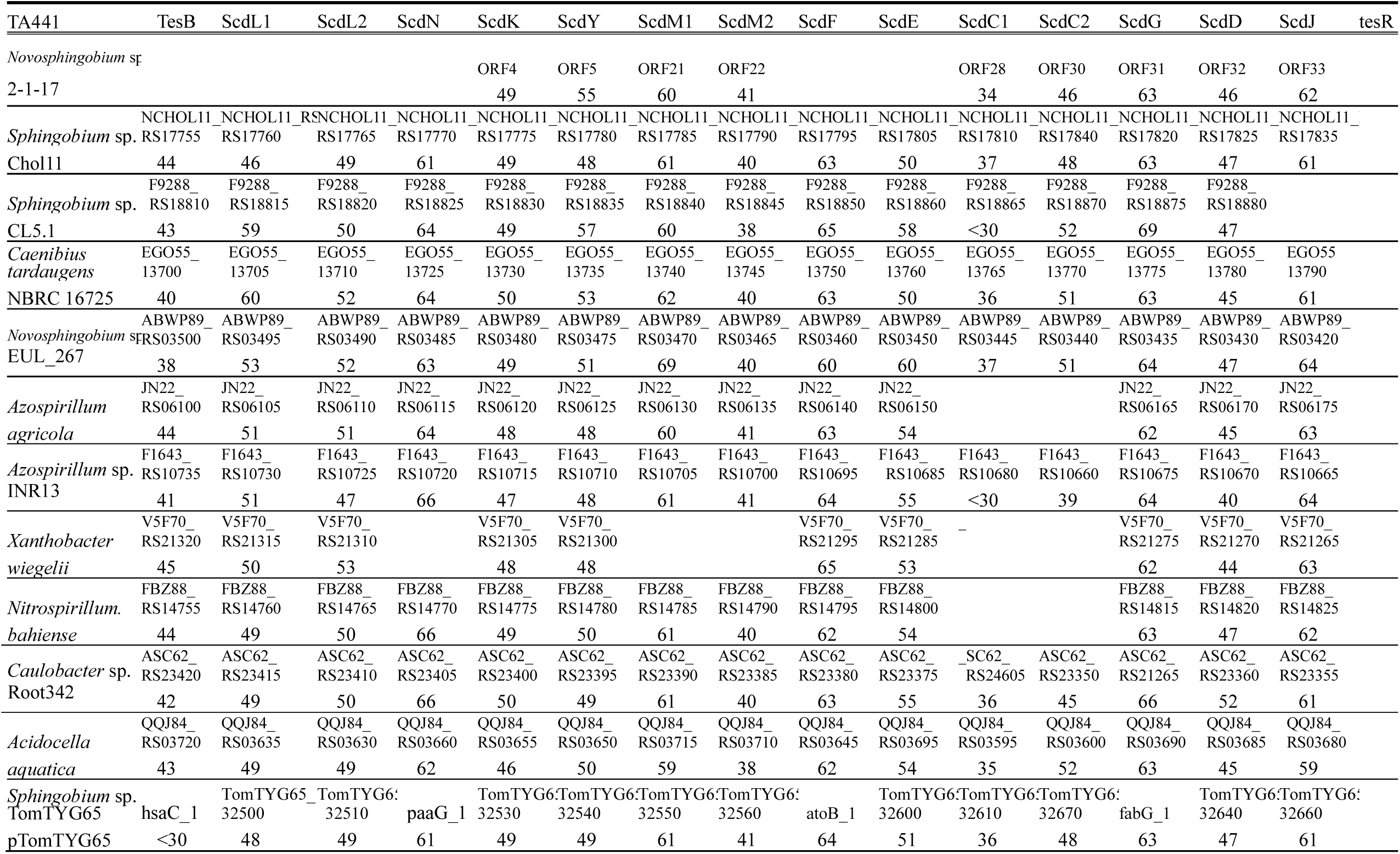
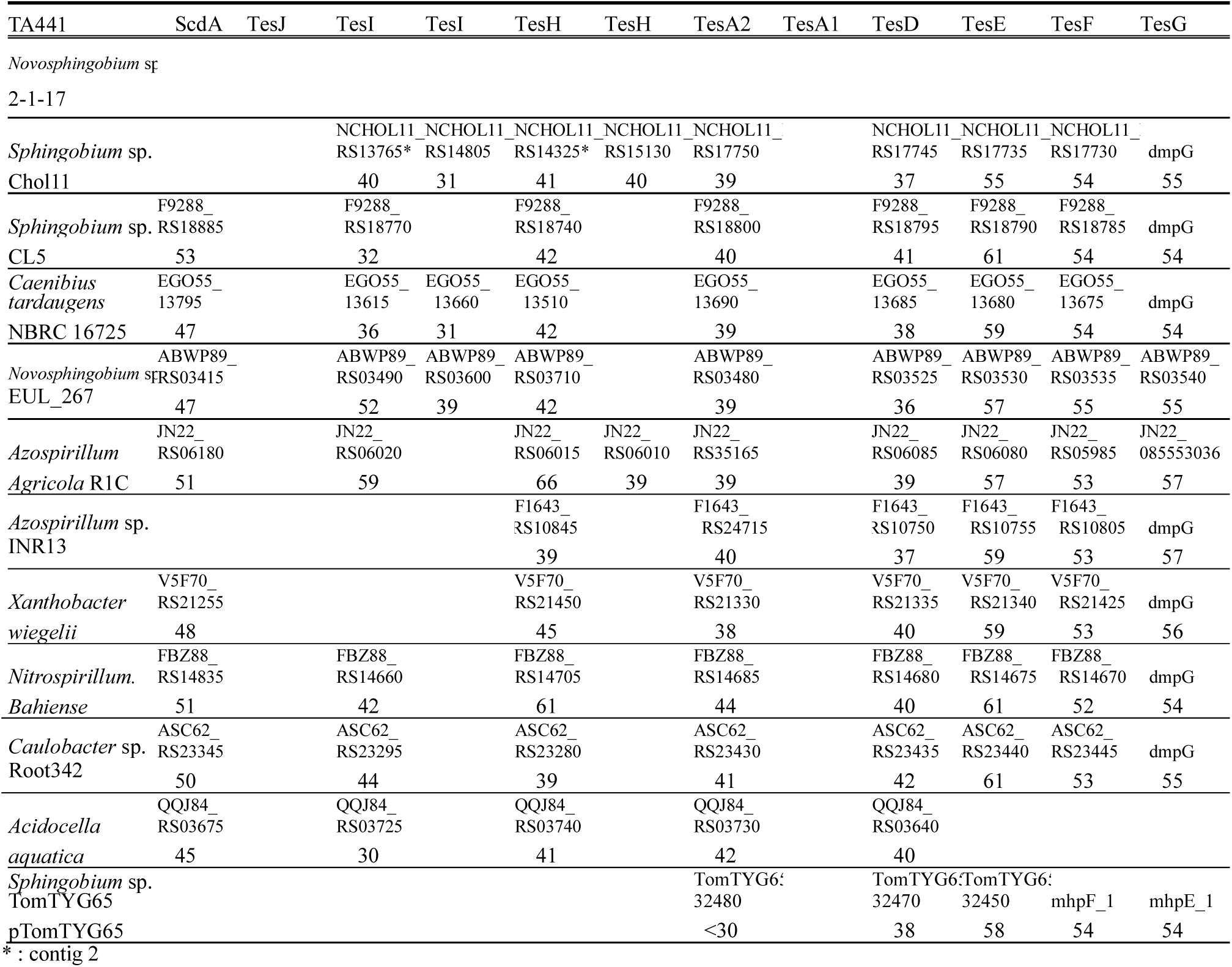
Amino acid identity (%) between the enzymes in Fig. 8B (locus tag or gene name) and corresponding enzymes in TA441.

**Table 9.** Amino acid identity (%) between the enzymes in Fig. 9B (locus tag or gene name) and corresponding enzymes in TA441.

| TA441 | TesB | ScdL1 | ScdL2 | ScdN | ScdK | ScdY | ScdM1 | ScdM2 | ScdF | ScdE | ScdC1 | ScdC2 | ScdG | ScdD | ScdJ |
| --- | --- | --- | --- | --- | --- | --- | --- | --- | --- | --- | --- | --- | --- | --- | --- |
| *Mycobacterium tuberculosis* | hsaC | ipdA*<br>Rv3551 | ipdB*<br>Rv3552 |  | ipdC*<br>Rv3553 | echA20* | ipdE3<br>fadE31 | ipdE4<br>fadE32 | fadA5? /43 | Rv3548c? | ipdE2<br>fadE33 | ipdE1<br>fadE30 | ipdF*<br>Rv3559c |  | fadA6* |
| H37Rv | 42 | 40 | 37 |  | 45 | 43 | 50 | <30 | fadA6? /60 | 54 | 31 | 52 | 61 |  | 37 |
| *Rhodococcus jostii* | hsaC |  |  | RHA1_<br>ro04594 |  |  | RHA1_<br>ro04593 | RHA1_<br>ro04592 |  |  | RHA1_<br>ro04591 | RHA1_<br>ro04596 | RHA1_<br>ro04597 |  | RHA1_<br>ro04599 |
| RHA1 | 44 |  |  | 62 |  |  | 53 | 32 |  |  | 32 | 51 | 62 |  | <30 |
| *Prescottella equi* | hsaC | HMPREF07<br>RS12560 | HMPREF07<br>RS12565 | HMPREF07<br>RS12640 | HMPREF07<br>RS12570 | echA20 | HMPREF07<br>RS12645 | HMPREF07<br>RS12650 | HMPREF07<br>RS12615 | HMPREF07<br>RS12545 | HMPREF07<br>RS12655 | HMPREF07<br>RS12630 | HMPREF07<br>RS12625 |  | HMPREF07<br>RS12615 |
| ATCC 33707 | 42 | 42 | 36 | 58 | 44 | 42 | 54 | 30 | 62 | 55 | 35 | 54 | 62 |  | <30 |
| *Tomitella biformata* | hsaC | JJ684_<br>RS00215 | JJ684_<br>RS00210 | JJ684_<br>RS00115 | JJ684_<br>RS00200 | echA20 | JJ684_<br>RS00110 | JJ684_<br>RS00105 | JJ684_<br>RS00155 | JJ684_<br>RS00225 | JJ684_<br>RS00100 | JJ684_<br>RS00125 | JJ684_<br>RS00130 |  | JJ684_<br>RS00240? 36 |
| DSM45403 | <30 | 40 | 37 | 60 | 43 | 42 | 52 | 31 | 68 | 52 | 35 | 54 | 59 |  | RS00150? <30 |
| *Antrihabitans* sp. | hsaC | JGU72_<br>RS14430 | JGU72_<br>RS14435 | JGU72_<br>RS14495 | JGU72_<br>RS14440 | JGU72_<br>RS14425 | JGU72_<br>RS14500 | JGU72_<br>RS14505 | JGU72_<br>RS14470 | JGU72_<br>RS14415 | JGU72_<br>RS14510 | JGU72_<br>RS14485 | JGU72_<br>RS14480 |  | JGU72_<br>RS14470 |
| YC2-6 | 42 | 43 | 35 | 59 | 43 | 47 | 55 | 33 | 61 | 57 | 31 | 53 | 60 |  | <30 |

\*: The substrate in *M. tuberculosis* and that in TA441 is different

| TA441 | ScdA | TesJ | TesI | TesH | TesA2 | TesA1 | TesD | TesE | TesF | TesG |
| --- | --- | --- | --- | --- | --- | --- | --- | --- | --- | --- |
| *Mycobacterium tuberculosis* | fadD3 | kshB |  | kshD | hsaA |  | hsaD_ | hsaE | hsaG | hsaF |
| H37Rv | 45 | 44 |  | 37 | 36 |  | 33 | 42 | 56 | 49 |
| *Rhodococcus jostii* | RHA1_<br>ro04595 |  |  | kshD | hsaA |  | hsaD_ | hsaE | hsaG | hsaF |
| RHA1 | 45 |  |  | 39 | 37 |  | 33 | 40 | 54 | 49 |
| *Prescottella equi* | HMPREF07<br>RS12635 | kshB |  | kshD | hsaA |  | hsaD |  |  |  |
| ATCC 33707 | 44 | <30 |  | 38 | 37 |  | 32 |  |  |  |
| *Tomitella biformata* | JJ684_<br>RS00120 | JJ684_<br>RS00045 | JJ684_<br>RS00025 | kshD | hsaA |  | hsaD |  |  |  |
| DSM45403 | 42 | 45 | 38 | 39 | 36 |  | <30 |  |  |  |
| *Antrihabitans* sp. | JGU72_<br>RS14490 |  |  | JGU72_<br>RS14595 | JGU72_<br>RS14545 |  | JGU72_<br>RS14540 | JGU72_<br>RS14590 | JGU72_<br>RS14585 | JGU72_<br>RS14580 |
| YC2-6 | 45 |  |  | 39 | 37 |  | 33 | 39 | 57 | <30 |

| ChsH2 | ChsE1 | ChsH1 | Ltp2 | ChsE2 |
| --- | --- | --- | --- | --- |
| Rv3542c | fadE28 | Rv3541c | ltp2 | fadE29 |
| 34 | <30 | 38 | 62 | <30 |
| RHA1_<br>ro04486 | RHA1_<br>ro04484 | RHA1_<br>ro04487 | RHA1_<br>ro04487 | RHA1_<br>ro04485 |
| 34 | <30 | 36 | 61 | 46 |
| HMPREF07 | HMPREF07 | HMPREF07 | HMPREF07 | HMPREF07 |
| RS12875 | RS12885 | RS12870 | RS12865 | RS12880 |
| 34 | <30 | 38 | 62 | <30 |
| JGU72_<br>RS14640 | JGU72_<br>RS14650 | JGU72_<br>RS14635 | JGU72_<br>284337730 | JGU72_<br>RS14645 |
| 38 | <30 | 36 | 63 | <30 |

**Table 10.** Blast analysis query and target.

| Figure | Query | BLAST search | Target |
| --- | --- | --- | --- |
| Fig. 5 | *tesB-tesR* * | BLASTn | *Comamonas* |
| Fig. 5 | TesH, ScdL1, ScdY* | BLASTp | *Comamonas* |
| Fig. 6A | *tesB-tesR, scdA-tesG* * | BLASTn | non-*Comamonas* β-Proteobacteria |
| Fig. 6B | TesH, ScdL1, ScdY* | BLASTp | non-*Comamonas* β-Proteobacteria |
| Fig. 7B | StdH (Chol1) | BLASTp | γ-Proteobacteria |
| Fig. 7C | ScdL1-like protein (Chol1) | BLASTp | *Pseudomonas* |
| Fig. 7D2 | ScdL1-like protein (NBRC 106553) | BLASTp | *Pseudomonas* / *Metapseudomonas* |
| Fig. 8B | TesH* | BLASTp | α-Proteobacteria |
| Fig. 9B | KstD (H37Rv) | BLASTp | Actinomycetota |

\*: TA441
Chol1: *Pseudomonas stutzeri* Chol1NBRC 106553: *Metapseudomonas resinovorans* NBRC 106553H37Rv: *Mycobacterium tuberculosis* H37Rv

Deletion of one of the duplicated sequences (“GGGCAACGGGGCAACG” to “GGGCAACG”) resulted in an enzyme (StdH/KstD1) showing 49% amino acid identity to TesH of TA441.

In *P. stutzeri* Chol1, degradation of the C17 side chain of cholic acid has been characterized in detail (Fig. 7A, degradation pathway). Genes involved in C17 side-chain degradation appear to form a large cluster together with genes involved in A- and B-ring cleavage and B-, C-, and D-ring degradation in Chol1 (Fig. 7A). The 3α-DH and 3β-DH genes are also included in this cluster, whereas the 3β-DH gene is located outside the mega-cluster in TA441. The locations of genes involved in B-, C-, and D-ring degradation were quite similar to those of the *scd* genes, except that *scdC1C2*-like genes were missing and two ORFs (locus tags C211_11562 and C211_11567) were inserted between genes corresponding to *scdJ* and *tesR*.

For γ-Proteobacteria, the amino acid sequence of StdH (KstD1) from Chol1 was used for a BLAST search, and DNA sequences containing putative steroid degradation genes were obtained and analyzed using CAGECAT in the same manner as described above. First, we obtained the sequence of Δ1-dehydrogenase from a strain of the same species, *Stutzerimonas degradans* DCP-Ps1 (86) (Chol1 was *S. degradans* for a while before reclassified to *P. stutzeri*), and compared it with the modified StdH (KstD1) sequence to confirm that the modification was correct. The amino acid identities between these enzymes were 99%. *S. stutzeri* 99A1 (87), *Halopseudomonas bauzanensis* AHM20C89I (88), *P. oryzae* KCTC 32247 (89), *Serpens gallinarum* Sa2CUA2 (90), and *P. jinjuensis* JCM 21621 (91) possessed putative A- and B-ring cleavage gene clusters, B-, C-, and D-ring degradation gene clusters, and genes involved in C17 side-chain degradation (Fig. 7B, identities between the corresponding enzymes and those of TA441 are listed in Table 7). All except *Serpens gallinarum* possessed Chol1-type A- and B-ring cleavage gene clusters, including *sad* and 3β-DH, and B-, C-, and D-ring degradation gene clusters characterized by the insertion of two ORFs between genes corresponding to *scdJ* and *tesR*. Genes corresponding to *scdC1C2* were found in the DNA region upstream of *tedB*-like gene in *P. oryzae*, *Serpens gallinarum*, and *P. jinjuensis* and in the same location as TA441 in *H. bauzanensis*.

The genes corresponding to *tesB* to *tesR* in the *Pseudomonas* strains shown in Fig. 7B differed from those shown in Fig. 6A. We therefore analyzed several other *Pseudomonas* strains, *P. putida* W5 (92), *Pseudomonas* sp. AN-1 (93), and *P. guangdongensis* CCTCC 2012022 (94), whose putative steroid degradation genes showed relatively high similarity to the corresponding genes of TA441 (Fig. 7C). As a result, all the examined *Pseudomonas* strains were found to possess different putative steroid degradation gene clusters but *scd* genes of *Pseudomonas* sp. AN-1 and *P. guangdongensis* CCTCC 2012022 were similar to those of *P. oryzae* KCTC 32247 and *P. jinjuensis* JCM 21621 in Fig. 7B.

When TesB in TA441 was isolated and identified in 1999 (published in 2001) as the first aromatized A-ring cleavage enzyme identified in bacterial steroid degradation (15), its amino acid sequence showed the highest homology to CarB2 (locus tag PCA10_13380), an aromatic ring-cleavage enzyme of unknown substrate in the well-known carbazole-degrading bacterium *Metapseudomonas resinovorans* NBRC 106553 (formerly *Pseudomonas resinovorans* CA10) (95). We isolated an approximately 20-kb DNA region containing *carB2* and found genes corresponding to *scdL1* to *scdE* in the same order as in TA441 (Fig. 7D1). However, we were unable to isolate genes corresponding to *steA*–*steD*, which are necessary for cholic acid degradation and are encoded in the DNA region immediately upstream of *tesB*, as well as other putative steroid degradation genes.

A complete genome sequence of *M. resinovorans* NBRC 106553 is now available, and we therefore analyzed it using CAGECAT as described above (Fig. 7D1). Genes corresponding to *scdC1*–*scdJ* in NBRC 106553 were found approximately 25 kb apart from the *scdE*-like gene. The functions of the genes in this 25-kb DNA region are unclear. *Pseudomonas* sp. AN-1 in Fig. 7C also has a long insertion of approximately 23 kb in its B-, C-, and D-ring degradation gene cluster. The Δ1-dehydrogenase gene *kstD*, corresponding to *tesH*, was found at locus tag PCA10_12340 together with genes similar to ORF50 and ORF51 of TA441, whose functions are unclear. Other possible steroid degradation genes were not found in the DNA region near *kstD* in NBRC 106553 or in the 25-kb DNA region within the B-, C-, and D-ring degradation gene cluster.

Among the *Pseudomonas* strains examined in Fig. 7C and NBRC 106553, the insertion in the putative *scd* clusters varied considerably. We therefore further examined *Pseudomonas* and *Metapseudomonas* strains for genes homologous to the NBRC 106553 ScdL1-like enzyme (locus tag PCA10_13390) to determine whether the long insertion observed in NBRC 106553 was also present in other strains. Six strains showing higher homology—*M. furukawaii* UC4535 (96), *Pseudomonas* sp. MOB-449 (97), *Pseudomonas* sp. TCU-HL1 (98), *M. lalkuanensis* ACYW.190 (99), *Pseudomonas* sp. LFM046 (100), and *M. resinovorans* NBRC 113413 (101)—were selected. Analysis of the DNA regions surrounding the *scdL1*-like genes in these six strains showed that all had insertions at the same location as the insertion in the NBRC 106553 cluster, although the genes within these regions differed (Fig. 7D2). The genes in *M. furukawaii* UC4535 and *Pseudomonas* sp. MOB-449 were similar, and the total length of the inserted regions was only approximately 2 kb, whereas that of *M. resinovorans* NBRC 113413, the longest among the six strains, was approximately 17 kb and was completely different from that of NBRC 106553. In our analysis of *Pseudomonas* and *Metapseudomonas* strains, NBRC 106553 had the longest insertion.

In contrast, the genes in the region from the *tesB*-like gene to the *scdE*-like gene (locus tags PCA10_13380–PCA10_13470 in NBRC 106553; region 1 in Fig. 7D2) and from the *scdC1*-like gene to the *tesG*-like gene (locus tags PCA10_13730–PCA10_13840 in NBRC 106553; region 2 in Fig. 7D2) were exactly the same as those in NBRC 106553 in all six strains.

NBRC 106553 showed growth on steroids with a hydroxyl or ketone group at C17, such as testosterone and androsterone, but did not grow on cholic acid or lithocholic acid (Fig. 7E1). Northern blot analysis showed that *carB2* was induced when NBRC 106553 was grown with testosterone but not with cholic acid or other aromatic compounds, including biphenyl, 3HPP, and phenol (Fig. 7E2A). An ORF (locus tag PCA10_13350), located in the gene region corresponding to the region containing genes involved in removal of the C12 hydroxyl group in TA441, was not induced by 1,4-androstadien-3,17-dione (ADD) or testosterone (Fig. 7E2B). The gene structure, the growth, and the induction were consistent.

Amino acid identities between TesH of TA441 and the corresponding enzymes in these γ-Proteobacteria were approximately 45–50% (Table 7).

### Comparison of the steroid degradation gene clusters V: α-Proteobacteria (**Fig. 8**; identities are shown in **Table 8**)

Along with the analysis of steroid degradation in TA441, we also conducted studies on the isolation and characterization of estrogen-degrading bacteria. At that time, estrogen was regarded as one of the “endocrine-disrupting chemicals” of environmental concern. We isolated approximately 10 estrogen-degrading bacteria, most of which were Actinomyces. The estrogen degradation pathway of one of the isolates, *Rhodococcus* sp. B50, was elucidated recently (102–104). Estrogen has an aromatic A-ring, and cleavage of the A-ring occurs first during its degradation. The cleaved A-ring is removed by β-oxidation, followed by hydrolysis of the B-ring to produce 9,17-DOHNA, the same intermediate compound produced during bacterial aerobic degradation of testosterone and lithocholic acid.

**Fig. 8.**
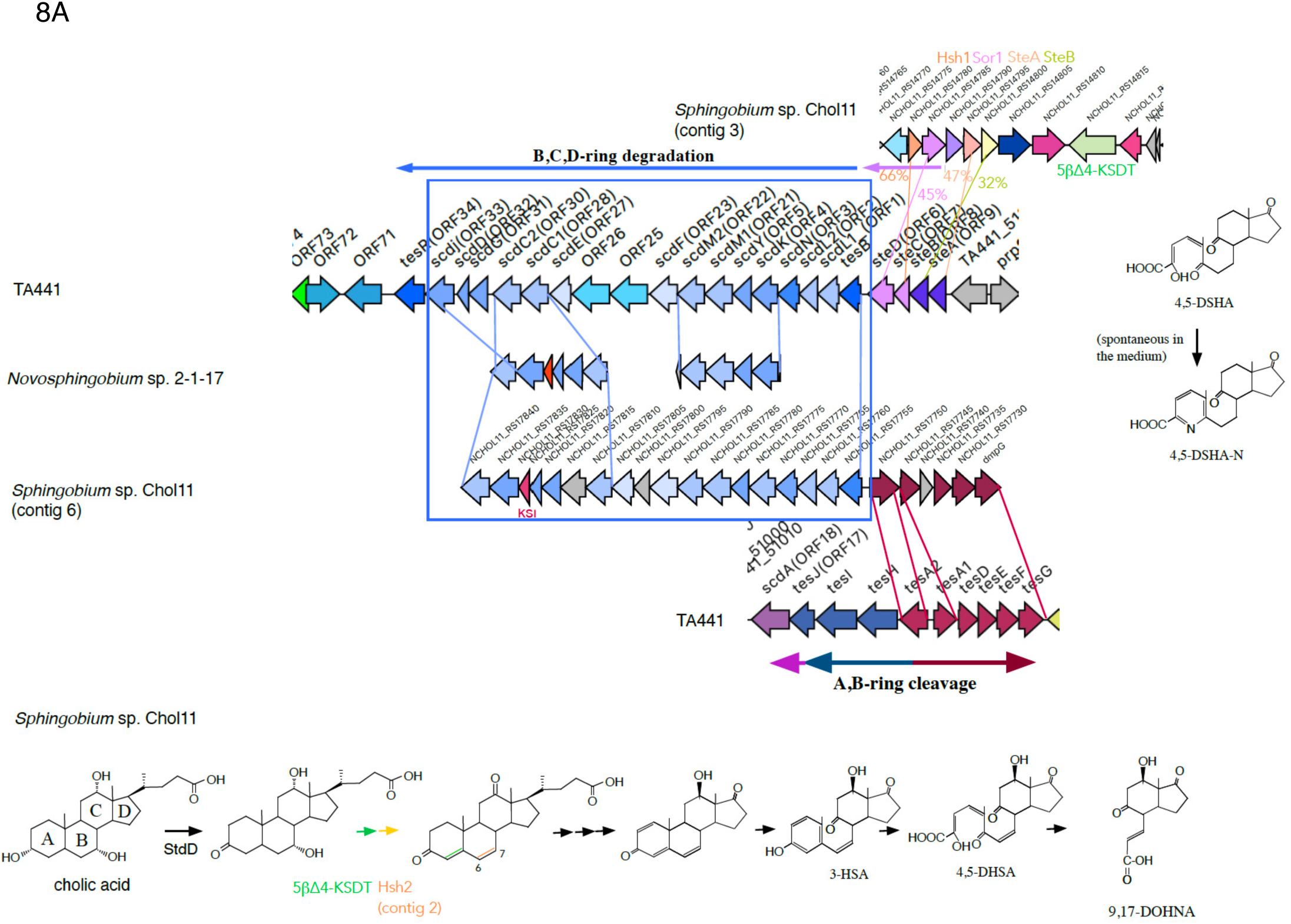
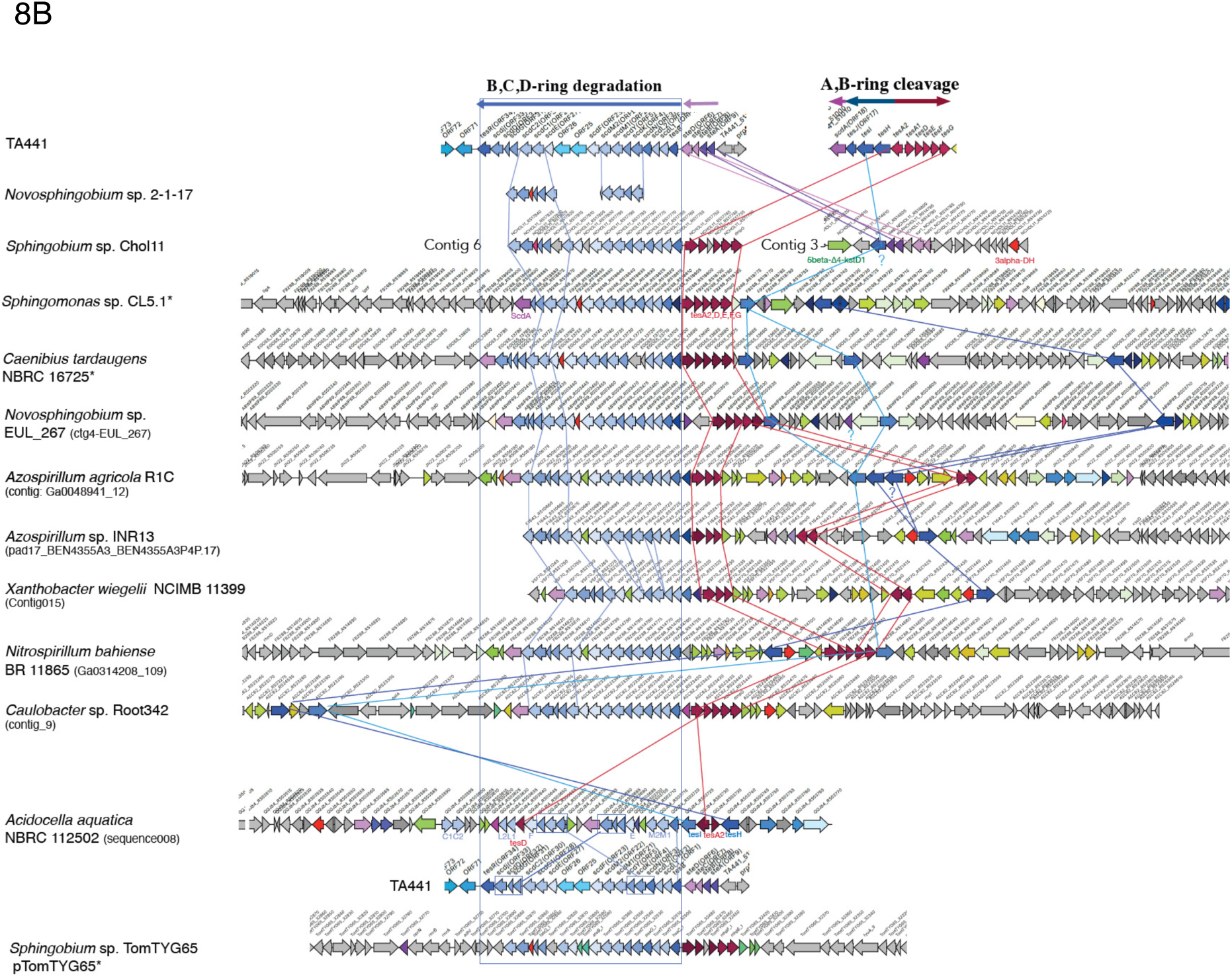
Comparison of the steroid degradation gene clusters using CAGECAT. The genes involved in A- and B-ring cleavage (*tes* genes) are shown in dark blue (*tesHIJ*) and dark red (*tesA1A2DEFG*); the CoA transferase acting after A- and B-ring cleavage (*scdA*) in pink; genes involved in B-, C-, and D-ring degradation (*scd* genes) in light blue; and *chs* genes in orange. The colors of genes outside the boxes do not necessarily indicate their functions and are used to indicate similar genes among the strains. **8A:** Comparison of the steroid degradation gene clusters of TA441 and *Sphingobium* sp. Chol11, together with the major intermediates of the cholic acid Δ4,6 degradation pathway in Chol11. **8B:** Comparison of the steroid degradation gene clusters of α-Proteobacteria. Identities between the TA441 enzymes and the corresponding enzymes in other strains are listed in Table 8.

Among the isolated estrogen degraders, *Novosphingomonas* sp. strain 2-1-17 was the only α-Proteobacterium. Strain 2-1-17 degraded both estrogen and cholic acid. Detection of a derivative of 4,5-DSHA-N (stable derivative of 4,5-DHSA spontaneously produced in the culture, Fig. 8A) in cultures of strain 2-1-17 incubated with testosterone suggested that its A- and B-ring cleavage process was similar to that of TA441. In addition, the partially determined DNA sequence suggested the presence of a gene cluster similar to that involved in B-, C-, and D-ring degradation in TA441 (*scdN*–*scdM2* and *scdC1*–*scdJ*, Fig. 8A).

We did not continue further studies of strain 2-1-17, but cholic acid degradation in α-Proteobacteria was subsequently characterized in *Sphingobium* sp. Chol11 (formerly *Novosphingobium* sp. Chol11) (105–108). Chol11 cleaves the A- and B-rings via aromatization of the A-ring, but the C6–C7 double bond is retained throughout the process. The genes involved in B-, C-, and D-ring degradation in Chol11 were found on contig 6 of the shotgun genome sequence (107) (Fig. 8A). The genes in this cluster are arranged in the same order as those in TA441, except that the *scdC2*-like gene is located after the *scdJ*-like gene and *ksi* is inserted between the *scdD*-like and *scdJ*-like genes. Adjacent to the *tesB*-like gene, genes corresponding to *tesA2*–*tesG* of TA441 are also encoded. Genes corresponding to *steA*–*steD*, the 5β-Δ4-KstD1 gene, and the 3α-DH gene form a cluster on contig 3 (Fig. 8A, upper panel and Fig. 8B) (105). All these genes are necessary for cholic acid degradation. The gene encoding the dehydrogenase responsible for introducing a double bond at C6–C7 (*hsh2*) is located on contig 2, whereas the gene corresponding to *tesH* has not been identified.

Therefore, to examine steroid degradation genes in α-Proteobacteria, we searched for possible TesH-like enzymes using the TesH sequence, limiting the search to α-Proteobacteria, and compared the surrounding 200-kb DNA regions containing the identified putative TesH-like genes using CAGECAT as described above (Fig. 8B; identities between the corresponding enzymes and those of TA441 are listed in in Table 8). Although the genes surrounding the putative TesH-like enzymes differed considerably among the strains examined, genes involved in A- and B-ring cleavage and B-, C-, and D-ring degradation were similar. A cluster corresponding to *tesB*–*tesJ*, with *tesA2*–*tesG*-like genes located nearby, was found in *Sphingomonas* sp. CL5.1 (109),

*Caenibius tardaugens* NBRC 16725 (105, 110), *Novosphingobium* sp. EUL_267 (111), *Azospirillum agricola* R1C (112), *Azospirillum* sp. INR13 (113), *Xanthobacter wiegelii* NCIMB 11399 (114), *Nitrospirillum bahiense* BR 11865(115), *Caulobacter* sp. Root342 (116), *Acidocella aquatica* NBRC 11250 (61), and in the mega plasmid pTomTYG65 in *Sphingomonas* sp. TomTYG65 (117).

*Sphingomonas* sp. TomTYG65 degrades α-tomatine, a steroid-type saponin secreted from the roots of tomato (*Solanum lycopersicum*). The cluster on pTomTYG65 was the only steroid degradation gene cluster found on a plasmid in this study. Genes corresponding to *tesH* and *tesI* are located separately from the cluster.

*Acidocella aquatica* NBRC 11250 has a unique putative steroid degradation gene cluster (Fig. 8B), but a BLAST search using the DNA sequence of the entire putative steroid degradation gene cluster of NBRC 11250 did not identify similar sequences. Amino acid identities between TesH of TA441 and the corresponding enzymes in these α-Proteobacteria were approximately 39–42%, except for *A. agricola* R1C (66%) and *Nitrospirillum bahiense* BR 11865 (61%) (Table 8).

We also conducted BLAST searches and CAGECAT analyses of δ-, ε-, and ζ-

Proteobacteria in the same manner, but identified only enzymes showing less than 30% amino acid identity, with no putative steroid degradation genes detected in the surrounding DNA regions.

### Comparison of the steroid degradation gene clusters VI: Actinomycetota (**Fig. 9**, identities are in **table 9**)

Actinomycetota *Rhodococcus hoagii* (= *Rhodococcus equi*, formerly *Nocardia restricta*) and the proteobacterium *Comamonas testosteroni* (formerly *Pseudomonas testosteroni*) have been studied as representative aerobic steroid-degrading bacteria since the 1950s. Major intermediate compounds in steroid degradation were identified in the 1960s, and these bacteria were shown to degrade steroids through similar processes (2, 118, 119).

**Fig. 9.**
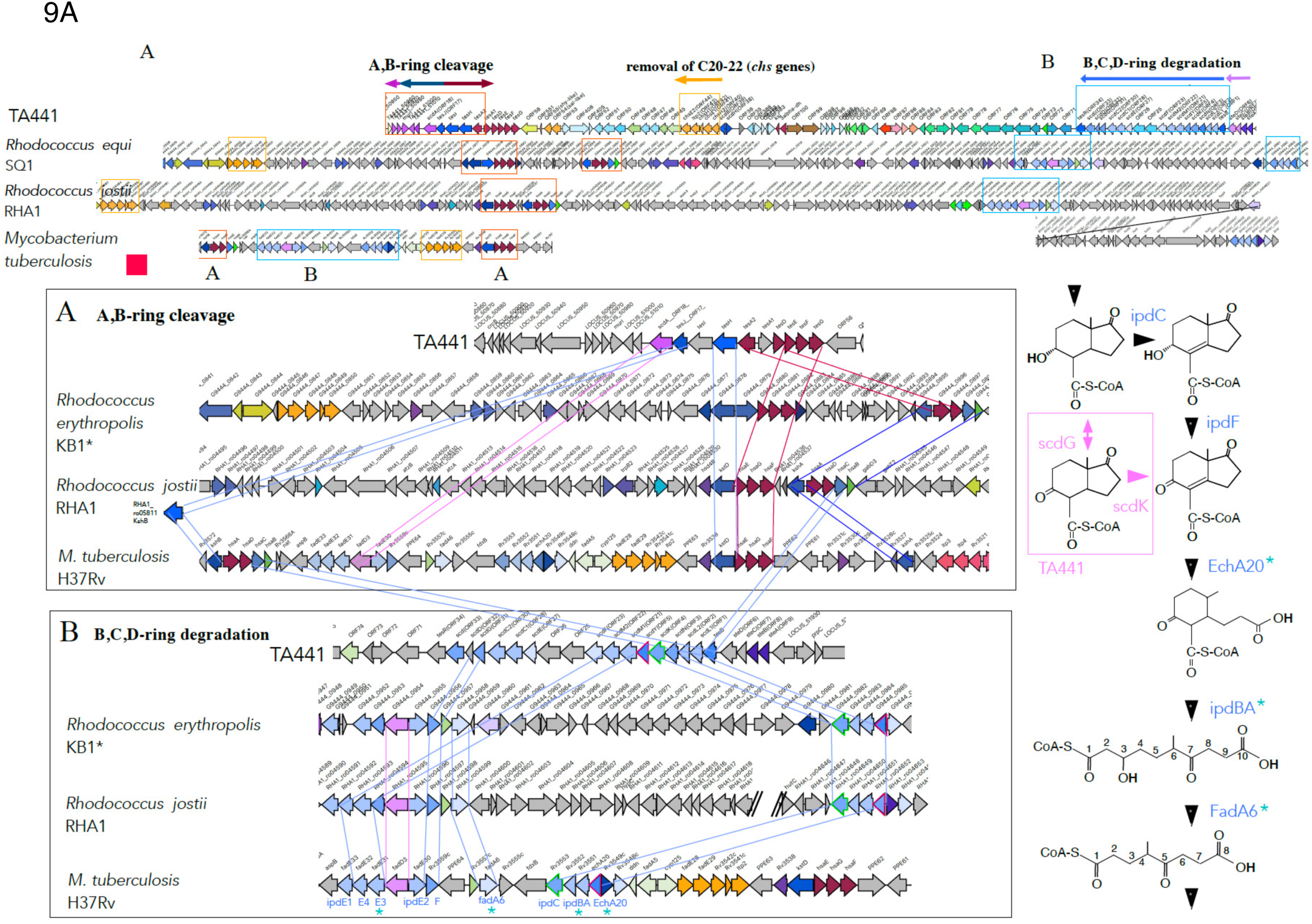
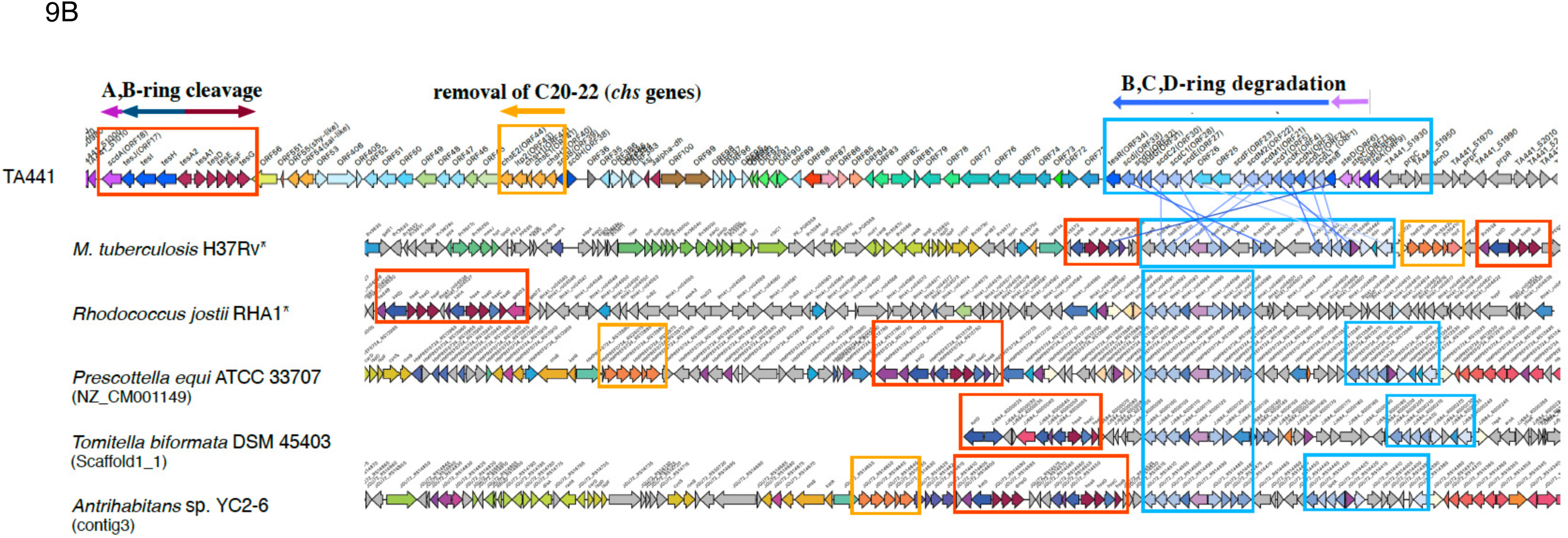
Comparison of the steroid degradation gene clusters using CAGECAT. The genes involved in A- and B-ring cleavage (*tes* genes) are shown in dark blue (*tesHIJ*) and dark red (*tesA1A2DEFG*); the CoA transferase acting after A- and B-ring cleavage (*scdA*) in pink; genes involved in B-, C-, and D-ring degradation (*scd* genes) in light blue; and *chs* genes in orange. The colors of genes outside the boxes do not necessarily indicate their functions and are used to indicate similar genes among the strains. **9A:** Comparison of the steroid degradation gene clusters of TA441, *Rhodococcus equi* SQ1, *R. jostii* RHA1, and *M. tuberculosis* H37Rv with a partial pathway for B-, C-, and D-ring degradation in Actinomycetota. ^★^: Reactions in Actinomycetota and those in TA441 are different (cf. Supplementary Materials Fig. S1). **9B:** Comparison of the steroid degradation gene clusters of Actinomycetota. Identities between the TA441 enzymes and the corresponding enzymes in other strains are listed in Table 9.

Among Actinomycetota, enzymes involved in steroid degradation have been well studied in *Rhodococcus jostii* RHA1, *R. erythropolis* SQ1, and *Mycobacterium tuberculosis* H37Rv. The Δ1-dehydrogenase gene *kstD* (10, 120), 3α-dehydrogenase gene (6), and Δ5-3-isomerase gene (121) of *Rhodococcus* were identified before we published a paper on *tesB* as the first gene encoding an A-ring cleavage enzyme in bacterial steroid degradation in 2001. After we revealed the complete set of A- and B-ring cleavage genes in TA441 (15–20, 33, 37), Δ1-dehydrogenases in *Mycobacterium* (122–125) and other genes involved in A- and B-ring cleavage were also identified in *Rhodococcus* and *Mycobacterium* (14, 126–128). These studies revealed that Proteobacteria and Actinomycetota not only share similar A- and B-ring cleavage pathways, but also have similar enzymes involved in these pathways. Enzymes responsible for removal of C20-22 from the C17 side chain were first identified in *M. tuberculosis* H37Rv (126, 129–133), and the corresponding enzymes were recently identified in TA441 (38). In addition, genes encoding enzymes involved in B-, C-, and D-ring degradation have also been reported in Actinomycetota (36). Therefore, we compared the steroid degradation gene clusters of TA441, *R. jostii* RHA1, and *M. tuberculosis* H37Rv using CAGECAT (Fig. 9A, identities between the corresponding enzymes and those of TA441 are listed in table 9).

The genes for A- and B-ring cleavage, *tesH* (*kstD* in Actinomycetota), *tesA2* (*hsaA*), and *tesDEFG* (*hsaEGF*) form a cluster in *R. jostii* RHA1, whereas they are divided into two smaller clusters located at both ends of the steroid degradation gene cluster in *M. tuberculosis* H37Rv (Fig. 9A). In contrast to the detailed studies on A- and B-ring cleavage, only one paper has described the overall B-, C-, and D-ring degradation pathway in Actinomycetota (36). Although the pathway is similar to that of TA441, there are differences in detail. FadA6 was reported to catalyze the same reaction as ScdN; however, CAGECAT indicated that FadA6 is more similar to ScdF. The substrates of IpdAB, IpdC, IpdF, and EchA20 are also different from those of the corresponding enzymes in TA441 (ScdL1L2, ScdK, ScdG, and ScdY, respectively) (the B-, C-, and D-ring degradation pathway of Actinomycetota is shown in Fig. 9A. TA441’s detailed pathway is shown in Supplementary Materials Fig. S1).

To obtain more information on steroid degradation genes in Actinomycetota, we conducted a BLAST search using the amino acid sequence of KstD from *M. tuberculosis* H37Rv, which corresponds to TesH in TA441, and compared the DNA regions containing *kstD* using CAGECAT in the same manner as described above (Fig. 9B, identities between the corresponding enzymes and those of TA441 are listed in in Table 9). Putative A- and B-ring cleavage genes were found in *Prescottella equi* ATCC 33707(134), *Tomitella biformata* AHU 1821 (DSM 45403, NBRC 106253) (135), and *Antrihabitans sp.* YC2-6 (136), in addition to *M. tuberculosis* H37Rv (137) and *R. jostii* RHA1 (138). The organization of the A- and B-ring cleavage genes differed from that in TA441, but was similar among the Actinomycetota except for *Mycobacterium*. Putative genes for removal of C20-22 from the C17 side chain, *igr* genes (*chs* genes), and genes involved in B-, C-, and D-ring degradation were also found in the nearby regions in similar gene order. The B-, C-, and D-ring degradation cluster appeared to consist of two smaller clusters, one containing *fadA5-ipdC* and the other containing *fadA6-fadE33* (blue boxes in Fig. 9B), with several genes of unknown function between them.

Amino acid identities between TesH in TA441 and the corresponding KstD enzymes in these Actinomycetota were approximately 37-39% (Table 9).

Figure 10A shows representative steroid degradation gene clusters from each bacterial group—α-, β-, and γ-Proteobacteria and Actinomycetota—analyzed in this study. The presence of these gene clusters strongly suggests the potential for steroid degradation and indicates that steroid degradation gene clusters have diverse structures among bacterial groups. These findings imply that aerobic bacterial steroid degradation may be widespread and may play an important role in the environment. Our analysis also suggests that Δ1-dehydrogenase (TesH/KstD) is a useful marker for identifying bacterial steroid degradation gene clusters.

**Fig. 10.**
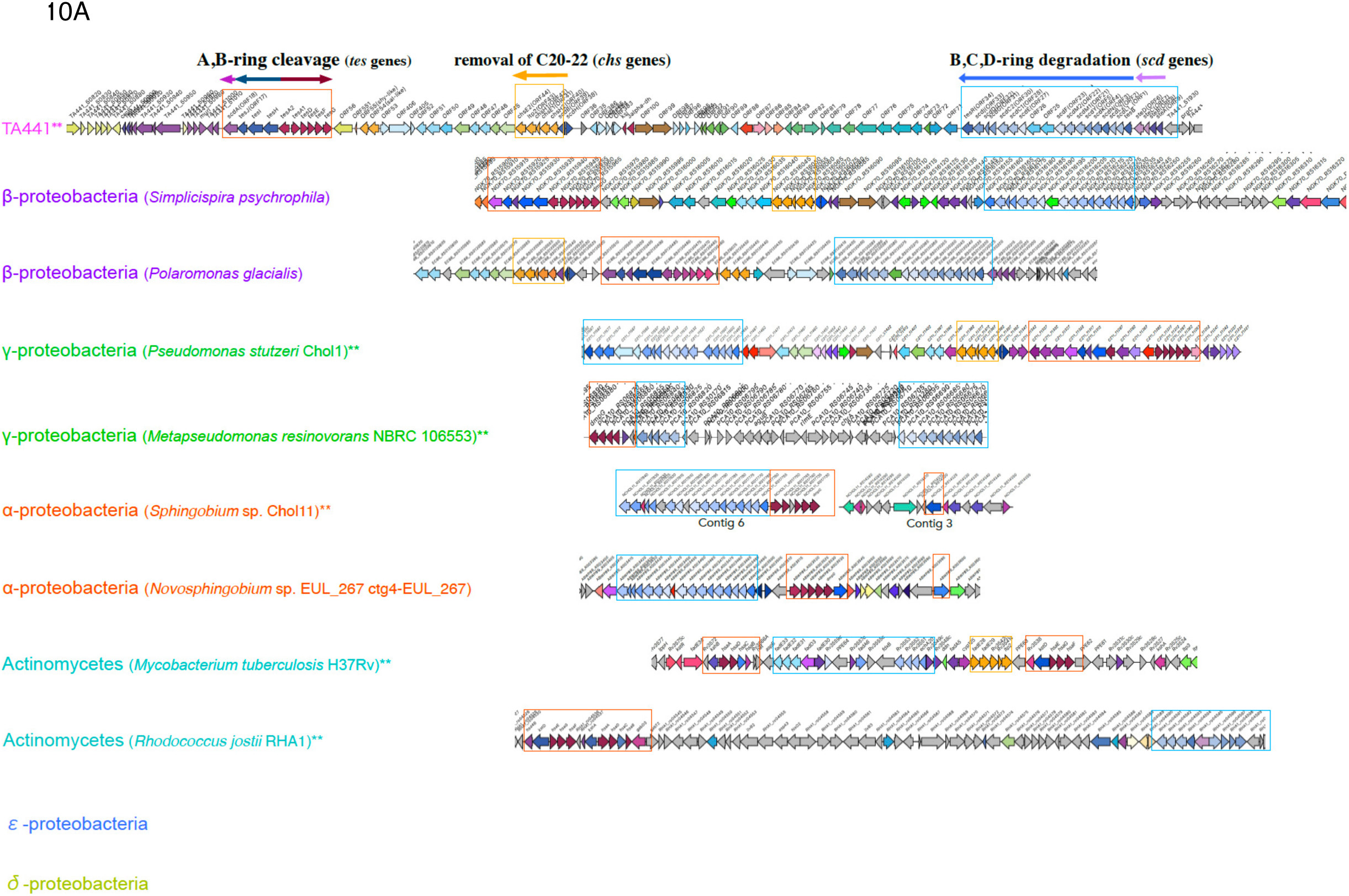
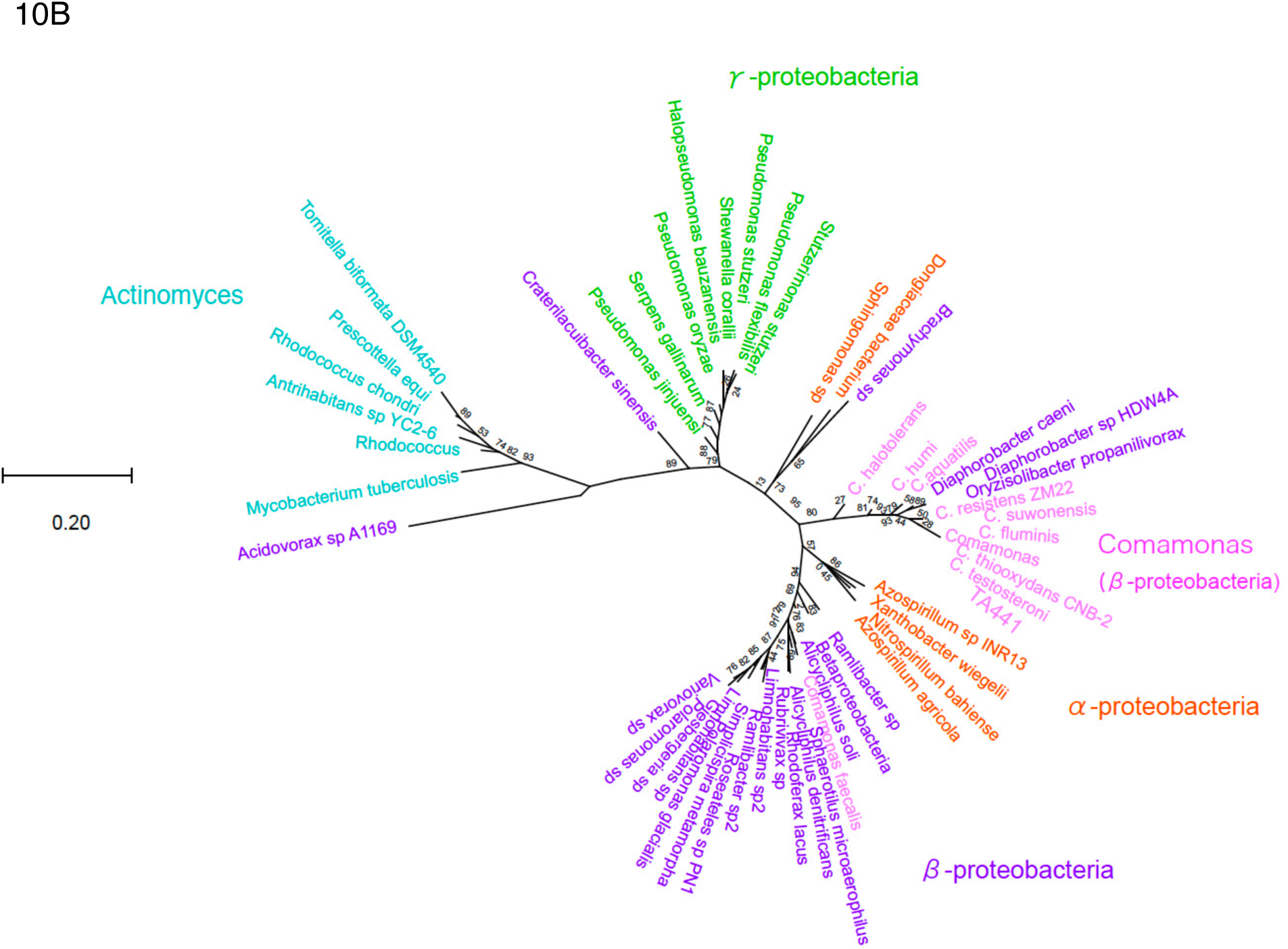
**10A:** Summary of the comparison of the steroid degradation gene clusters from two typical clusters from β-, γ-, α-Proteobacteria, and Actinomyces, respectively. Representative steroid degradation gene clusters were selected from each group. When available, strains whose steroid degradation has been experimentally studied (**) were selected; when such strains were not available, strains with characteristic putative steroid degradation gene clusters were selected. The cluster of genes involved in A- and B-ring cleavage (*tes* genes), dark blue (*tesHIJ*) and dark red (*tesA1A2DEFG*); the CoA transferase acting after A- and B-ring cleavage (*scdA*), pink; genes involved in B-, C-, and D-ring degradation (*scd* genes), light blue; and *chs* genes, orange. Color of the genes outside the boxes basically does not indicate the function and just suggests similar genes among the strains. **10B:** Phylogenetic tree based on amino acid sequences of Δ1-dehydrogenases (TesH, KstD, and StdH). *Comamonas*, pink; β-Proteobacteria except for *Comamonas*, purple; γ-Proteobacteria, green; α-Proteobacteria, red; and Actinomycetota, light blue.

A phylogenetic tree based on the amino acid sequences of Δ1-dehydrogenases from β-and γ-Proteobacteria and Actinomycetota formed distinct groups (Fig. 10B), including Δ1-dehydrogenases from bacteria whose genomic sequences are not available in the database. α-Proteobacteria did not form a distinct group, probably because they often possess several Δ1-dehydrogenase-like enzymes (cf. Fig. 8B). Compared with the 16S rRNA phylogenetic tree, Actinomycetota was positioned close to Proteobacteria, whereas *Comamonas* formed a distinct group from the other β-Proteobacteria.

## Acknowledgments

The authors sincerely thank Dr. Reizo Kato (Head of Condensed Molecular Materials Laboratory, RIKEN) and Dr. Yousoo Kim (Head of Surface and Interface Science Laboratory, RIKEN) for their thoughtful support and guidance.

## Materials and Methods

### Bacterial strain and growth conditions

*Metapseudomonas resinovorans* NBRC 106553 was used for steroid degradation experiments. Cells were grown at 30°C in LB medium for approximately 15 h. Cells were collected by centrifugation and resuspended in C medium (15) to an optical density at 600 nm (OD600) of 10. An aliquot (50 μl) of the cell suspension was inoculated into 10 ml of C medium containing 0.1% (w/v) androsta-1,4-diene-3,17-dione (ADD), testosterone, epiandrosterone, dehydroandrosterone, cholic acid, or lithocholic acid. Cultures were incubated at 30°C, and growth was monitored every 3 h for 12 h. Viable cell numbers were determined by colony-forming unit (CFU) counts on LB agar plates after approximately 15 h of incubation.

### RNA extraction and Northern analysis

Total RNA was extracted from *M. resinovorans* NBRC 106553 cells cultured for 8 h in LB medium supplemented with testosterone, ADD, cholic acid, biphenyl, 3-hydroxyphenylpropionic acid (3HPP), or succinate as a negative control. Northern analysis was performed as described previously (16).

### Genome sequencing, assembly, and annotation

Genomic DNA of *C. testosteroni* TA441 was subjected to PacBio long-read sequencing using the SMRTbell prep kit 3.0 on a PacBio Revio system. Sequence reads were filtered to remove reads shorter than 1,000 bp using SeqKit v0.15.0. Genome assembly was performed using hifiasm v0.15.5-r350. Gene prediction and functional annotation were performed using DFAST v1.3.6. The completeness of the genome assembly and gene annotation was evaluated using BUSCO v4.1.2 with the betaproteobacteria_odb10 dataset. The resulting genome sequence was deposited in the DDBJ under accession number AP046141.

### Comparative genome analysis

Comparative analyses of steroid degradation gene clusters were performed using BLAST, Mauve v2.4.0, and CAGECAT. For each BLAST search, homologous sequences were identified using selected steroid degradation genes or proteins as query sequences. The query sequences and search targets used for each analysis are summarized in Table 10.

For each species, one strain showing the highest similarity to the query sequence was generally selected for further analysis. Multiple strains were selected for species of particular interest or when additional comparison was considered necessary. These included multiple *C. testosteroni* strains for detailed comparison with TA441, multiple *P. stutzeri* strains including strain Chol1 because of uncertainty in its sequence data, and multiple strains related to Chol1 because its steroid degradation gene cluster was located on different contigs. Additional strains were examined when an unusual gene cluster organization was observed to determine whether the feature was specific to an individual strain or conserved within the species.

For comparative genome analysis, a genomic region of approximately 200 kb was examined for each selected strain. The region extended approximately 100 kb upstream and downstream from the beginning of the genomic sequence showing homology to the BLAST query sequence unless otherwise indicated. The resulting genomic regions were compared using CAGECAT based on the presence, order, and sequence similarity of genes. When the relevant region was located at the end of a contig, additional contigs were examined when necessary to determine the organization of the corresponding gene cluster.

Genome-scale synteny among selected *Comamonas* strains was examined using Mauve v2.4.0.

### Phylogenetic analysis

Phylogenetic analyses were performed using 16S rRNA, gyrB, selected steroid degradation genes, and aromatic compound degradation genes. Sequences were obtained by BLAST searches against available genome sequences. Phylogenetic trees were generated from the resulting BLAST sequence sets using the Tree View function of BLAST, unless otherwise indicated.

For the phylogenetic analysis of Δ1-dehydrogenases, amino acid sequences identified in the preceding comparative analyses were collected, generally using one representative strain per species. A phylogenetic tree was constructed using MEGA Tree 12.1.1.

## Data Availability

The authors affirm that materials and data reasonably requested by others will be made available from a publicly accessible collection or provided in a timely manner at reasonable cost and in limited quantities to members of the scientific community for noncommercial purposes.

## REFERENCES

1. Dodson RM, Muir RD. 1961. Microbiological transformations. IV. The microbiological aromatization of steroids. J Am Chem Soc 83:4627–4631.

2. Gibson DT, Wang KC, Sih CJ, Whitlock H, Jr. 1966. Mechanisms of steroid oxidation by microorganisms. IX. On the mechanism of ring A cleavage in the degradation of 9,10-seco steroids by microorganisms. J Biol Chem 241:551–559.

3. Sih CJ, Lee SS, Tsong YK, Wang KC. 1966. Mechanisms of steroid oxidation by microorganisms. VIII. 3,4-Dihydroxy-9,10-secoandrosta-1,3,5(10)-triene-9,17-dione, an intermediate in the microbiological degradation of ring A of androst-4-ene-3,17-dione. J Biol Chem 241:540–550.

4. Coulter AW, Talalay P. 1968. Studies on the microbiological degradation of steroid ring A. J Biol Chem 243:3238–3247.

5. Oppermann UC, Maser E. 1996. Characterization of a 3 α -hydroxysteroid dehydrogenase/carbonyl reductase from the gram-negative bacterium *Comamonas testosteroni*. Eur J Biochem 241:744–749.

6. Mobus E, Maser E. 1999. Cloning and sequencing of a new *Comamonas testosteroni* gene encoding 3 alpha-hydroxysteroid dehydrogenase/carbonyl reductase. Adv Exp Med Biol 463:395–402.

7. Maser E, Xiong G, Grimm C, Ficner R, Reuter K. 2001. 3alpha-Hydroxysteroid dehydrogenase/carbonyl reductase from *Comamonas testosteroni*: biological significance, three-dimensional structure and gene regulation. Chem Biol Interact 130-132:707–722.

8. Genti-Raimondi S, Tolmasky ME, Patrito LC, Flury A, Actis LA. 1991. Molecular cloning and expression of the b-hydroxysteroid dehydrogenase gene from *Pseudomonas testosteroni*. Gene 105:43–49.

9. Itagaki E, Wakabayashi T, Hatta T. 1990. Purification and characterization of 3-ketosteroid-Δ1-dehydrogenase from *Nocardia corallina*. Biochim Biophys Acta 1038:60–67.

10. Morii S, Fujii C, Miyoshi T, Iwami M, Itagaki E. 1998. 3-Ketosteroid-Δ1-dehydrogenase of *Rhodococcus rhodochrous*: sequencing of the genomic DNA and hyperexpression, purification, and characterization of the recombinant enzyme. J Biochem (Tokyo) 124:1026–1032.

11. Cho HS, Ha NC, Choi G, Kim HJ, Lee D, Oh KS, Kim KS, Lee W, Choi KY, Oh BH. 1999. Crystal structure of Δ (5)-3-ketosteroid isomerase from *Pseudomonas testosteroni* in complex with equilenin settles the correct hydrogen bonding scheme for transition state stabilization. J Biol Chem 274:32863–32868.

12. Kim SW, Kim CY, Benisek WF, Choi KY. 1994. Cloning, nucleotide sequence, and overexpression of the gene coding for Δ 5-3-ketosteroid isomerase from *Pseudomonas putida* biotype B. J Bacteriol 176:6672–6676.

13. Molnar I, Choi KP, Yamashita M, Murooka Y. 1995. Molecular cloning, expression in *Streptomyces lividans*, and analysis of a gene cluster from *Arthrobacter simplex* encoding 3-ketosteroid-Δ1-dehydrogenase, 3-ketosteroid-Δ5-isomerase and a hypothetical regulatory protein. Mol Microbiol 15:895–905.

14. van der Geize R, Hessels GI, van Gerwen R, van der Meijden P, Dijkhuizen L. 2002. Molecular and functional characterization of *kshA* and *kshB*, encoding two components of 3-ketosteroid 9alpha-hydroxylase, a class IA monooxygenase, in *Rhodococcus erythropolis* strain SQ1. Mol Microbiol 45:1007–1018.

15. Horinouchi M, Yamamoto T, Taguchi K, Arai H, Kudo T. 2001. *Meta*-cleavage enzyme gene *tesB* is necessary for testosterone degradation in *Comamonas testosteroni* TA441. Microbiology 147:3367–3375.

16. Horinouchi M, Hayashi T, Yamamoto T, Kudo T. 2003. A new bacterial steroid degradation gene cluster which consists of aromatic compound degradation genes for seco-steroids and 3-ketosteroid dehydrogenase genes in *Comamonas testosteroni* TA441. Appl Environ Microbiol 69:4421–4430.

17. Horinouchi M, Hayashi T, Koshino H, Yamamoto T, Kudo T. 2003. Gene encoding the hydrolase for the product of the *meta*-cleavage reaction in testosterone degradation by *Comamonas testosteroni*. Appl Environ Microbiol 69:2139–2152.

18. Horinouchi M, Hayashi T, Kudo T. 2004. The genes encoding the hydroxylase of 3-hydroxy-9,10-secoandrosta-1,3,5(10)-triene-9,17-dione in steroid degradation in *Comamonas testosteroni* TA441. J Steroid Biochem Mol Biol 92:143–154.

19. Horinouchi M, Kurita T, Yamamoto T, Hatori E, Hayashi T, Kudo T. 2004. Steroid degradation gene cluster of *Comamonas testosteroni* consisting of 18 putative genes from *meta*-cleavage enzyme gene *tesB* to regulator gene *tesR*. Biochem Biophys Res Commun 324:597–604.

20. Horinouchi M, Hayashi T, Koshino H, Kurita T, Kudo T. 2005. Identification of 9,17-dioxo-1,2,3,4,10,19-hexanorandrostan-5-oic acid, 4-hydroxy-2-oxohexanoic acid, and 2-hydroxyhexa-2,4-dienoic acid and related enzymes involved in testosterone degradation in *Comamonas testosteroni* TA441. Appl Environ Microbiol 71:5275–5281.

21. Horinouchi M, Hayashi T, Koshino H, Kudo T. 2006. ORF18-disrupted mutant of *Comamonas testosteroni* TA441 accumulates significant amounts of 9,17-dioxo-1,2,3,4,10,19-hexanorandrostan-5-oic acid and its derivatives after incubation with steroids. J Steroid Biochem Mol Biol 101:78-84.

22. Horinouchi M, Hayashi T, Kudo T. 2012. Steroid degradation in *Comamonas testosteroni*. J Steroid Biochem Mol Biol 129:4–14.

23. Van der Geize R, Yam K, Heuser T, Wilbrink MH, Hara H, Anderton MC, Sim E, Dijkhuizen L, Davies JE, Mohn WW, Eltis LD. 2007. A gene cluster encoding cholesterol catabolism in a soil actinomycete provides insight into *Mycobacterium tuberculosis* survival in macrophages. Proc Natl Acad Sci U S A 104:1947–1952.

24. Yam KC, D’Angelo I, Kalscheuer R, Zhu H, Wang JX, Snieckus V, Ly LH, Converse PJ, Jacobs WR, Jr., Strynadka N, Eltis LD. 2009. Studies of a ring-cleaving dioxygenase illuminate the role of cholesterol metabolism in the pathogenesis of *Mycobacterium tuberculosis*. PLoS Pathog 5:e1000344.

25. Horinouchi M, Hayashi T, Koshino H, Malon M, Hirota H, Kudo T. 2014. Identification of 9alpha-Hydroxy-17-Oxo-1,2,3,4,10,19-Hexanorandrostan-5-Oic Acid in Steroid Degradation by *Comamonas testosteroni* TA441 and Its Conversion to the Corresponding 6-En-5-Oyl Coenzyme A (CoA) Involving Open Reading Frame 28 (ORF28)- and ORF30-Encoded Acyl-CoA Dehydrogenases. J Bacteriol 196:3598–608.

26. Horinouchi M, Hayashi T, Koshino H, Malon M, Hirota H, Kudo T. 2014. Identification of 9alpha-hydroxy-17-oxo-1,2,3,4,10,19-hexanorandrost-6-en-5-oic acid and beta-oxidation products of the C-17 side chain in cholic acid degradation by *Comamonas testosteroni* TA441. J Steroid Biochem Mol Biol 143:306–322.

27. Horinouchi M, Koshino H, Malon M, Hirota H, Hayashi T. 2018. Steroid degradation in *Comamonas testosteroni* TA441: identification of metabolites and the genes involved in the reactions necessary before D-ring cleavage. Appl Environ Microbiol 84:e01324–18.

28. Horinouchi M, Koshino H, Malon M, Hirota H, Hayashi T. 2019. Seroid Degradation in *Comamonas testosteroni* TA441: Identification of the Entire ?-Oxidation Cycle of the Cleaved B Ring. Appl Environ Microbiol 85:e01204–19.

29. Horinouchi M, Malon M, Hirota H, Hayashi T. 2019. Identification of 4-methyl-5-oxo-octane-1,8-dioic acid and the derivatives as metabolites of steroidal C,D-ring degradation in *Comamonas testosteroni* TA441. J Steroid Biochem Mol Biol 185:277–286.

30. Horinouchi M, Koshino H, Malon M, Hirota H, Hayashi T. 2019. Identification of 9-oxo-1,2,3,4,5,6,10,19-octanor-13,17-secoandrost-8(14)-ene-7,17-dioic acid as a metabolite of steroid degradation in *Comamonas testosteroni* TA441 and the genes involved in the conversion. J Steroid Biochem Mol Biol 185:268–276.

31. Horinouchi M, Hayashi T. 2021. Identification of the Coenzyme A (CoA) Ester Intermediates and Genes Involved in the Cleavage and Degradation of the Steroidal C-Ring by *Comamonas testosteroni* TA441. Appl Environ Microbiol 87:e0110221.

32. Horinouchi M, Hayashi T. 2023. Identification of “missing links” in C- and D-ring cleavage of steroids by *Comamonas testosteroni* TA441. Appl Environ Microbiol 89:e0105023.

33. Horinouchi M, Hayashi T. 2023. Comprehensive summary of steroid metabolism in C*omamonas testosteroni* TA441: entire degradation process of basic four rings and removal of C12 hydroxyl group. Appl Environ Microbiol 89:e0014323.

34. Horinouchi M, Hayashi T. 2025. Comprehensive review of steroid metabolism in C*omamonas testosteroni* TA441 with insights from other aerobic steroid-degrading bacteria. Advances in Applied Microbiology 131:1–20.

35. Horinouchi M. 2026. Identification of the C9-hydrogenase for 9,17-dioxo-1,2,3,4,10,19-hexanorandrostan-5-oic acid (9,17-DOHNA) and the 7α-dehydratase essential for initiating β-oxidation of the B-, C-, and D-rings in steroid degradation by *Comamonas testosteroni* TA441. Appl Environ Microbiol 92:e0233125.

36. Crowe AM, Casabon I, Brown KL, Liu J, Lian J, Rogalski JC, Hurst TE, Snieckus V, Foster LJ, Eltis LD. 2017. Catabolism of the last two steroid rings in *Mycobacterium tuberculosis* and other bacteria. MBio 8:doi: 10.1128/mBio.00321-17.

37. Horinouchi M, Kurita T, Hayashi T, Kudo T. 2010. Steroid degradation genes in *Comamonas testosteroni* TA441: Isolation of genes encoding a Delta4(5)-isomerase and 3alpha- and 3beta-dehydrogenases and evidence for a 100 kb steroid degradation gene hot spot. J Steroid Biochem Mol Biol 122:253–263.

38. Horinouchi M. 2025. Identification of dehydrogenase, hydratase, and aldolase responsible for the propionyl residue removal in degradation of cholic acid C-17 side chain in *Comamonas testosteroni* TA441. Microbiol Spectr 13:e0030825.

39. Arai H, Akahira S, Ohishi T, Maeda M, Kudo T. 1998. Adaptation of *Comamonas testosteroni* TA441 to utilize phenol: organization and regulation of the genes involved in phenol degradation. Microbiology 144:2895–2903.

40. Arai H, Yamamoto T, Ohishi T, Shimizu T, Nakata T, Kudo T. 1999. Genetic organization and characteristics of the 3-(3-hydroxyphenyl)propionic acid degradation pathway of *Comamonas testosteroni* TA441. Microbiology 145:2813–2820.

41. Ma YF, Zhang Y, Zhang JY, Chen DW, Zhu Y, Zheng H, Wang SY, Jiang CY, Zhao GP, Liu SJ. 2009. The complete genome of *Comamonas testosteroni* reveals its genetic adaptations to changing environments. Appl Environ Microbiol 75:6812–9.

42. Weiss M, Kesberg AI, Labutti KM, Pitluck S, Bruce D, Hauser L, Copeland A, Woyke T, Lowry S, Lucas S, Land M, Goodwin L, Kjelleberg S, Cook AM, Buhmann M, Thomas T, Schleheck D. 2013. Permanent draft genome sequence of Comamonas testosteroni KF-1. Stand Genomic Sci 8:239–54.

43. Fukuda K, Hosoyama A, Tsuchikane K, Ohji S, Yamazoe A, Fujita N, Shintani M, Kimbara K. 2014. Complete Genome Sequence of Polychlorinated Biphenyl Degrader *Comamonas testosteroni* TK102 (NBRC 109938). Genome Announc 2:e00865–14.

44. Park EH, Kim YS, Cha CJ. 2022. *Comamonas fluminis* sp. nov., isolated from the Han River, Republic of Korea. Int J Syst Evol Microbiol 72.

45. Yin Y, Han J, Wu H, Lu Y, Bao X, Lu Z. 2024. *Comamonas resistens* sp. nov. and *Pseudomonas triclosanedens* sp. nov., two members of the phylum Pseudomonadota isolated from the wastewater treatment system of a pharmaceutical factory. Int J Syst Evol Microbiol 74.

46. Plesiat P, Grandguillot M, Harayama S, Vragar S, Michel-Briand Y. 1991. Cloning, sequencing, and expression of the *Pseudomonas testosteroni* gene encoding 3-oxosteroid Δ1-dehydrogenase. J Bacteriol 173:7219–7227.

47. Florin C, Kohler T, Grandguillot M, Plesiat P. 1996. *Comamonas testosteroni* 3-ketosteroid-Δ4(5α)-dehydrogenase: gene and protein characterization. J Bacteriol 178:3322–3330.

48. Skowasch D, Mobus E, Maser E. 2002. Identification of a novel *Comamonas testosteroni* gene encoding a steroid-inducible extradiol dioxygenase. Biochem Biophys Res Commun 294:560–566.

49. Pruneda-Paz JL, Linares M, Cabrera JE, Genti-Raimondi S. 2004. TeiR, a LuxR-type transcription factor required for testosterone degradation in *Comamonas testosteroni*. J Bacteriol 186:1430–1437.

50. Gong W, Kisiela M, Schilhabel MB, Xiong G, Maser E. 2012. Genome sequence of *Comamonas testosteroni* ATCC 11996, a representative strain involved in steroid degradation. J Bacteriol 194:1633–1634.

51. Narayan KD, Pandey SK, Das SK. 2010. Characterization of *Comamonas thiooxidans* sp. nov., and comparison of thiosulfate oxidation with *Comamonas testosteroni* and *Comamonas composti*. Curr Microbiol 61:248–253.

52. (NITE) NIoTaE. Discrimination of C. testosteroni-C. thiooxidans. https://www.nite.go.jp/nbrc/safety/mlsa.html

53. Azwani F, Suzuki K, Honjyo M, Tashiro Y, Futamata H. 2017. Draft Genome Sequence of *Comamonas testosteroni* R2, Consisting of Aromatic Compound Degradation Genes for Phenol Hydroxylase. Genome Announc 5.

54. van den Belt M, Gilchrist C, Booth TJ, Chooi YH, Medema MH, Alanjary M. 2023. CAGECAT: The CompArative GEne Cluster Analysis Toolbox for rapid search and visualisation of homologous gene clusters. BMC Bioinformatics 24:181.

55. Frederico TD, Cunha-Ferreira IC, Vizzotto CS, de Sousa JF, Portugal MM, Tótola MR, Krüger RH, Peixoto J. 2025. Genomic and taxonomic characterization of the *Comamonas* sp. nov., a bacterium isolated from Brazilian Cerrado soil. Braz J Microbiol 56:137–154.

56. Park KH, Yu Z, Dong K, Lee SS. 2021. *Comamonas suwonensis* sp. nov., isolated from stream water in the Republic of Korea. Int J Syst Evol Microbiol 71.

57. Kämpfer P, Busse HJ, Baars S, Wilharm G, Glaeser SP. 2018. *Comamonas aquatilis* sp. nov., isolated from a garden pond. Int J Syst Evol Microbiol 68:1210–1214.

58. Park Y, Kim B, Min J, Park W. 2025. *Comamonas halotolerans* sp. nov., isolated from the faecal sample of a zoo animal, *Naemorhedus caudatus*. Int J Syst Evol Microbiol 75.

59. Zhang J, Wang Y, Zhou S, Wu C, He J, Li F. 2013. *Comamonas guangdongensis* sp. nov., isolated from subterranean forest sediment, and emended description of the genus *Comamonas*. Int J Syst Evol Microbiol 63:809–814.

60. Kim D, Lee SS. 2014. *Comamonas faecalis* sp. nov., isolated from domestic pig feces. Curr Microbiol 69:102–107.

61. Wu L, Ma J. 2019. The Global Catalogue of Microorganisms (GCM) 10K type strain sequencing project: providing services to taxonomists for standard genome sequencing and annotation. Int J Syst Evol Microbiol 69:895–898.

62. Ryan MP, Sevjahova L, Gorman R, White S. 2022. The Emergence of the Genus Comamonas as Important Opportunistic Pathogens. Pathogens 11:1032.

63. Wang Y, Zhou Z, Zhang W, Guo J, Li N, Zhang Y, Gong D, Lyu Y. 2024. Metabolic mechanism of Cr(VI) pollution remediation by *Alicycliphilus denitrificans* Ylb10. Sci Total Environ 912:169135.

64. Moriuchi R, Dohra H, Kanesaki Y, Ogawa N. 2019. Complete Genome Sequence of 3-Chlorobenzoate-Degrading Bacterium. Front Microbiol 10:133.

65. Hyun DW, Bae JW. 2020. Diaphorobacter sp. HDW4A chromosome, complete genome (Direct Submission).

66. Hwang CY, Cho ES, Seo MJ. 2025. Pseudomonas sp. MBLB4123 whole genome sequence (Direct Submission).

67. Busquets A, Mulet M, Gomila M, García-Valdés E. 2021. *Pseudomonas lalucatii* sp. nov. isolated from Vallgornera, a karstic cave in Mallorca, Western Mediterranean. Syst Appl Microbiol 44:126205.

68. Varghese N, Submissions S. 2025. Oryzisolibacter propanilivorax strain EPL6, whole genome shotgun sequence (Direct Submission).

69. Park M, Song J, Nam GG, Cho JC. 2019. *Rhodoferax lacus* sp. nov., isolated from a large freshwater lake. Int J Syst Evol Microbiol 69:3135–3140.

70. Wang Z, Chang X, Yang X, Pan L, Dai J. 2014. Draft Genome Sequence of *Polaromonas glacialis* Strain R3-9, a Psychrotolerant Bacterium Isolated from Arctic Glacial Foreland. Genome Announc 2.

71. Lefler FW, Berthold DE, Barbosa M, Hu J, Taylor A, Moretto J, Chaffin JD, Raymond H, Laughinghouse HD. 2025. Metagenomes from cyanobacterial harmful algal blooms from lakes in Ohio (USA). Microbiol Resour Announc 14:e0040025.

72. Bay SK, Ni G, Lappan R, Leung PM, Wong WW, Ry Holland SI, Athukorala N, Knudsen KS, Fan Z, Kerou M, Jain S, Schmidt O, Eate V, Clarke DA, Jirapanjawat T, Tveit A, Featonby T, White S, White N, McGeoch MA, Singleton CM, Cook PLM, Chown SL, Greening C. 2025. Microbial aerotrophy enables continuous primary production in diverse cave ecosystems. Nat Commun 16:10295.

73. Goeker M, Huntemann M, Clum A, Pillay M, Palaniappan K, Varghese N, Mikhailova N, Stamatis D, Reddy T, Daum C, Shapiro N, Ivanova N, Kyrpides N, Woyke T. 2019. Simplicispira metamorpha strain DSM 1837 Ga0310511_136, whole genome shotgun sequence (Direct Submission).

74. Wang S. 2025. Phenotypic and genotypic characterization of Roseateles flocculans sp. nov., isolated from Municipal sewage treatment plant, and Comparative Genomics of the Genus Roseateles (unpublished), whole genome shotgun sequence (Direct Submission).

75. Schneider D, Zühlke D, Poehlein A, Riedel K, Daniel R. 2021. Metagenome-Assembled Genome Sequences from Different Wastewater Treatment Stages in Germany. Microbiol Resour Announc 10:e0050421.

76. Mukherjee S, Seshadri R, Varghese NJ, Eloe-Fadrosh EA, Meier-Kolthoff JP, Göker M, Coates RC, Hadjithomas M, Pavlopoulos GA, Paez-Espino D, Yoshikuni Y, Visel A, Whitman WB, Garrity GM, Eisen JA, Hugenholtz P, Pati A, Ivanova NN, Woyke T, Klenk HP, Kyrpides NC. 2017. 1,003 reference genomes of bacterial and archaeal isolates expand coverage of the tree of life. Nat Biotechnol 35:676–683.

77. Narihara S, Chida S, Matsunaga N, Akimoto R, Akimoto M, Hagio A, Mori T, Nittami T, Sato M, Mun S, Kang H, Back JH, Takeda M. 2024. Taxonomic characterization of *Sphaerotilus microaerophilus* sp. nov., a sheath-forming microaerophilic bacterium of activated sludge origin. Arch Microbiol 206:252.

78. Beals DG, Carper DL, Hochanadel LH, Jawdy SS, Klingeman DM, Piatkowski BT, Weston DJ, Doktycz MJ, Pelletier DA. 2026. Genomic signatures in *Variovorax* enabling colonization of the Populus endosphere. mSystems 11:e0160525.

79. Ma B, Lu C, Wang Y, Yu J, Zhao K, Xue R, Ren H, Lv X, Pan R, Zhang J, Zhu Y, Xu J. 2023. A genomic catalogue of soil microbiomes boosts mining of biodiversity and genetic resources. Nat Commun 14:7318.

80. Holert J, Alam I, Larsen M, Antunes A, Bajic VB, Stingl U, Philipp B. 2014. Genome Sequence of *Pseudomonas* sp. Strain Chol1, a Model Organism for the Degradation of Bile Salts and Other Steroid Compounds. Genome Announc 1:e00014–12.

81. Holert J, Yucel O, Jagmann N, Prestel A, Moller HM, Philipp B. 2016. Identification of bypass reactions leading to the formation of one central steroid degradation intermediate in metabolism of different bile salts in *Pseudomonas* sp. strain Chol1. Environ Microbiol 18:3373–3389.

82. Holert J, Yucel O, Suvekbala V, Kulic Z, Moller H, Philipp B. 2014. Evidence of distinct pathways for bacterial degradation of the steroid compound cholate suggests the potential for metabolic interactions by interspecies cross-feeding. Environ Microbiol 16:1424–1440.

83. Holert J, Jagmann N, Philipp B. 2013. The essential function of genes for a hydratase and an aldehyde dehydrogenase for growth of *Pseudomonas* sp. strain Chol1 with the steroid compound cholate indicates an aldolytic reaction step for deacetylation of the side chain. J Bacteriol 195:3371–3380.

84. Holert J, Kulić Ž, Yücel O, Suvekbala V, Suter MJ, Möller HM, Philipp B. 2013. Degradation of the acyl side chain of the steroid compound cholate in *Pseudomonas* sp. strain Chol1 proceeds via an aldehyde intermediate. J Bacteriol 195:585–595.

85. Birkenmaier A, Moller HM, Philipp B. 2011. Identification of a thiolase gene essential for beta-oxidation of the acyl side chain of the steroid compound cholate in *Pseudomonas* sp. strain Chol1. FEMS Microbiol Lett 318:123–130.

86. Sanford RA, Chee-Sanford J, Orellana L, Konstantinidis K. 2017. Stutzerimonas degradans strain DCP-Ps1 NODE_22_length_297821_cov_80.631668, whole genome shotgun sequence (Direct Submission).

87. McFarland AG, Bertucci HK, Littman E, Shen J, Huttenhower C, Hartmann EM. 2021. Triclosan Tolerance Is Driven by a Conserved Mechanism in Diverse *Pseudomonas* Species. Appl Environ Microbiol 87:e02924–20.

88. He W, Gao M, Lv L, Wang J, Cai Z, Bai Y, Gao X, Gao G, Pu W, Jiao Y, Wan M, Song Q, Chen S, Liu JH. 2023. Persistence and molecular epidemiology of *bla*NDM-positive Gram negative bacteria in three broiler farms: A longitudinal study (2015–2021). J Hazard Mater 446:130725.

89. Varghese N, Submissions S. 2016. Pseudomonas oryzae strain KCTC 32247 genome assembly, chromosome: I (Direct Submission).

90. Gilroy R, Ravi A, Getino M, Pursley I, Horton DL, Alikhan NF, Baker D, Gharbi K, Hall N, Watson M, Adriaenssens EM, Foster-Nyarko E, Jarju S, Secka A, Antonio M, Oren A, Chaudhuri RR, La Ragione R, Hildebrand F, Pallen MJ. 2021. Extensive microbial diversity within the chicken gut microbiome revealed by metagenomics and culture. PeerJ 9:e10941.

91. Hosoyama A, Noguchi M, Numata M, Tsuchikane K, Hirakata S, Uohara A, Kitahashi Y, Ohji S, Ichikawa N, Kimura A, Yamazoe A, Fujita N. 2023. Pseudomonas jinjuensis NBRC 103047, whole genome shotgun sequencing project (Direct Submission).

92. Girard L, Lood C, Höfte M, Vandamme P, Rokni-Zadeh H, van Noort V, Lavigne R, De Mot R. 2021. The Ever-Expanding *Pseudomonas* Genus: Description of 43 New Species and Partition of the *Pseudom*onas *putida* Group. Microorganisms 9:1766.

93. Qiao A, Zhou H. 2023. Pseudomonas sp. AN-1 chromosome, complete genome (direct submission).

94. Yang G, Han L, Wen J, Zhou S. 2013. *Pseudomonas guangdongensis* sp. nov., isolated from an electroactive biofilm, and emended description of the genus *Pseudomonas Migula* 1894. Int J Syst Evol Microbiol 63:4599–4605.

95. Sato SI, Ouchiyama N, Kimura T, Nojiri H, Yamane H, Omori T. 1997. Cloning of genes involved in carbazole degradation of *Pseudomonas* sp. strain CA10: nucleotide sequences of genes and characterization of *meta*-cleavage enzymes and hydrolase. J Bacteriol 179:4841–4849.

96. Canavera G, Bellotti G, Tiwari H, Frioni T, Puglisi E. 2026. Drought-tolerant rhizobacterial consortia enhance grapevine growth and tolerance to water deficit. 17- 2026.

97. Piazza A, Casalini LC, Pacini VA, Sanguinetti G, Ottado J, Gottig N. 2019. Environmental Bacteria Involved in Manganese(II) Oxidation and Removal From Groundwater. Front Microbiol 10 - 2019.

98. Tsang HL, Huang JL, Lin YH, Huang KF, Lu PL, Lin GH, Khine AA, Hu A, Chen HP. 2016. Borneol Dehydrogenase from *Pseudomonas* sp. Strain TCU-HL1 Catalyzes the Oxidation of (+)-Borneol and Its Isomers to Camphor. Appl Environ Microbiol 82:6378–6385.

99. Wong AC, Lai GK, Griffin SD, Leung FC. 2021. Complete genome sequence of Pseudomonas lalkuanensis ACYW.190, a toluene-degrading strain isolated from garden soil in Hong Kon (unpublished). complete genome (Direct Submission)

100. Cardinali-Rezende J, Alexandrino PM, Nahat RA, Sant’Ana DP, Silva LF, Gomez JG, Taciro MK. 2015. Draft Genome Sequence of *Pseudomonas* sp. Strain LFM046, a Producer of Medium-Chain-Length Polyhydroxyalkanoate. Genome Announc 3.

101. Ichikawa N, Sato H, Tonouchi N. 2023. Pseudomonas resinovorans NBRC 113413, whole genome shotgun sequencing project (Direct Submission).

102. Hsiao TH, Chen PH, Wang PH, Brandon-Mong GJ, Li CW, Horinouchi M, Hayashi T, Ismail W, Meng M, Chen YL, Chiang YR. 2023. Harnessing microbial phylum-specific molecular markers for assessment of environmental estrogen degradation. Sci Total Environ 896:165152.

103. Hsiao TH, Lee TH, Chuang MR, Wang PH, Meng M, Horinouchi M, Hayashi T, Chen YL, Chiang YR. 2022. Identification of essential β-oxidation genes and corresponding metabolites for oestrogen degradation by actinobacteria. Microb Biotechnol 15:949–966.

104. Hsiao TH, Chen YL, Meng M, Chuang MR, Horinouchi M, Hayashi T, Wang PH, Chiang YR. 2021. Mechanistic and phylogenetic insights into actinobacteria-mediated oestrogen biodegradation in urban estuarine sediments. Microb Biotechnol 14:1212–1227.

105. Ibero J, Hernández-Fernández G, García JL, Galán B. 2026. Metabolism of bile salts in the estrogen degrading bacterium *Caenibius tardaugens*. Biodegradation 37:31.

106. Feller FM, Richtsmeier P, Wege M, Philipp B. 2021. Comparative Analysis of Bile-Salt Degradation in S*phingobium* sp. Strain Chol11 and *Pseudomonas stutzeri* Strain Chol1 Reveals Functional Diversity of Proteobacterial Steroid Degradation Enzymes and Suggests a Novel Pathway for Side Chain Degradation. Appl Environ Microbiol 87:e0145321.

107. Yucel O, Wibberg D, Philipp B, Kalinowski J. 2018. Genome Sequence of the Bile Salt-Degrading Bacterium *Novosphingobium* sp. Strain Chol11, a Model Organism for Bacterial Steroid Catabolism. Genome Announc 6.

108. Yucel O, Holert J, Ludwig KC, Thierbach S, Philipp B. 2017. A Novel Steroid-Coenzyme A Ligase from *Novosphingobium* sp. Strain Chol11 Is Essential for an Alternative Degradation Pathway for Bile Salts. Appl Environ Microbiol 84:e01492–17.

109. Zhang J, Gan W, Zhao R, Yu K, Lei H, Li R, Li X, Li B. 2020. Chloramphenicol biodegradation by enriched bacterial consortia and isolated strain *Sphingomonas* sp. CL5.1: The reconstruction of a novel biodegradation pathway. Water Res 187:116397.

110. Ibero J, Galán B, Díaz E, García J. 2019. Testosterone Degradative Pathway of *Novosphingobium tardaugens*. Genes 10:pii: E871. doi: 10.3390/genes10110871.

111. Shah M, Bornemann TLV, Nuy JK, Hahn MW, Probst AJ, Beisser D, Boenigk J. 2024. Genome-resolved metagenomics reveals the effect of nutrient availability on bacterial genomic properties across 44 European freshwater lakes. Environ Microbiol 26:e16634.

112. Lin SY, Liu YC, Hameed A, Hsu YH, Huang HI, Lai WA, Young CC. 2016. *Azospirillum agricola* sp. nov., a nitrogen-fixing species isolated from cultivated soil. Int J Syst Evol Microbiol 66:1453–1458.

113. Benaud N, Edwards RJ, Amos TG, D’Agostino PM, Gutiérrez-Chávez C, Montgomery K, Nicetic I, Ferrari BC. 2021. Antarctic desert soil bacteria exhibit high novel natural product potential, evaluated through long-read genome sequencing and comparative genomics. Environ Microbiol 23:3646–3664.

114. Soltysiak MPM, Jalihal AP, Christophersen CE, Ory ALH, Lee AD, Boulton J, Springer M. 2025. Expansion and revision of the genus *Xanthobacter* and proposal of *Roseixanthobacter* gen. nov. Int J Syst Evol Microbiol 75.

115. Zilli JE, Schwab S, Dos Santos Ferreira N, Reis VM, de Oliveira Junior AF, Simões-Araujo JL, de Barros Soares LH, Dos Santos Dourado F, Bach E, Roesch LFW, Rossi CN, de Oliveira Lima de Souza KM, Alves BJR, Silva AL, Baldani JI. 2025. Diversity of grass-associated *Nitrospirillum* and proposal of six novel species. Braz J Microbiol 56:2827–2843.

116. Bai Y, Müller DB, Srinivas G, Garrido-Oter R, Potthoff E, Rott M, Dombrowski N, Münch PC, Spaepen S, Remus-Emsermann M, Hüttel B, McHardy AC, Vorholt JA, Schulze-Lefert P. 2015. Functional overlap of the Arabidopsis leaf and root microbiota. Nature 528:364–369.

117. Nakayasu M, Takamatsu K, Kanai K, Masuda S, Yamazaki S, Aoki Y, Shibata A, Suda W, Shirasu K, Yazaki K, Sugiyama A. 2023. Tomato root-associated *Sphingobium* harbors genes for catabolizing toxic steroidal glycoalkaloids. mBio 14:e0059923.

118. Sih CJ, Tai HH, Tsong YY, Lee SS, Coombe RG. 1968. Mechanisms of steroid oxidation by microorganisms. XIV. Pathway of cholesterol side-chain degradation. Biochemistry 7:808–818.

119. Tai HH, Sih CJ. 1970. 3,4-dihydroxy-9,10-secoandrosta-1,3,5(10)-triene-9,17-dione 4,-5-dioxygenase from *Nocardia restrictus*. I. Isolation of the enzyme and study of its physical and chemical properties. J Biol Chem 245:5062–5071.

120. van Der Geize R, Hessels GI, van Gerwen R, Vrijbloed JW, van Der Meijden P, Dijkhuizen L. 2000. Targeted disruption of the *kstD* gene encoding a 3-ketosteroid- Δ (1)-dehydrogenase isoenzyme of *Rhodococcus erythropolis* strain SQ1. Appl Environ Microbiol 66:2029–2036.

121. Kuliopulos A, Shortle D, Talalay P. 1987. Isolation and sequencing of the gene encoding Δ5-3-ketosteroid isomerase of *Pseudomonas testosteroni*: overexpression of the protein. Proc Natl Acad Sci U S A 84:8893–8897.

122. Brzostek A, Sliwinski T, Rumijowska-Galewicz A, Korycka-Machala M, Dziadek J. 2005. Identification and targeted disruption of the gene encoding the main 3-ketosteroid dehydrogenase in *Mycobacterium smegmatis*. Microbiology 151:2393–2402.

123. Knol J, Bodewits K, Hessels GI, Dijkhuizen L, van der Geize R. 2008. 3-Keto-5alpha-steroid Delta(1)-dehydrogenase from Rhodococcus erythropolis SQ1 and its orthologue in Mycobacterium tuberculosis H37Rv are highly specific enzymes that function in cholesterol catabolism. Biochem J 410:339–346.

124. Sukhodolskaya GV, Nikolayeva VM, Khomutov SM, Donova MV. 2007. Steroid-1-dehydrogenase of *Mycobacterium* sp. VKM Ac-1817D strain producing 9alpha-hydroxy-androst-4-ene-3,17-dione from sitosterol. Appl Microbiol Biotechnol 74:867–873.

125. Fernández de las Heras L, van der Geize R, Drzyzga O, Perera J, María Navarro Llorens J. 2012. Molecular characterization of three 3-ketosteroid-Δ(1)-dehydrogenase isoenzymes of *Rhodococcus ruber* strain Chol-4. J Steroid Biochem Mol Biol 132:271–281.

126. Thomas ST, VanderVen BC, Sherman DR, Russell DG, Sampson NS. 2011. Pathway profiling in *Mycobacterium tuberculosis*: elucidation of cholesterol-derived catabolite and enzymes that catalyze its metabolism. J Biol Chem 286:43668–43678.

127. van der Geize R, Hessels GI, Nienhuis-Kuiper M, Dijkhuizen L. 2008. Characterization of a second *Rhodococcus erythropolis* SQ1 3-ketosteroid 9alpha-hydroxylase activity comprising a terminal oxygenase homologue, KshA2, active with oxygenase-reductase component KshB. Appl Environ Microbiol 74:7197–203.

128. Dresen C, Lin LY, D’Angelo I, Tocheva EI, Strynadka N, Eltis LD. 2010. A flavin-dependent monooxygenase from *Mycobacterium tuberculosis* involved in cholesterol catabolism. J Biol Chem 285:22264–75.

129. Thomas ST, Sampson NS. 2013. *Mycobacterium tuberculosis* utilizes a unique heterotetrameric structure for dehydrogenation of the cholesterol side chain. Biochemistry 52:2895–2904.

130. Wipperman MF, Yang M, Thomas ST, Sampson NS. 2013. Shrinking the FadE proteome of *Mycobacterium tuberculosis*: insights into cholesterol metabolism through identification of an α2β2 heterotetrameric acyl coenzyme A dehydrogenase family. J Bacteriol 195:4331–41.

131. Yang M, Guja KE, Thomas ST, Garcia-Diaz M, Sampson NS. 2014. A distinct MaoC-like enoyl-CoA hydratase architecture mediates cholesterol catabolism in *Mycobacterium tuberculosis*. ACS Chem Biol 9:2632–2645.

132. Yang M, Lu R, Guja KE, Wipperman MF, St Clair JR, Bonds AC, Garcia-Diaz M, Sampson NS. 2015. Unraveling Cholesterol Catabolism in *Mycobacterium tuberculosis*: ChsE4-ChsE5 alpha2beta2 Acyl-CoA Dehydrogenase Initiates beta-Oxidation of 3-Oxo-cholest-4-en-26-oyl CoA. ACS Infect Dis 1:110–125.

133. Yuan T, Yang M, Gehring K, Sampson NS. 2019. *Mycobacterium tuberculosis* Exploits a Heterohexameric Enoyl-CoA Hydratase Retro-Aldolase Complex for Cholesterol Catabolism. Biochemistry 58:4224–4235.

134. Sangal V, Jones AL, Goodfellow M, Sutcliffe IC, Hoskisson PA. 2014. Comparative genomic analyses reveal a lack of a substantial signature of host adaptation in *Rhodococcus equi* (’*Prescottella equi*’). Pathog Dis 71:352–356.

135. Funo K, Kitagawa W, Tanaka M, Sone T, Asano K, Kamagata Y. 2014. Draft Genome Sequence of *Tomitella biformata* AHU 1821T, Isolated from a Permafrost Ice Wedge in Alaska. Genome Announc 2.

136. Lee SD, Yang HL, Kim IS. 2025. *Antrihabitans auranticaus* sp. nov., Isolated from a Cave, and Emended Description of the Genus *Antrihabitans*. Curr Microbiol 83:108.

137. Cole ST, Barrell BG. 1998. Analysis of the genome of *Mycobacterium tuberculosis* H37Rv. Novartis Found Symp 217:160–172.

138. McLeod MP, Warren RL, Hsiao WW, Araki N, Myhre M, Fernandes C, Miyazawa D, Wong W, Lillquist AL, Wang D, Dosanjh M, Hara H, Petrescu A, Morin RD, Yang G, Stott JM, Schein JE, Shin H, Smailus D, Siddiqui AS, Marra MA, Jones SJ, Holt R, Brinkman FS, Miyauchi K, Fukuda M, Davies JE, Mohn WW, Eltis LD. 2006. The complete genome of *Rhodococcus* sp. RHA1 provides insights into a catabolic powerhouse. Proc Natl Acad Sci U S A 103:15582–7.

